# Leveraging Homologous Recombination Deficiency via the Repositioned Prodrug CB1954

**DOI:** 10.64898/2026.07.08.737246

**Authors:** James L. Elia, Jarvis Hill, Collin D. Heer, Siji Smolev, Abbey M. Sykes, Sofia R. Arbelaez, Karlie N. Lucas, Spenser S. Johnson, Ranjini K. Sundaram, Seth B. Herzon, Ranjit S. Bindra

## Abstract

Homologous recombination deficiency (HRD) is an actionable vulnerability found in a substantial fraction of human cancers, yet current HRD-directed therapies are limited by toxicity, incomplete responses, and acquired resistance. Many DNA-damaging agents were developed before DNA repair biomarkers were available, suggesting that abandoned agents may harbor previously unrecognized genotype-selective activity. Here, through a focused screen of DNA-damaging agents in isogenic homologous recombination-proficient and -deficient models, we identify CB1954, a decades-old nitrobenzamide aziridine prodrug, as highly selective for BRCA2-deficient tumor cells. CB1954 forms DNA interstrand crosslinks independent of HR status, but selectively induces DNA-damage signaling, apoptosis, and loss of clonogenic survival in HR-deficient cells. Targeted DDR CRISPR screening and isogenic validation define a distinct repair dependency for the Fanconi anemia and homologous recombination pathways, with limited dependence on mismatch repair or nucleotide excision repair. Genetic and pharmacologic perturbation of NQO2, the bioactivating enzyme for CB1954, reveals a bifurcated mechanism in which NQO2-dependent activation selectively contributes to HRD cytotoxicity, while aziridine-dependent lesions likely account for residual activity in HR-proficient cells. CB1954 exhibits favorable preclinical pharmacokinetic properties and genotype-dependent antitumor activity in BRCA2-deficient xenografts. These findings reposition CB1954 as a historically overlooked HRD-selective agent and demonstrate that biomarker-guided profiling of DNA-damaging agents can uncover new opportunities for precision oncology.

## Introduction

Homologous recombination deficiency (HRD) is a key actionable DNA repair vulnerability in human cancer. HR-deficient tumors are impaired in the accurate repair of replication-associated double-strand breaks and in the resolution of repair intermediates generated during interstrand crosslink (ICL) repair [1, 2]. This defect underlies the clinical activity of platinum-based chemotherapy [3] and poly(ADP-ribose) polymerase inhibitors in BRCA1/2-mutant and other HR-deficient cancers [4]. However, both therapeutic classes have important limitations. Platinum agents induce a heterogeneous spectrum of DNA lesions that engage multiple repair pathways and are limited by systemic toxicity [5], while PARP inhibitors indirectly exploit HRD through perturbation of DNA repair signaling and are limited by incomplete responses and acquired resistance [6, 7, 8]. These limitations highlight the need for additional HRD-directed agents with distinct mechanisms of DNA damage and repair dependencies.

Many DNA-damaging agents were discovered decades before tumor genotyping, synthetic lethality, and DNA repair biomarkers became central to oncology drug development. As a result, these compounds were generally evaluated as broadly cytotoxic agents rather than as genotype-directed therapies. This historical gap raises an important question: whether older DNA-reactive agents, previously abandoned or clinically deprioritized, might possess unrecognized selectivity when applied to tumors with defined DNA repair defects.

CB1954, also known as tretazicar, provides a compelling example of this possibility. CB1954 is a 2,4-dinitrobenzamide aziridine prodrug first described more than fifty years ago as a highly active agent in Walker carcinoma models [9]. Subsequent work attributed this activity to enzymatic nitroreduction, which generates DNA-reactive species capable of forming cytotoxic DNA adducts, including ICLs [10]. However, the extraordinary sensitivity of Walker tumors was later linked to unusually high nitroreductase activity in that model, while human tumors were not expected to achieve comparable tumor-selective activation [11, 12]. Clinical development therefore shifted toward enzyme-prodrug strategies designed to introduce bacterial nitroreductase into tumors [13, 14]. These approaches did not produce sufficient efficacy at tolerated doses, and CB1954 was largely abandoned as a systemic anticancer agent at the time.

Importantly, these historical studies did not consider the DNA-damage response (DDR) status of the tumor. In particular, they did not ask whether CB1954-induced lesions create a selective dependency on the Fanconi anemia (FA) and homologous recombination (HR) pathways, nor whether HR-deficient tumors might be uniquely sensitive to CB1954 at concentrations that are tolerated by HR-proficient cells. This distinction is critical because the therapeutic window for a DNA-damaging agent is determined not only by where and how efficiently the lesion is formed, but also by whether tumor cells retain the repair pathways required to survive that lesion [15, 16].

Here, we evaluate CB1954 in the context of HR deficiency. Through a focused screen of DNA-damaging agents in isogenic HR-proficient and -deficient models, we identify CB1954 as a highly selective agent against BRCA2-deficient cells. We show that CB1954 forms ICLs independent of HR status, but our data suggest that HR-deficient cells respond to these lesions with lethal DNA-damage signaling and apoptosis at markedly lower concentrations than HR-proficient cells. Targeted CRISPR screening and isogenic validation define a repair dependency centered on the Fanconi anemia and HR pathways, without strong involvement of mismatch repair or nucleotide excision repair. Finally, genetic and pharmacologic studies of dihydronicotinamdide riboside (NRH) quinone oxidoreductase 2 (NQO2) reveal a bifurcated mechanism in which NQO2-dependent bioactivation selectively contributes to HRD cytotoxicity, while distinct aziridine-dependent lesions likely account for toxicity in HR-proficient cells. Together, these findings reposition CB1954 as a historically overlooked HRD-selective agent and suggest that biomarker-guided profiling of DNA-damaging agents may uncover new opportunities for precision oncology.

## Results

### Selective cytotoxicity of CB1954 in HRD cell lines

To determine if DNA crosslinkers with diverse structures (Figure S1A) could yield enhanced selectivity for HR-deficient tumors, we screened a panel of clinically relevant agents against BRCA2-isogenic cell lines. We utilized 5-day growth delay assays in DLD1 and HCT116 pairs (± BRCA2), alongside a doxycycline-inducible U2OS shBRCA2 model and U2OS ± RAD51D. We define the therapeutic index (TI) as the ratio of the IC_50_ in DNA repair-proficient cells to the IC_50_ in repair-deficient cells. While standard crosslinkers exhibited TIs ranging from 3-to 50-fold, CB1954 emerged as an outlier, yielding a TI of approximately 1000-fold across the BRCA2-deficient pairs (Figure 1A). The IC_50_ for HR-deficient cells clustered around 50 nM, whereas HR-proficient cell IC_50_ remained near 50 *µ*M across several isogenic pairs (Figure 1B). This corroborates *in vitro* phenotypic susceptibility data first disclosed in a thesis from the Masson Laboratory [17]. To the best of our knowledge, these findings have not been disclosed in a peer-reviewed publication.

**Figure 1.**
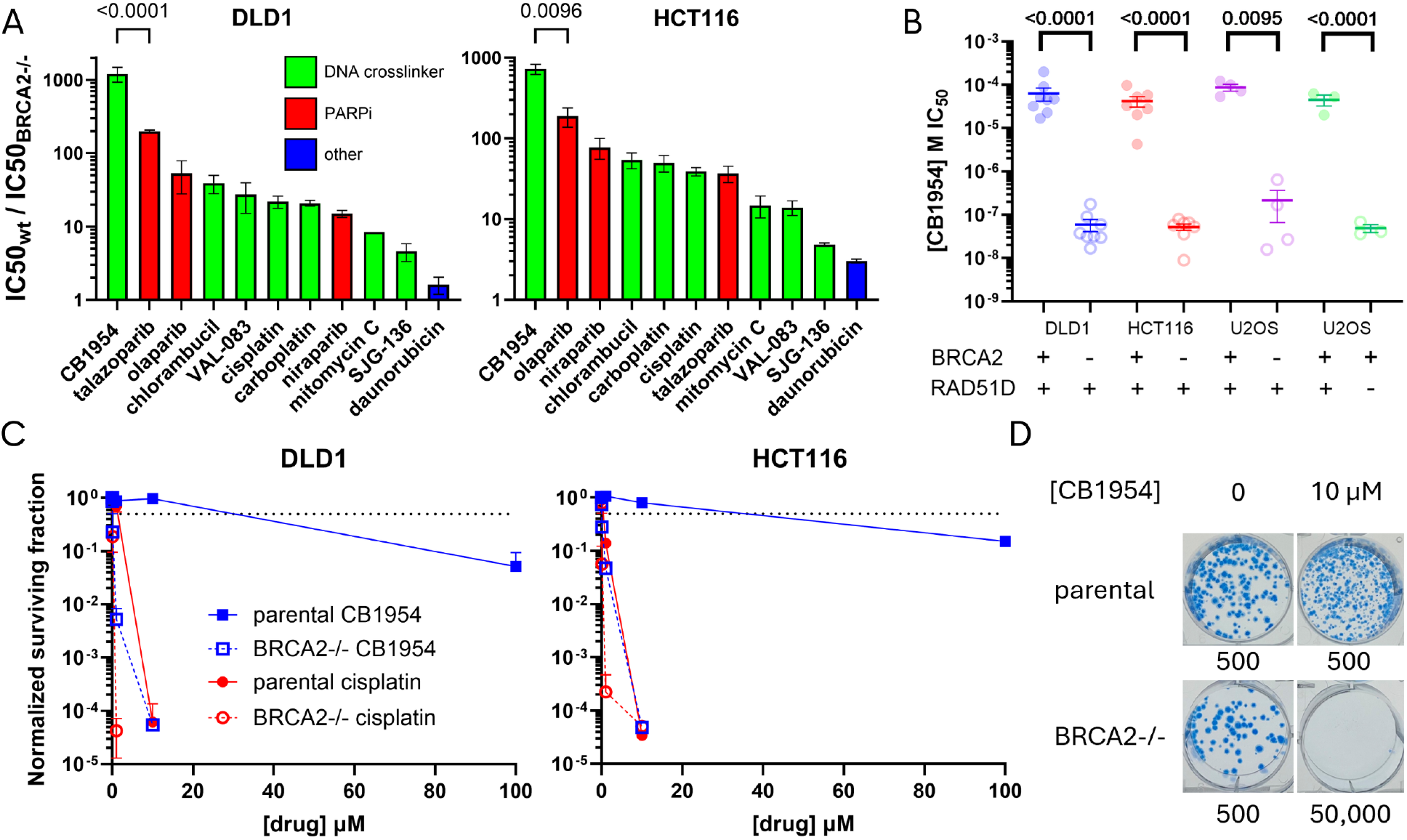
CB1954 is a uniquely selective DNA crosslinker. **A)** 5-day growth delay assay IC_50_ ratios with BRCA2 isogenic DLD1 and HCT116 using clinically relevant DNA crosslinkers (green), PARPi (red), and daunorubicin is a non-selective control (blue). Mean of biological replicates ± SEM as error bars and lognormal Welch’s t test for statistical significance. **B)** CB1954 growth delay assay IC_50_ across three BRCA2 isogenic models and one RAD51D isogenic model. Mean of biological replicates ± SEM as error bars. Lognormal Welch’s t test for statistical significance. **D)** 14-day clonogenic survival assays (CSAs) for CB1954 and cisplatin. Mean of biological replicates ± SEM as error bars. The dotted line marks 50% surviving fraction. **E)** HCT116 ± BRCA2 CSA representative wells after treatment with vehicle (0.1% DMSO) or 10 *µ*M CB1954. 500 and 50,000 refer to seeding density.

To validate this acute sensitivity in a long-term proliferation model, we performed clonogenic survival assays comparing CB1954 to the clinical standard, cisplatin. While no concentration of cisplatin could eradicate BRCA2-deficient colonies without inflicting severe, concurrent toxicity on the HR-proficient lines, treatment with 10 *µ*M CB1954 completely ablated colony formation in BRCA2-deficient cells, even at plating densities of 50,000 cells per well. Conversely, this same concentration had negligible impact on the cloning efficiency of the parental cell lines (Figures 1C, 1D).

### Genotype-agnostic ICL formation but selective apoptosis

To establish whether CB1954’s selectivity stems from differential initial lesion formation, we first confirmed its capacity to form ICLs *in vitro*, designed in concordance with the reported bioactivation mechanism [13] (Figure S1B). Incubation of linearized plasmid DNA with CB1954 and an NQO2 reaction mixture yielded dose-dependent ICL formation after 16 hours at biologically relevant concentrations (Figure 2A). We then quantified cellular ICL formation using the reverse comet assay (single-cell alkaline gel electrophoresis post-ionizing radiation), where the retention of high-molecular-weight DNA fragments in the comet head serves as a physical proxy for crosslinking [18] (Figure S2B). Using mitomycin C as a positive control due to its high proportion of ICL formation [19], we observed comparable, dose-dependent ICL formation in both BRCA2-proficient and BRCA2-deficient DLD1 cells (Figure 2B), a trend consistently mirrored in the HCT116 isogenic pair (Figure S3A).

**Figure 2.**
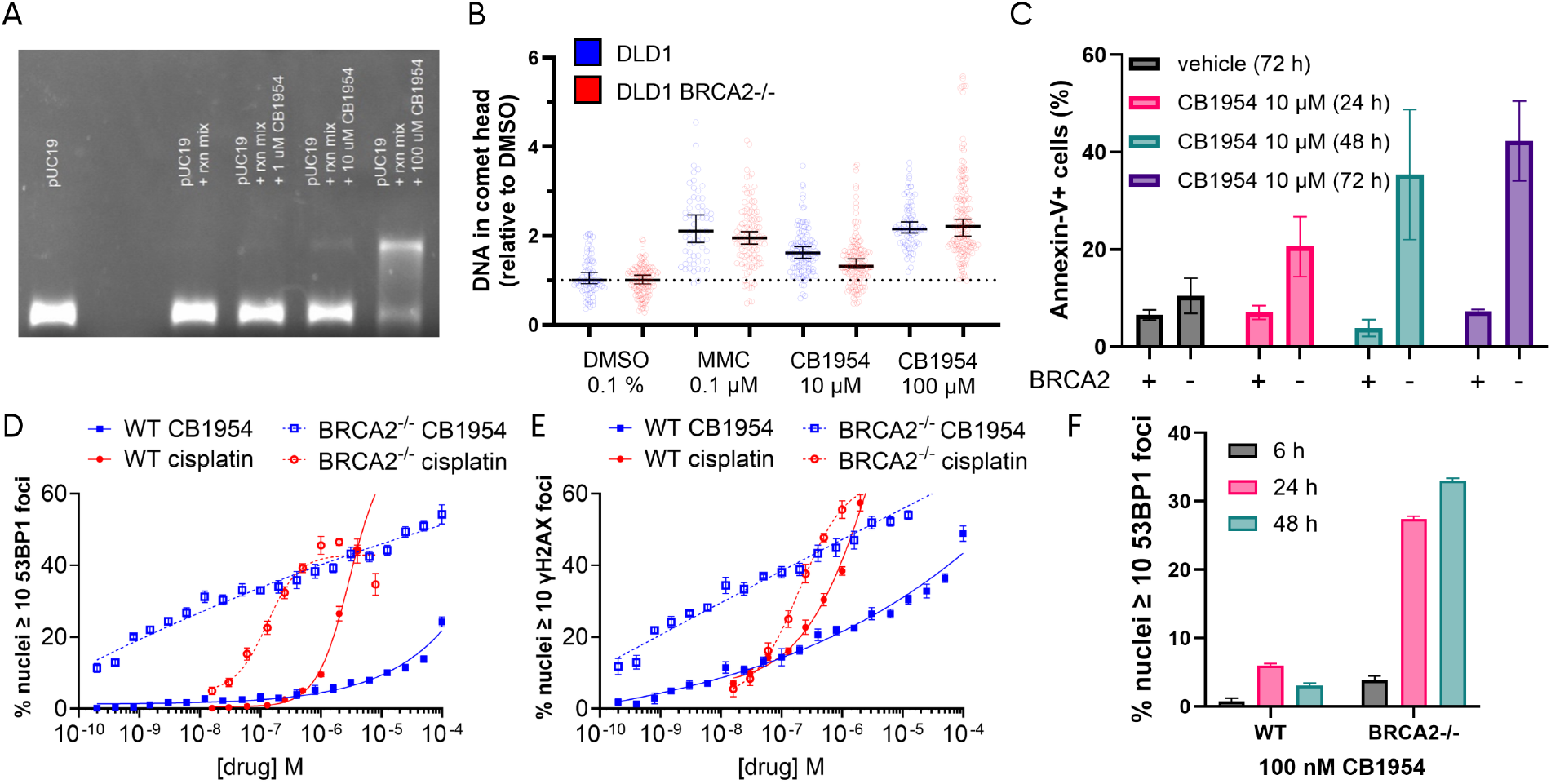
CB1954 is genotype-agnostic in ICL formation but selective in apoptosis and DDR. **A)** pDNA was treated with noted conditions for 16 h, then denatured into ssDNA and run on a gel. ICLs create dsDNA that runs more slowly. **B)** DNA in comet head is proportional to in cellulo ICLs in the reverse comet assay, normalized to vehicle within each cell line. Median of all nuclei with 95% confidence intervals as error bars. **C)** Annexin V flow cytometry signal in response to 10 *µ*M CB1954 at 24 h, 48 h, and 72 h. Mean of biological replicates with SEM as error bars. **D)** High-throughput immunofluorescence assays for 53BP1, a marker for DNA DSBs; cells with 10+ discrete foci in the nucleus are considered positive. Data shown are 48 h after treatment and have background (vehicle-only mean foci) subtracted. Mean of technical replicates with SD as error bars. **E)** High-throughput immunofluorescence assays for *γ*H2AX, a marker for general DNA damage; cells with 10+ discrete foci in the nucleus are considered positive. Data shown are 48 h after treatment and have background (vehicle-only mean foci) subtracted. Mean of technical replicates with SD as error bars. **F)** High-throughput immunofluorescence 6 h, 24 h, and 48 h time-course at 100 nM CB1954. Mean of technical replicates with SD as error bars.

Despite forming ICLs independent of HR status, CB1954 triggers a divergent cellular fate. Flow cytometric analysis of Annexin V revealed that 10 *µ*M CB1954 continuously induced apoptosis over 72 hours exclusively in BRCA2^−/−^ cells (Figure 2C), with the parental cells showing no meaningful apoptotic signal until exposed to 100 *µ*M (Figure S3B).

To determine if this bifurcated survival response originates at the level of DNA-damage signaling, we performed high-throughput fluorescence microscopy dose-response assays for *γ*H2AX (a marker of general DNA damage and replication stress) and 53BP1 (a marker of DSB recognition) [20, 21] (Figure S2A). Strikingly, sub-nanomolar concentrations of CB1954 induced abundant, dose-dependent DDR foci in BRCA2^−/−^ cells at 48 hours, with both markers tightly coupled (Figures 2D, 2E).

By contrast, the parental cells exhibited signaling tolerance and a distinct uncoupling of these markers. Among parental cells, <20% were *γ*H2AX+ until 1 *µ*M and <20% were 53BP1+ until 100 *µ*M, approximately two-fold their IC_50_ (Figures 2D, 2E). This signaling divergence led us to formulate a dual-lesion model of cytotoxicity: in HR-proficient cells, toxicity is primarily driven by abundant DNA monoadducts that induce general DDR signaling (*γ*H2AX) but do not collapse into DSBs (53BP1) until high concentrations. Conversely, in HR-deficient cells, the failure to process complex lesions like ICLs drives lethal DSB formation through replication fork collapse and excision by the Fanconi anemia/ICL repair pathway.

Time-course analysis at 100 nM further highlighted this discrepancy, revealing a compounding accumulation of 53BP1+ cells in the BRCA2^−/−^ population over 48 hours, compared to a transient, low-level response in the parental line that resolved by 48 hours (Figure 2F). By contrast, *γ*H2AX+ parental cells do increase through 48 hours after treatment with 100 nM CB1954 (Figure S3C). Along with full dose-response data for cisplatin, we included additional single-concentration controls, which demonstrate that foci-positive BRCA2^−/−^ cells greatly increase only in response to selective agents, like mitomycin C and olaparib, unlike non-HR-selective DNA-damaging agents, like daunorubicin or etoposide (Figures S3D, S3E).

### DDR CRISPR screen identifies key mediators of CB1954 repair

To systematically map the genetic networks required for the repair of CB1954-induced damage and to evaluate its divergence from classical clinical crosslinkers, we performed a targeted DDR CRISPR screen in the HR-proficient glioblastoma cell line LN229. Utilizing a custom library targeting 353 DDR genes [22], we compared the genetic dependencies of CB1954 against cisplatin (data from [22]) over four treatment cycles across 12 days. All screens maintained robust quality control metrics and >500x read-per-guide coverage (Figure S4A). Three concentrations of CB1954 across separate biological replicates yielded similar hit profiles (Figure S4B).

Concordant with its capacity to form ICLs, CB1954 shared fundamental negative selection hits with cisplatin across the Fanconi anemia pathway, with BRIP1 (FANCJ) emerging among strongest sensitizing knockouts for both agents (Figures 3A, 3B). Furthermore, TP53 was the sole shared hit for positive selection. Crucially, the screen illuminated distinct mechanistic departures: while loss of the mismatch repair (MMR) pathway conferred expected resistance to cisplatin [23], no MMR components emerged as resistance hits for CB1954 across any biological replicates (Figure 3C), with the exception of *EXO1* which also performs end resection during HR [24, 25]. Additionally, loss of the transcription-coupled nucleotide excision repair (TC-NER) sensor ERCC8 (CSA) increased sensitivity to cisplatin but not CB1954 (Figure S4C).

**Figure 3.**
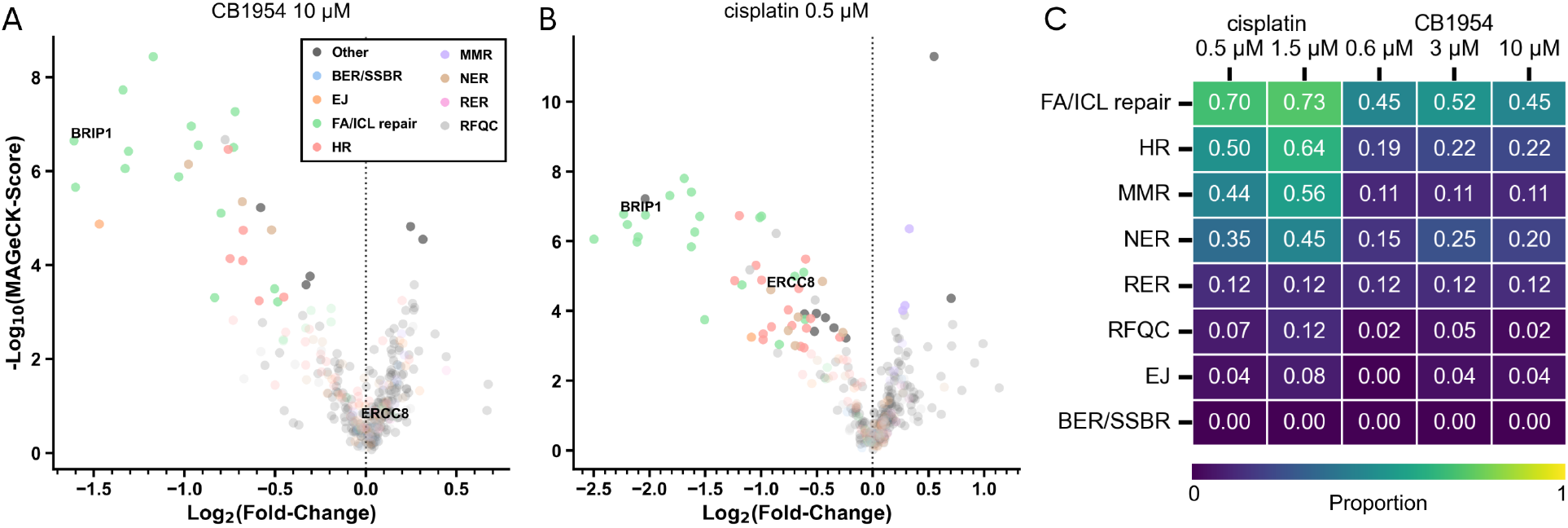
DDR CRISPR screen identifies key mediators of CB1954 repair. **A)** Comparison of LFC for CB1954- and cisplatin-treated LN229. **B)** IC_50_ ratios of U2OS Cas9 ± RAD51D with cisplatin, talazoparib, and CB1954. Mean of biological replicates ± SEM as error bars. **C)** Heatmap representing proportion of significant hits (FDR < 0.05) out of all genes with pathway designation; pathways are non-exclusive.

Considering BRIP1’s role in ICL repair and emergence as a top sensitizing hit, we sought to validate the finding and determine the breadth of selectivity with an isogenic model. Genetic loss of BRIP1 led to a 5-10-fold sensitization to CB1954 and cisplatin, with no effect on talazoparib sensitivity (Figure S4D).

Together, these data indicate that CB1954 repair is dependent on the FA and HR cascades, exhibiting potent chemosensitivity in specific downstream HR deficiencies. CB1954 has little reliance on MMR or NER, suggesting its DNA lesions are not good substrates for these pathways and thus require replication to be detected, where FA and HR play the primary role.

### NQO2 bioactivation dictates HR-deficient, but not HR-proficient, cytotoxicity

While NQO2 reportedly utilizes non-biogenic NRH as a co-substrate, the strongest correlation for CB1954 sensitivity across the Cancer Dependency Map (DepMap) is NQO2, its primary bioactivating enzyme (Figure 4A) [11, 12]. Immunoblotting confirmed that NQO2 protein expression is consistent across the DLD1 and HCT116 isogenic pairs (Figure 4B), aligning with the genotype-agnostic ICL formation observed previously (Figure 2B). To determine the precise contribution of NQO2 to BRCA2 selectivity, we utilized both pharmacologic and genetic interventions.

**Figure 4.**
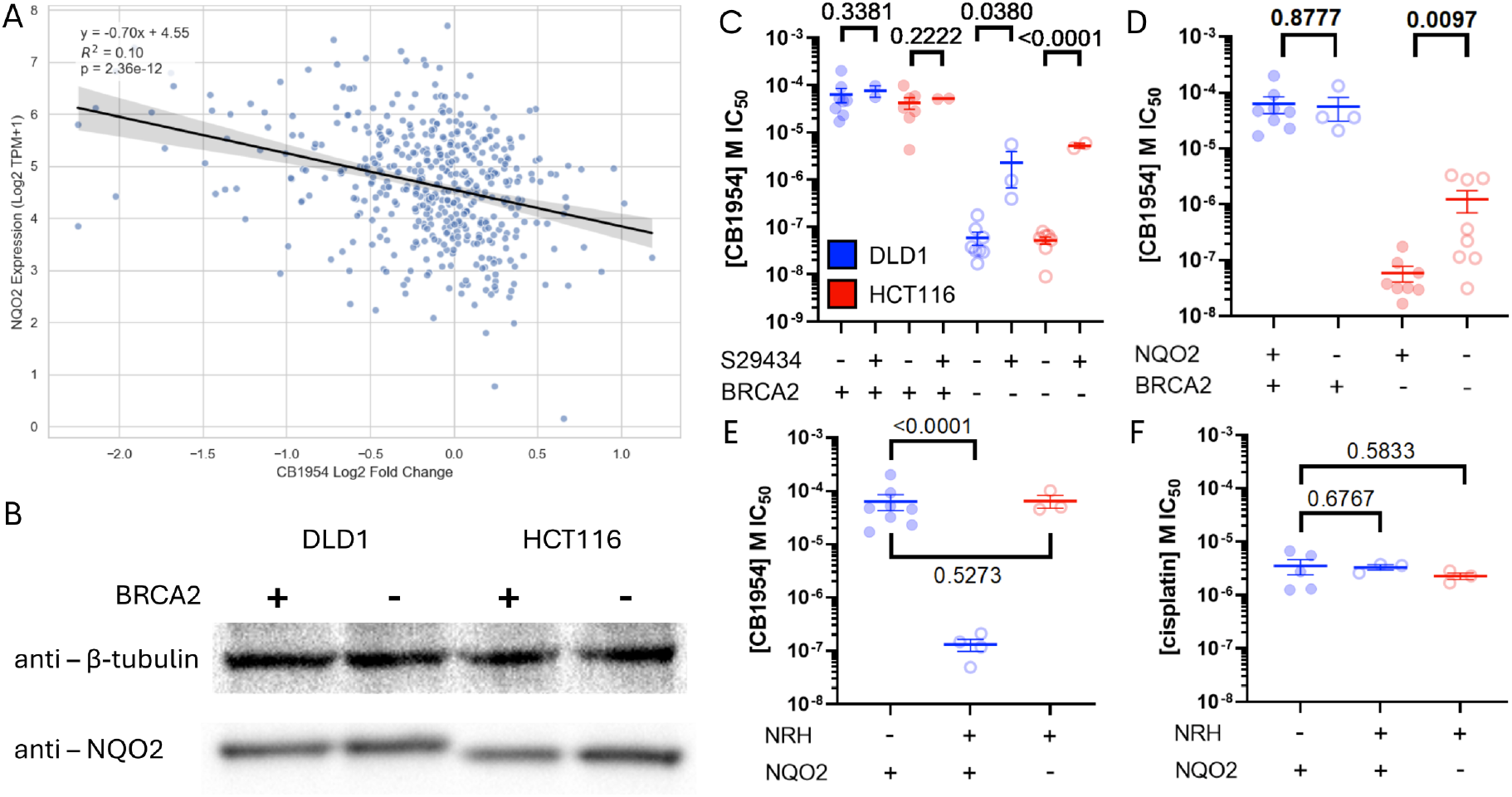
NQO2 modulates CB1954 sensitivity in BRCA2^−/−^ but not HR-proficient isogenic models. **A)** DepMap correlation of NQO2 expression and CB1954 (DepMap name: tretazicar) sensitivity. Points are profiled cell lines. Pearson t-test for linear model statistical significance. **B)** Western blot for DLD1 ± BRCA2 and HCT116 ± BRCA2 NQO2 with *β*-tubulin as a loading control. **C)** IC_50_ of DLD1 ± BRCA2 and HCT116 ± BRCA2 treated with CB1954 with 200 nM of S29434, an NQO2 inhibitor. Mean of biological replicates ± SEM as error bars. Lognormal Welch’s t test for statistical significance. **D)** IC_50_ of DLD1 ± BRCA2 ± NQO2 (single-cell clones) treated with CB1954. Mean of biological replicates ± SEM as error bars. **E)** IC_50_ of DLD1 ± 50 *µ*M NRH ± NQO2 (single-cell clones) treated with CB1954. Mean of biological replicates ± SEM as error bars. **F)** IC_50_ of DLD1 ± NRH ± NQO2 (single-cell clones) treated with cisplatin. Mean of biological replicates ± SEM as error bars.

First, concurrent treatment with CB1954 and S29434, a potent and specific NQO2 inhibitor [26], partially rescued viability in both BRCA2^−/−^ cell lines. Surprisingly, this inhibition had no effect on the sensitivity of either HR-proficient parental line (Figure 4C). To rigorously validate this phenotype, we utilized CRISPR-Cas9 ribonucleoproteins to generate NQO2 knockout clones in the DLD1 ± BRCA2 background. Genetic ablation of NQO2 mirrored the inhibitor data, yielding a partial rescue of CB1954 sensitivity in the BRCA2^−/−^ cells while leaving the parental response unaltered (Figure 4D).

To test whether NRH-induced chemosensitization operated exclusively through the CB1954–NQO2 axis, we evaluated the DLD1 ± 50 *µ*M NRH ± NQO2 models. The sensitization induced by NRH was entirely abrogated in the NQO2 knockout clones (Figure 4E). Finally, neither the addition of NRH nor the ablation of NQO2 altered sensitivity to cisplatin, a crosslinker that does not interact with the NQO2/NRH axis (Figure 4F).

Taken together, these genetic and pharmacological data support the dual-lesion model of cytotoxicity: aziridine monoadducts primarily drive toxicity in HR-proficient cells, whereas bioactivated CB1954 forms the low abundance ICLs responsible for the hypersensitivity observed in HR-deficient cells.

### Structural determinants of CB1954 selectivity and cytotoxicity

To chemically validate our dual-lesion hypothesis and identify the structural features underlying the HRD therapeutic index, we synthesized ten CB1954 analogs to probe its four primary functional groups: the 1-benzamide, 2-nitro, 4-nitro, and 5-(aziridin-1-yl) substituents. Analogs were evaluated via growth delay assays in the DLD1 and HCT116 isogenic pairs (Figure 5A).

**Figure 5.**
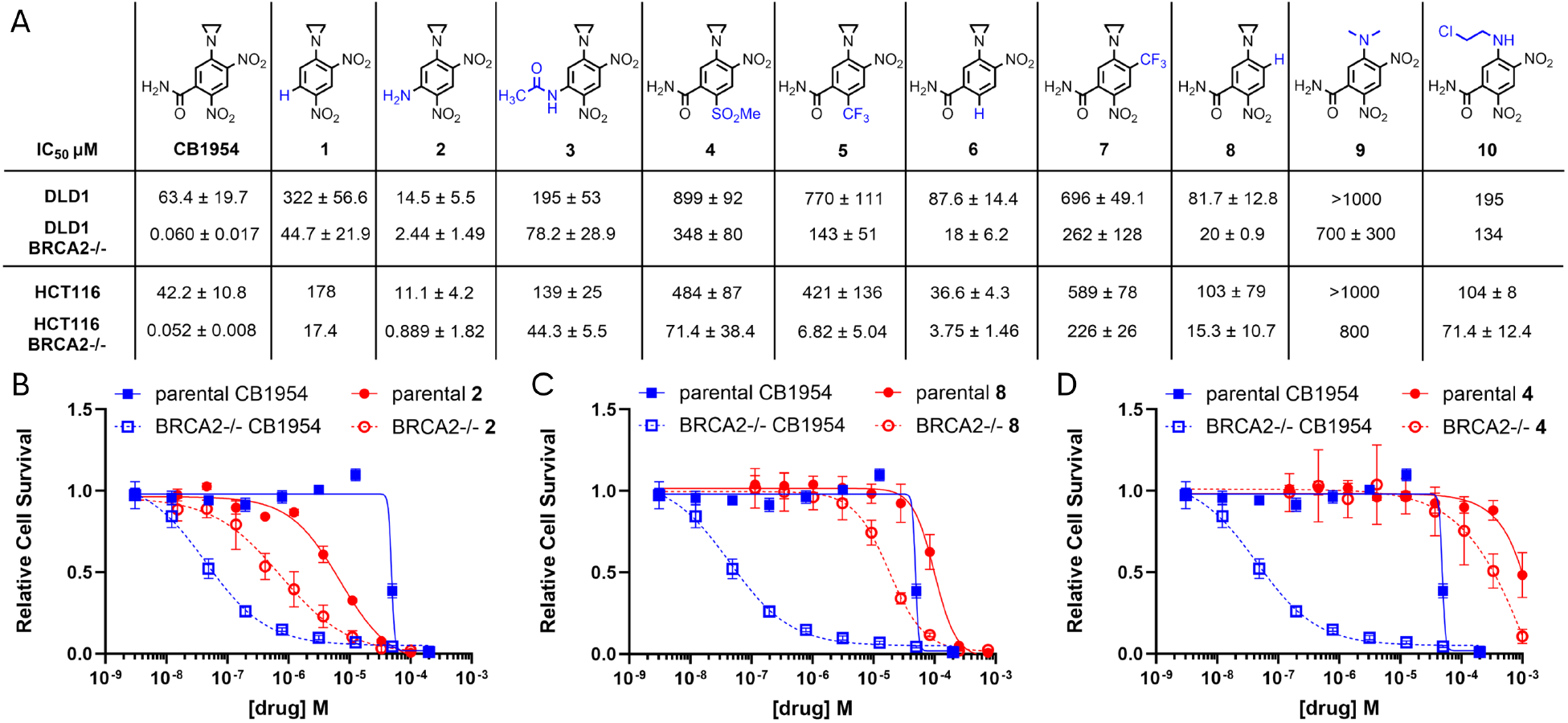
CB1954 structure-activity relationship studies reveal key dependencies for selectivity and potency. **A)** Structures of synthesized derivatives with variable functional groups in blue. Mean IC_50_ for DLD1 ± BRCA2 and HCT116 ± BRCA2 with SEM as error. No SEM indicates a single biological replicate. **B)** Representative dose-response curves for CB1954 and **2** in DLD1 ± BRCA2 with error bars representing SD. **C)** Representative dose-response curves for CB1954 and **8** in DLD1 ± BRCA2. **D)** Representative dose-response curves for CB1954 and **4** in DLD1 ± BRCA2.

We first examined the 1-benzamide moiety, which, according to structural studies, is crucial for anchoring CB1954 within the NQO2 catalytic pocket via a backbone interaction with Gly149 [27]. Consistent with our model, abolishing this coordination capability by removing the 1-benzamide (**1** and **3**) decreased BRCA2^−/−^ potency by 1000-fold. Crucially, these modifications only increased the IC_50_ in the HR-proficient cells by 4-fold. This divergence supports the hypothesis that aziridine monoadducts drive toxicity in HR-proficient cells, whereas NQO2-dependent ICL formation drives the selectivity for HRD cells. Interestingly, the aniline analog (**2**) exhibited increased potency in the HR-proficient cell lines relative to CB1954, likely due to increased electron density activating the aziridine (*σ*_*ρ*_ = -0.66) [28], though it maintained the trend of decreased potency in the BRCA2^−/−^ context (Figure 5B).

Next, we interrogated the 4-nitro group. Because the reduced 4-hydroxylamino species is reported to be one of the two electrophilic centers directly participating in DNA alkylation [29], its removal should abrogate ICL-driven BRCA2^−/−^ sensitivity while sparing monoadduct-driven HR-proficient sensitivity. Indeed, replacing the 4-nitro with a 4-hydrogen (**8**) resulted in a 300-fold reduction in BRCA2^−/−^ potency, but only a 2-fold reduction in the HR-proficient lines (Figure 5C). However, substitution with a 4-trifluoromethyl group (**7**) effectively abrogated potency across all contexts. We suspect the steric bulk of the 4-trifluoromethyl group *ortho* to the 5-aziridin-1-yl hinders nucleophilic attack by nucleobases, while its irreducible electron-withdrawing potential further stabilizes the aziridine ring, disfavoring the reactive aziridinium intermediate.

This functional dependence on aziridine activation was further corroborated by studies of the 2-nitro group. Three analogs (**4**–**6**) lacking a reducible electron-withdrawing group *para* to the aziridine entirely lost selectivity, exhibiting a 1000-fold increase in their BRCA2^−/−^ IC_50_ (Figure 5D). Analogs **4** and **5** bear 2-sulfonyl and 2-trifluoromethyl groups, respectively, which possess Hammett electron-withdrawing potentials (*σ*_*ρ*_ = 0.72 and *σ*_*ρ*_ = 0.54) comparable to the native nitro substituent (*σ*_*ρ*_ = 0.78) [28]. This suggests that the baseline reactivity of the 5-aziridin-1-yl group is contingent upon the reduction of the 2-nitro residue; irreducible electron-withdrawing groups effectively trap the aziridine in an inactive state.

Finally, we validated the absolute requirement for covalent DNA alkylation by replacing the 5-aziridin-1-yl group. The 5-dimethylamino analog (**9**) exhibited a near-complete loss of cytotoxicity, with IC_50_ approaching or exceeding 1000 *µ*M across all cell lines. This demonstrates that the inherent DNA reactivity of the 5-aziridin-1-yl is strictly required for toxicity in either genetic context. This would argue against mechanisms wherein CB1954 acts as a non-covalent inhibitor of an undefined protein to induce these phenotypes. Furthermore, substituting the aziridine with a 5-chloroethylamino group (**10**) eliminated the therapeutic window entirely, vastly increasing the BRCA2^−/−^ IC_50_ while leaving the HR-proficient sensitivity largely unaffected.

### Preclinical ADME profiling, pharmacokinetics, and *in vivo* efficacy

Having established the structural and mechanistic basis for CB1954’s selectivity *in vitro*, we next evaluated its translational potential. We first profiled its preliminary *in vitro* absorption, distribution, metabolism, and excretion (ADME) properties to ensure systemic viability (Figure 6A). CB1954 exhibited high aqueous solubility at pH 7.4 (> 300 *µ*M) and a log*D*_7.4_ of 0.2. Despite possessing a high topological polar surface area (TPSA = 149.7) and a hydrogen bond donor, CB1954 demonstrated excellent permeability in the colon carcinoma Caco-2 model (*P*_*app*_ = 123 nm/sec), with no evidence of active efflux (efflux ratio < 2). Crucially, in both human and mouse hepatocytes, CB1954 exhibited low clearance (< 2.7 *µ*L/min/10^6^ cells) and a robust half-life exceeding 8 hours, despite high expression of the bioactivating enzymes [30, 31]. This extended stability was mirrored in both human and mouse plasma, confirming that the prodrug resists premature systemic degradation.

**Figure 6.**
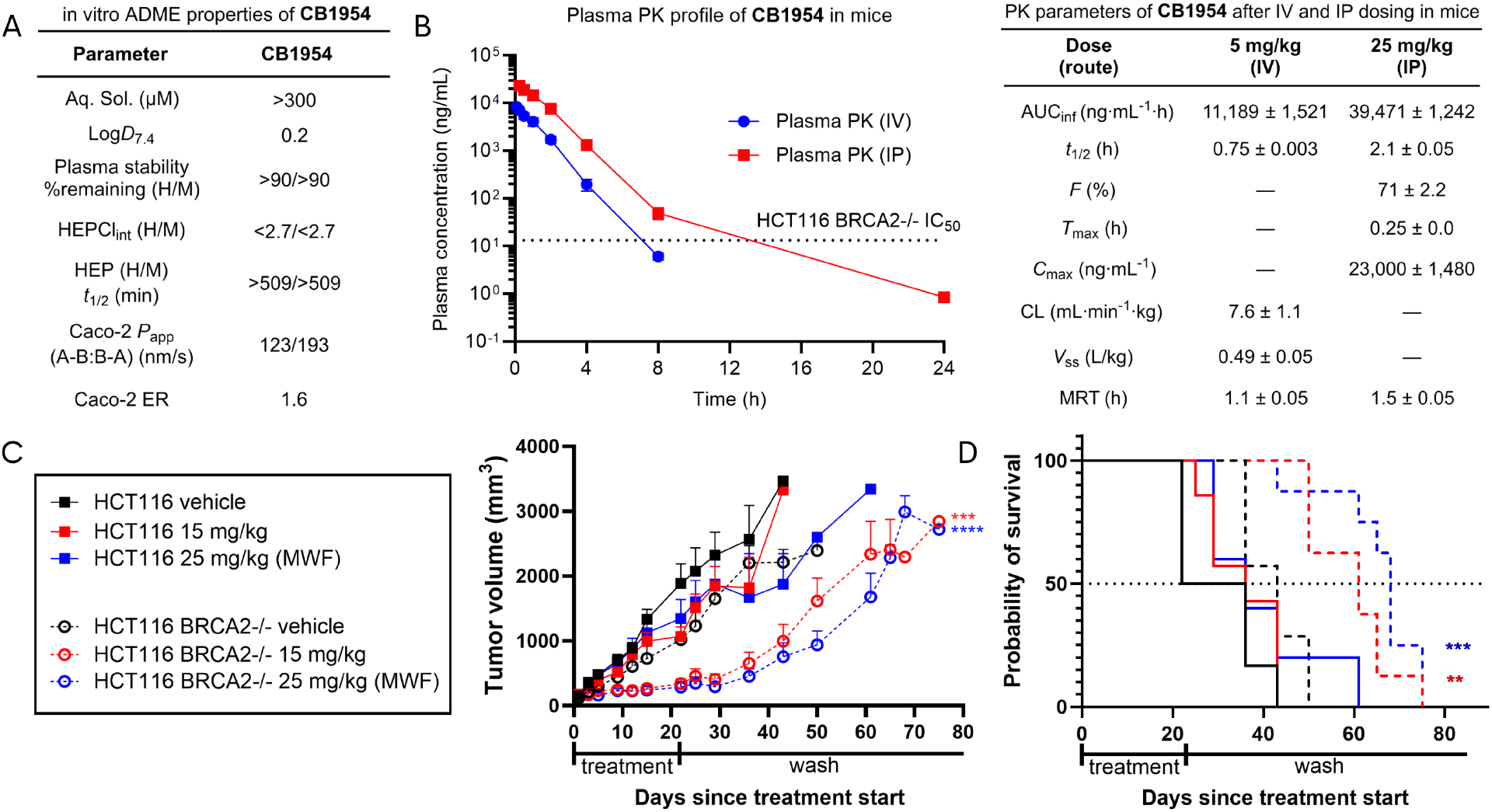
ADME, pharmacokinetics, and *in vivo* efficacy of CB1954. **A)** *In vitro* physicochemical properties and ADME profile of CB1954, evaluating aqueous solubility, lipophilicity (log*D*_7.4_), bidirectional permeability in Caco-2 cells, and metabolic stability across human (H) and mouse (M) plasma and hepatocytes. **B)** *In vivo* plasma pharmacokinetic profile and calculated parameters of CB1954 in wild-type mice following single-dose intravenous (IV, 5 mg/kg) or intraperitoneal (IP, 25 mg/kg) administration. Points on the concentration-time curve represent mean plasma concentrations across three mice with SD error bars. **C)** Mean tumor volume for HCT116 ± BRCA2^−/−^ mouse flank xenografts with shared legend. Error bars represent SEM. Statistical significance was determined using a linear mixed-effects model analyzing the Time × Treatment interaction up to Day 61 between BRCA2^−/−^ vehicle and treatment groups. **D)** Kaplan-Meier overall survival analysis defining survival as time to reach a tumor burden of >3000 mm^3^ or death (right). Statistical significance represented by logrank Mantel-Cox test between BRCA2^−/−^ vehicle and treatment groups.

To test whether these highly favorable *in vitro* properties translated to a physiological system, we evaluated the *in vivo* pharmacokinetics of CB1954 in mice following both intravenous (IV) and intraperitoneal (IP) administration (Figure 6B). CB1954 demonstrated excellent systemic exposure and bioavailability, exhibiting an *in vivo* half-life of 2.1 hours in IP and a clearance rate of 7.6 mL/min/kg in IV. Notably, IP delivery resulted in a peak plasma concentration (C_*max*_) of 91 *µ*M. This safely tolerated systemic exposure exceeds the HCT116 BRCA2^−/−^ IC_50_ (52 ± 8 nM) by more than 1000-fold, establishing an actionable pharmacokinetic therapeutic window.

Capitalizing on this systemic margin, we proceeded to test the *in vivo* efficacy of CB1954 using murine flank models. Athymic BALB/c mice bearing either HCT116 or HCT116 BRCA2^−/−^ cell-line-derived xenografts were treated with CB1954 or vehicle for three one-week cycles (15 mg/kg M-F or 25 mg/kg MWF) once tumors exceeded 100 mm^3^. Mirroring our *in vitro* therapeutic index, CB1954 treatment exerted little to no effect on the growth kinetics of HR-proficient HCT116 parental tumors yet drove a significant anti-tumor response in the HCT116 BRCA2^−/−^ xenografts (Figure 6C). Individual mouse tumor volumes are provided in the supplementary materials (Figures S5A, S5B). Mice treated with CB1954 lost body weight, but quickly recovered after treatment cessation (Figure S5C). Treatment with CB1954 significantly extended median overall survival in only the BRCA2^−/−^ groups, yielding extensions of 18 days for the 15 mg/kg (M-F) cohort and 25 days for the 25 mg/kg (MWF) cohort (Figure 6D).

## Discussion

The present study identifies CB1954 as a highly selective DNA-damaging agent for HR-deficient tumor cells and redefines the mechanism by which this historically abandoned prodrug can be exploited therapeutically. Although CB1954 has long been recognized as a bioactivated DNA-reactive compound, its prior clinical development focused primarily on the challenge of tumor-selective activation. Here, we show that an additional and previously underappreciated determinant of selectivity is the DNA repair genotype of the tumor. When evaluated in HR-deficient models, CB1954 displays a striking therapeutic index, induces selective DNA-damage signaling and apoptosis, and produces genotype-dependent antitumor activity *in vivo*.

These findings provide a mechanistic explanation for why CB1954 may have been overlooked. Earlier efforts treated CB1954 primarily as a prodrug whose success depended on achieving selective enzymatic activation in tumor tissue. This led to virus-directed enzyme-prodrug therapy strategies designed to force nitroreductase expression in tumors. However, those studies were conducted outside a modern HRD biomarker framework. Our data suggest that the biological context most relevant to CB1954 activity is not only the level of bioactivation, but also the ability of the tumor cell to repair the resulting DNA lesions. In HR-proficient cells, CB1954-induced damage appears to be more tolerable and may be dominated by lesions that generate replication stress without efficient conversion to lethal double-strand breaks. In contrast, HR-deficient cells are unable to resolve the critical repair and replication intermediates generated by CB1954, resulting in profound cytotoxicity.

A central conclusion from this work is that CB1954 does not behave like a conventional, broadly acting DNA-crosslinking agent. Although CB1954 forms ICLs in both HR-proficient and HR-deficient cells, its cytotoxic consequences are sharply genotype-dependent. Targeted CRISPR screening identifies Fanconi anemia and HR pathway genes as key mediators of CB1954 sensitivity, consistent with a lesion-repair model in which CB1954 imposes an obligate FA/HR repair burden. In contrast to platinum agents, which generate diverse lesions processed by multiple pathways including nucleotide excision repair and mismatch repair, CB1954 shows limited dependence on these alternative pathways in our genetic screens. This narrower repair dependency likely contributes to the unusually large selectivity window observed in HR-deficient cells.

The NQO2 data further refine this model. Genetic loss or pharmacologic inhibition of NQO2 partially rescues CB1954 sensitivity in BRCA2-deficient cells, while having little effect on HR-proficient cells. These results support a bifurcated mechanism in which NQO2-dependent bioactivation generates low-abundance but highly lethal crosslinking lesions that are selectively toxic in HR-deficient cells. In parallel, aziridine-dependent monoadducts or related lesions may account for the residual toxicity observed in HR-proficient cells at high concentrations. This dual-lesion framework is also supported by the structure-activity relationship data, which show that functional groups required for bioactivation and DNA alkylation differentially influence potency in HR-deficient versus HR-proficient contexts.

From a translational perspective, these findings suggest that CB1954 deserves reconsideration as a genotype-directed DNA-damaging agent rather than as an unselected cytotoxic prodrug. The favorable *in vitro* ADME profile, systemic exposure achieved in mice, and selective *in vivo* efficacy in BRCA2-deficient xenografts support further optimization and preclinical evaluation. Importantly, the goal would not be to revive CB1954 exactly as it was historically developed, but to apply modern biomarker logic, repair-pathway biology, and medicinal chemistry to a chemotype whose original clinical testing preceded the HRD era.

Several questions remain. Future studies will define the precise chemical structures and abundance of the DNA lesions generated by CB1954 in cells, determine whether CB1954 retains activity in models of acquired PARP inhibitor or platinum resistance, and evaluate its activity in additional HRD contexts beyond engineered BRCA2 loss. It will also be important to determine whether NQO2 expression or activity can serve as a pharmacodynamic or patient-selection biomarker alongside HRD status. Nonetheless, the present work establishes a clear conceptual advance: CB1954 is not simply an old cytotoxin, but a historically overlooked HRD-selective agent whose therapeutic potential becomes apparent only when viewed through the lens of DNA repair deficiency.

## Materials and Methods

### Cell culture

DLD1 and HCT116 human colorectal carcinoma cells were cultured in RPMI 1640 (Gibco, 11875119) and McCoy’s 5A Modified medium (Gibco, 16600108), respectively. U2OS and LN229 cells were cultured in McCoy’s 5A Modified medium and DMEM, high glucose medium (Gibco, 11965092), respectively. All media were supplemented with 10% (v/v) fetal bovine serum (FBS; Sigma-Aldrich, F0926) without antibiotics. Cells were maintained at 37°C in a humidified 5% CO_2_ atmosphere. All cell lines were authenticated by short tandem repeat (STR) profiling (>80% match) and routinely tested negative for mycoplasma contamination.

### Growth delay assay

Cells in the logarithmic growth phase were seeded in 96-well plates at densities of 1,000–2,000 cells/well, depending on the doubling time, and allowed to adhere for 24 hours. Cells were subsequently treated with a concentration gradient of the indicated compounds or vehicle control (1% final solvent concentration) for 120 hours. Following treatment, cells were washed with Dulbecco’s Phosphate-Buffered Saline (DPBS; Gibco, 14190144) and fixed with 4% formaldehyde (Thermo Scientific, 033314AP). Nuclei were stained with 1 *µ*g/mL Hoechst 33342 (Invitrogen, H1399) for 30 minutes. Plates were imaged using a Cytation 3 Cell Imaging Multi-Mode Reader (BioTek) with a DAPI filter set. Nuclei were quantified and analyzed using CellPyAbility software (https://github.com/bindralab/cellpyability).

### Clonogenic survival assay

Cells were treated with the indicated drug concentrations or vehicle control for 72 hours, subsequently harvested, and seeded into 6-well plates (Corning, 353046) at serial ten-fold dilutions (50 to 50,000 cells/well). After 14 days of undisturbed incubation, colonies were washed with DPBS and simultaneously fixed and stained with a solution of 0.01% (w/v) Coomassie Brilliant Blue (Thermo Scientific, 20278) in 50% (v/v) methanol and 10% (v/v) acetic acid for 30 minutes. Plates were rinsed with cold water and air-dried. Colony counting was performed in a blinded manner. Colonies containing ≥50 cells were scored as positive. Survival fractions were calculated by normalizing to the plating efficiency of the vehicle-treated controls.

### *In vitro* interstrand crosslink (ICL) detection gel

In 100 *µ*L of total volume, CB1954 [X *µ*M] (Selleck Chem S7829), NRH [2 mM] (MedChemExpress HY-W614507), acetyl-CoA [10 mM] (Sigma-Aldrich A2056), NQO2 [50 nM] (Reaction Biology NQO-11-312), and linearized pUC19 DNA (500 ng) (Thermo Scientific SD0061) were mixed and incubated for 16 hours at 37°C in DPBS. CB1954 stock concentration is adjusted so the final volume of the reaction mixture is 1% DMSO. Samples were denatured in 50% DMSO at 80 °C for 2 minutes, then placed immediately on ice and loaded into a 0.6% (w/v) agarose gel with 0.5 *µ*g/mL ethidium bromide in TAE buffer. The gel was run at 8 V / L cm, where L is the distance between electrodes, for one hour.

### Reverse comet assay

Alkaline single-cell gel electrophoresis with ionizing radiation assays were performed using the CometAssay kit (R&D Systems, 4250-050-ESK) according to the manufacturer’s protocol, with modifications for ionizing radiation (IR). Cells were seeded in 6-well plates to reach ∼70% confluence after 24 hours, followed by treatment with the indicated DNA-damaging agents or vehicle control. Mitomycin C (0.1 *µ*M; Sigma-Aldrich, M5353) was utilized as a positive control for ICLs. Post-treatment, cells were harvested in cold DPBS, resuspended at 1 × 10^4^ cells/mL in 40°C low-melting-point agarose, and spread onto CometSlides. After 15 minutes at 4°C to solidify, slides were incubated in lysis buffer overnight at 4°C. Following a wash in TE buffer, slides were subjected to 10 Gy irradiation using an X-RAD irradiator to induce uniform DNA fragmentation. Slides were then equilibrated in pre-chilled alkaline electrophoresis buffer for 1 hour at 4°C and electrophoresed at 21 V for 50 minutes using the CometAssay Electrophoresis System II. Slides were washed in TE buffer, dehydrated in ice-cold 70% ethanol for 10 minutes, and air-dried. DNA was stained with 0.3 × SYBR-Gold (Invitrogen, S11494) for 30 minutes at room temperature, washed twice with TE buffer, and imaged via fluorescence microscopy. All procedures were conducted under minimal lighting to prevent ambient DNA damage.

### High-throughput immunofluorescence foci assays

High-throughput immunofluorescence foci assays were performed at the Yale Center for Molecular Discovery (YCMD). Cells were seeded at 2,000 cells/well in black polystyrene flat bottom 384-well plates (Greiner Bio-One) and allowed to adhere overnight. Compound addition was performed utilizing a Labcyte Echo 550 liquid handler (Beckman Coulter), with 6 replicates per test condition and 12 replicates per control condition. Following drug incubation, cells were fixed and stained for phospho-SER139-H2AX (*γ*H2AX) or 53BP1.

For the *γ*H2AX protocol, cells were fixed with 4% paraformaldehyde in 1× PBS for 15 minutes, washed twice with 1×PBS, incubated in extraction buffer (0.5% Triton X-100 in 1× PBS) for 10 minutes, washed twice with 1× PBS, and incubated in blocking buffer (Blocker Casein in PBS, Thermo Scientific + 5% goat serum, Life Technologies) for 1 hour. Mouse anti-phospho-histone H2A.X (Ser139) antibody (clone JBW301, Millipore, 05-636) was added 1:1000 in blocking buffer at 4 °C overnight. After washing with 1× PBS, cells were incubated with goat anti-mouse IgG (H+L) highly cross absorbed secondary antibody, Alexa Fluor 647 (Invitrogen, A-21236) 1:500 and with 1 *µ*g/mL Hoechst nucleic acid dye in blocking buffer for 2 hours, and then washed with 1× PBS.

For the 53BP1 protocol, cells were fixed with 4% paraformaldehyde + 0.02% Triton X-100 in 1 × PBS for 20 minutes, washed twice with 1 × PBS, and incubated in blocking buffer (10% FBS, 0.5% Triton X-100 in 1× PBS) for 1 hour. Rabbit anti-53BP1 antibody (Novus Biologicals, NB100-904) was added 1:1000 in blocking buffer at 4 °C overnight. After washing with 1 × PBS, cells were incubated with goat anti-rabbit IgG (H+L) highly cross-adsorbed secondary antibody, Alexa Fluor 647 (Invitrogen, A-21245) 1:500 and with 1 *µ*g/mL Hoechst nucleic acid dye in blocking buffer for 2 hours, and then washed with 1× PBS.

Imaging was performed on an ImageXpress Micro 4 (Molecular Devices) at 40 × magnification. Twenty fields-of-view were captured per well. Foci analysis was performed using MetaXpress, and outer wells were excluded from analysis to limit variation due to edge effects.

### DDR CRISPR screening

The CRISPR assay was adapted from [22]. In brief, Cas9-expressing LN229 cells were combined with the packaged library at low MOI (< 0.3) and 8 µg/mL Polybrene (Sigma–Aldrich, TR-1003) and plated into 15 cm dishes (Corning, 430599). Medium was replaced the next day.

After 48–72 h, infected cells were selected by exposure to 750 *µ*g of hygromycin (Gibco, 10687010) for 96–120 h. Following hygromycin selection (Time 0), coverage was calculated, and cells were then split into control and treatment groups. Each condition was performed in duplicate and plated at ≥600-fold coverage. In total, cells were exposed to chemical agents for 12 d and received 4 doses of each agent (days 0, 3, 6, and 9).

After 72 h, cells were trypsinized, counted with an automated cell counter (Bio-Rad TC20), replated at desired coverage, and retreated. This procedure was repeated on days 6 and 9. On day 12, cell pellets (consisting of 2–3 million cells) were collected for each sample and washed with DPBS. The DPBS was removed by aspiration and the cells were flash-frozen and stored at -80 °C.

Sequencing preparation and analysis was performed as previously described [22]. Equal masses ( ≥ 20 ng of amplicon) of each sample were combined into a single tube prior to submission for sequencing. 150 bp paired-end sequencing was performed on the NovaSeq X Plus (Illumina) at the Yale Center for Genome Analysis (YCGA), except for CB1954 0.6 *µ*M batch which used the NovaSeq 6000. Significant changes in sgRNA abundance between different conditions were determined using MAGeCK-RRA algorithm v0.5.9.4 [32] with median normalization.

### Generation of stable knockout cell lines

Isogenic knockout cell lines for LN229 Cas9, DLD1, and DLD1 BRCA2^−/−^ were generated following the EditCo RNP protocol. In brief, high-efficiency pools of sgRNAs for BRIP1 and NQO2 were ordered from EditCo. Cells were reverse transfected with either RNP in the DLD1 background or sgRNA pool alone in the LN229 Cas9 background due to high stable Cas9 expression. Cells were single-cell-flow-sorted into 96-well plates, then expanded and assessed for gene knockout, first with PCR, then Sanger sequencing, and finally Western blot.

### Chemical synthesis and analysis

General chemical experimental procedures are described in detail in the supplementary information, along with the catalog of nuclear magnetic resonance spectra.

### *In vitro* ADME and *in vivo* PK

All *in vitro* absorption, distribution, metabolism, excretion (ADME) and *in vivo* PK studies were adapted from industry-standard protocols and performed at Pharmaron Inc. We provide detailed descriptions in the supplemental material.

### Flank cell-line derived xenograft studies

Athymic nude mice (Hsd:Athymic Nude-*Foxn1*^nu^) were purchased from Envigo. HCT116 ± BRCA2 cells were used to establish tumor models by subcutaneous implantation. Briefly, HCT116 cells (1 × 10^6^ parental or 2 × 10^6^ BRCA2^−/−^ cells in 50 *µ*L DPBS) were diluted in 50 *µ*L Matrigel (Corning) and were injected into the flanks of female nude mice. Treatment was initiated when the average tumor volume reached approximately 100–300 mm^3^, at which time mice were randomly assigned to treatments while enforcing similar average tumor volume across groups and assigned to treatment groups. Mice were dosed by intraperitoneal (IP) administration daily with vehicle control (10:10:80 DMSO:solutol HS15:dextrose 5% in H_2_O) or CB1954 at 15 mg/kg M-F, or 25 mg/kg MWF for 3 cycles. Tumors were measured using calipers and tumor volume was calculated using the formula: *V* = 0.52 × Length × Width^2^. Mice were euthanized if body weight loss exceeded 20%, if tumor volume exceeded 3000 mm^3^, or for other humane considerations like ulceration. All experimental procedures were performed according to Yale University Institutional Animal Care and Use Committee (IACUC) guidelines.

### Statistical analysis

Statistical analysis was performed using GraphPad Prism software unless otherwise noted. Non-parametric tests were used when possible. Data are presented as mean ± SD or SEM as indicated. For growth delay assays, IC_50_ values were determined using CellPyAbility software (https://github.com/bindralab/cellpyability) by applying a fitted 5-parameter logistic followed by the algebraic inverse to solve for *x* when *y* = 0.5 and *D* < 0.5 ≥ *A*, where *x* is the concentration of drug, *y* is the relative cell survival, *A* is the maximum of the logistic, and *D* is the minimum of the logistic. HRD selectivity index was calculated by log-transformed paired analysis of biological replicates. For xenograft experiments, comparisons for survival were made using a logrank Mantel-Cox test.

## Acknowledgements

Financial support from the NIH-NCI (5R01CA276186-02 to S.B.H. and R.S.B.) and from Yale University (Yale Cancer Center Class of ‘61 Cancer Research Award to S.B.H.) is gratefully acknowledged. J.L.E. is supported by the National Institutes of Health through the Yale Cancer Biology Training Program (T32CA193200) and the Ruth L. Kirschstein Predoctoral Individual National Research Service Award (F31CA294600). J.H. is supported by a postdoctoral fellowship from the Jane Coffin Childs Memorial Fund for Medical Research. C.D.H. is supported by a postdoctoral fellowship from the NIH (K00CA245722). This research made use of the Chemical and Biophysical Instrumentation Center at Yale University (RRID:SCR021738). Equipment was purchased with funds from Yale University. This work utilized ImageXpress Micro 4 high content imager that was purchased with funding from a NIH SIG grant 1S10OD032384, and Echo 650 Acoustic Liquid Handler that was purchased with funding from a National Institutes of Health SIG grant 1S10OD038347. Research reported in this publication was supported by the National Institute of General Medical Sciences of the National Institutes of Health under award Number (1S10OD030363-01A1). We thank Dr. Ryan Jensen (Yale University) for generously sharing isogenic cell line models. We thank the Yale Center for Genome Analysis for assistance with the CRISPR screen, and specifically Dr. Francesc Lopez-Giraldez. We thank Dr. Yulia Surovtseva (Yale University) and Sheila Umlauf (Yale University) for their assistance with DDR foci and cell cycle analysis experiments. We thank Yale Flow Cytometry for their assistance with single-cell sorting and Annexin V studies; the Core is supported in part by an NCI Cancer Center Support Grant NIH P30 CA016359. The BD Symphony was funded by shared instrument grant NIH S10 OD026996. We thank the scientists at Pharmaron Inc. for their help in conducting the *in vitro* ADME and *in vivo* PK experiments.

