## Supplementary Figures for "Leveraging Homologous Recombination Deficiency via the Repositioned Prodrug CB1954"

A bioRχiv Preprint

James L. Elia[1,†], Jarvis Hill[2,†], Collin D. Heer[3], Siji Smolev[3], Abbey M. Sykes[1], Sofia R. Arbelaez[3], Karlie N. Lucas[3], Spenser S. Johnson[3], Ranjini K. Sundaram[3], Seth B. Herzon[2,3,4,‡], and Ranjit S. Bindra[1,3,‡]

[1]Department of Pathology, Yale University, New Haven, CT, USA
[2]Department of Chemistry, Yale University, New Haven, CT, USA
[3]Department of Therapeutic Radiology, Yale University, New Haven, CT, USA
[4]Department of Pharmacology, Yale University, New Haven, CT, USA
[†]These authors contributed equally
[‡]Co-senior authors

July 8, 2026

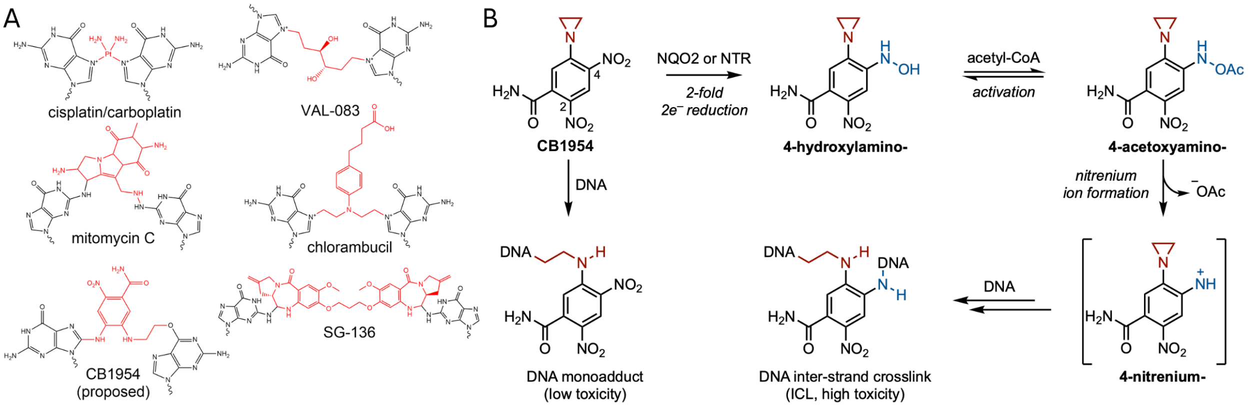


Figure S1: Chemical schema for the DNA crosslinker viability screen and CB1954 bioactivation. **A)** Reported and proposed structures for clinically relevant DNA crosslinkers. **B)** Literature-proposed CB1954 bioactivation mechanism.

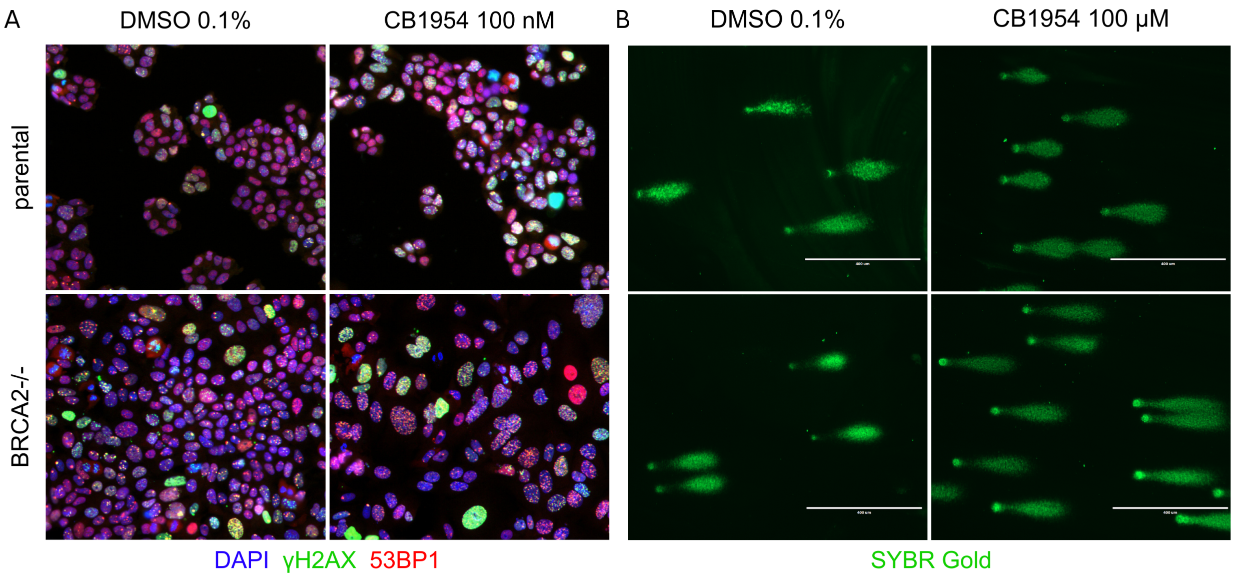


Figure S2: Representative DDR foci and reverse comet images. **A)** Immunofluorescence microscopy images of DLD1 $\pm$ BRCA2 treated with vehicle or CB1954 100 nM for 48 h. **B)** Reverse comet assay images of DLD1 $\pm$ BRCA2 treated with vehicle or CB1954 100 $\mu$M for 24 h.

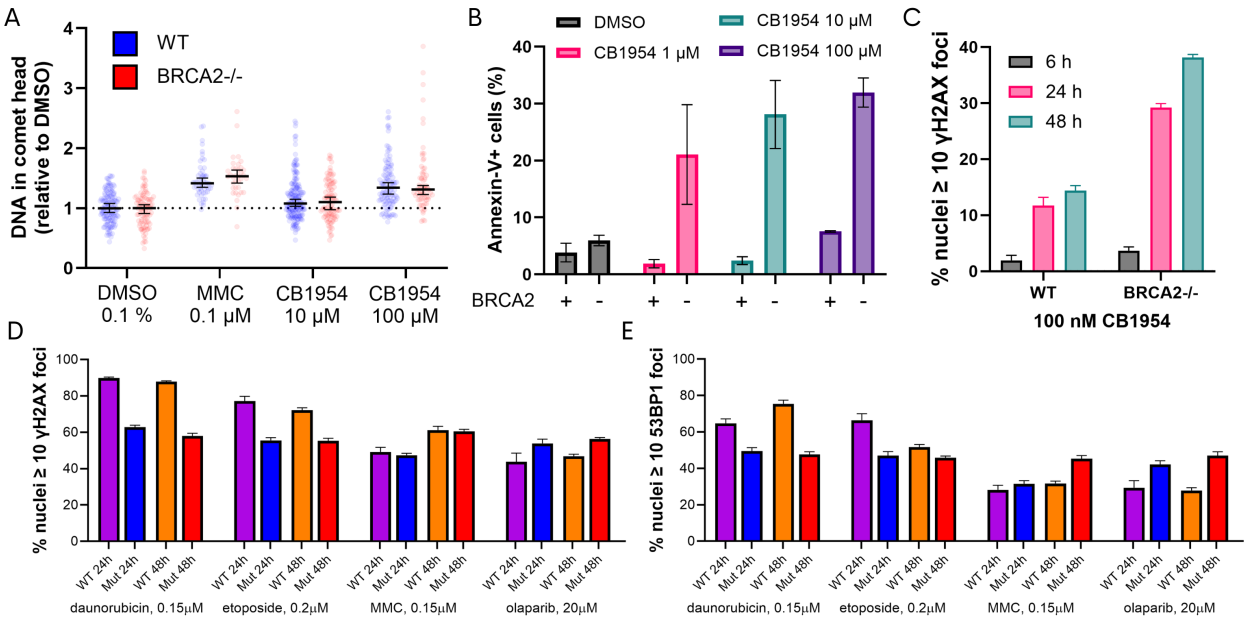


Figure S3: CB1954 cellular mechanism extended data. **A)** Reverse comet assay for HCT116 $\pm$ BRCA2. Median of all nuclei with 95% confidence intervals as error bars. **B)** Flow cytometry analysis of Annexin V+ DLD1 $\pm$ BRCA2 in response to 72 hours of indicated concentrations of CB1954. Mean of biological replicates with SEM as error bars. **C)** High-throughput immunofluorescence $\gamma$H2AX+ DLD1 $\pm$ BRCA2 at 6 h, 24 h, and 48 h after treatment with 100 nM CB1954. Mean of technical replicates with SD as error bars. **D)** High-throughput immunofluorescence for $\gamma$H2AX+ and **E)** 53BP1+ DLD1 $\pm$ BRCA2 after treatment with indicated concentrations of control compounds. Mean of technical replicates with SD as error bars.

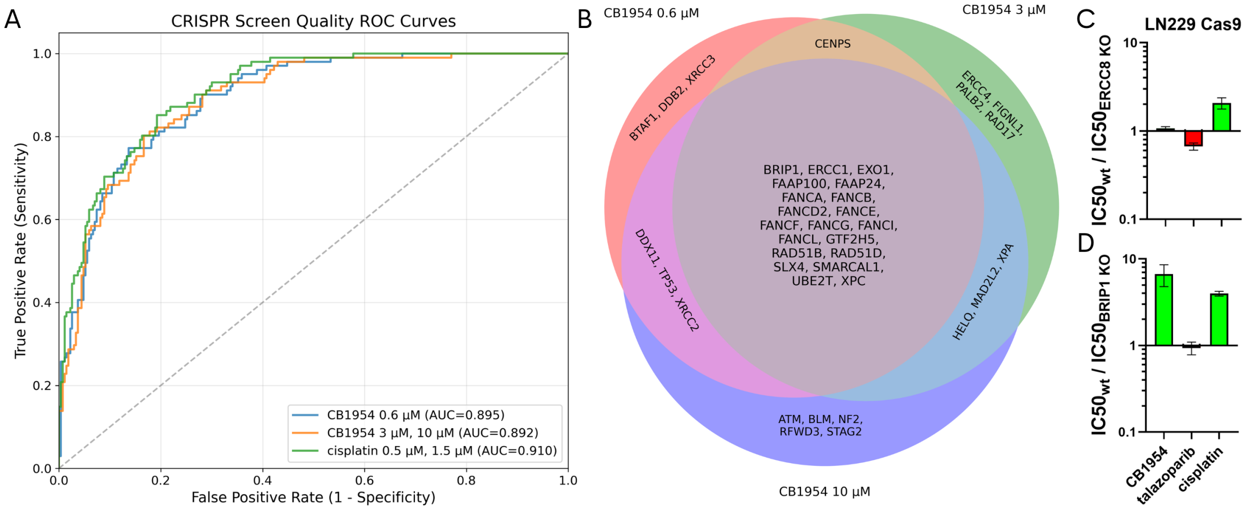


Figure S4: DDR CRISPR screen extended data. **A)** Receiver-operator curves with area under the curve (AUC) for each batch of CRISPR screen. True positives represent essential genes that cause dropout upon loss between Day 0 and Day 12 vehicle. **B)** Venn diagram of the three CB1954 DDR CRISPR screens. **C)** $IC_{50}$ ratios of LN229 Cas9 $\pm$ ERCC8 with cisplatin, talazoparib, and CB1954. Mean of biological replicates $\pm$ SEM as error bars. **D)** $IC_{50}$ ratios of LN229 Cas9 $\pm$ BRIP1 with cisplatin, talazoparib, and CB1954. Mean of biological replicates $\pm$ SEM as error bars.

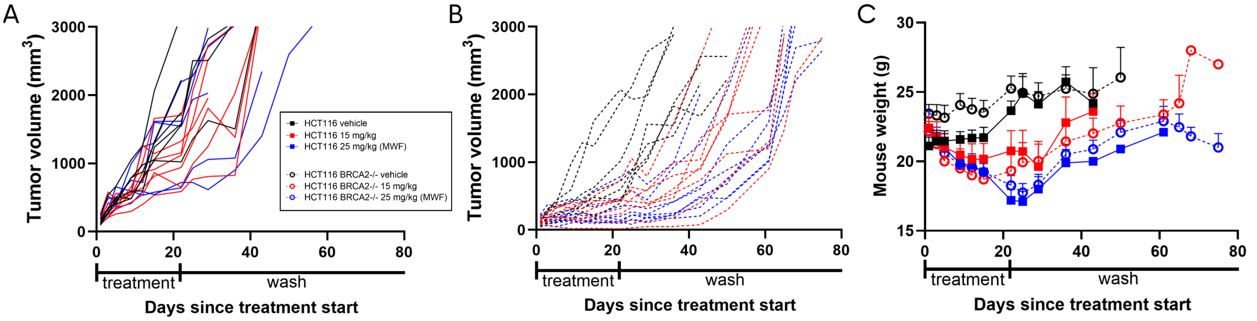


Figure S5: Cell-line derived xenografts extended data. **A)** Individual mouse tumor volumes for HCT116 parental flank xenografts with shared legend. **B)** Individual mouse tumor volumes for HCT116 $BRCA2^{-/-}$ flank xenografts. **C)** Mean mouse body weight in grams. Error bars represent SEM.
