## Supplementary Data for "Leveraging Homologous Recombination Deficiency via the Repositioned Prodrug CB1954"

A bioRχiv Preprint

James L. Elia[1,†], Jarvis Hill[2,†], Collin D. Heer[3], Siji Smolev[3], Abbey M. Sykes[1], Sofia R. Arbelaez[3], Karlie N. Lucas[3], Spenser S. Johnson[3], Ranjini K. Sundaram[3], Seth B. Herzon[2,3,4,‡], and Ranjit S. Bindra[1,3,‡]

[1]Department of Pathology, Yale University, New Haven, CT, USA
[2]Department of Chemistry, Yale University, New Haven, CT, USA
[3]Department of Therapeutic Radiology, Yale University, New Haven, CT, USA
[4]Department of Pharmacology, Yale University, New Haven, CT, USA
[†]These authors contributed equally
[‡]Co-senior authors

July 8, 2026

**This PDF file includes:**

Supplementary Figures (Figs. S1-S5)

Chemistry Materials and Methods

Synthetic Procedures

Catalog of Nuclear Magnetic Resonance Spectra

References (1-5)

**Supplementary Figures.**

| Compound ID | Replicate | Log D |
|---|---|---|
| Progesterone | Replicate 1 | 3.90 |
| | Replicate 2 | 3.85 |
| | Mean | 3.87 |
| CB1954 | Replicate 1 | 0.19 |
| | Replicate 2 | 0.23 |
| | Mean | 0.21 |

| Compound ID | Buffer | Upper limit (μM) | Solubility (μM) |
|---|---|---|---|
| Progesterone | PBS pH 7.4 | 300 | 18.17 |
| CB1954 | PBS pH 7.4 | 300 | 304.55 |

**Fig. S1. Basic physicochemical properties of CB1954.** Aqueous solubility is defined as the kinetic aqueous solubility at pH 7.4.

| Compound ID | Species | Assay Format | Remaining Percentage (%) | | | | | |
|---|---|---|---|---|---|---|---|---|
| | | | 0.5 min | 15 min | 30 min | 45 min | 60 min | 120 min |
| Verapamil | Human | Hepatocytes | 100.00 | 59.58 | 30.77 | 16.83 | 11.08 | 2.70 |
| | | Inactiveated hepatocytes | 100.00 | 99.89 | 93.06 | 95.34 | 95.83 | 99.71 |
| | Mouse | Hepatocytes | 100.00 | 12.25 | 2.32 | 0.00 | 0.00 | 0.00 |
| | | Inactivated hepatocytes | 100.00 | 94.52 | 93.57 | 86.63 | 91.29 | 94.16 |
| CB1954 | Human | Hepatocytes | 100.00 | 92.35 | 94.85 | 93.14 | 89.47 | 89.66 |
| | | Inactivated hepatocytes | 100.00 | 99.55 | 100.83 | 97.81 | 101.70 | 110.51 |
| | Mouse | Hepatocytes | 100.00 | 102.72 | 103.18 | 106.75 | 101.40 | 103.39 |
| | | Inactivated hepatocytes | 100.00 | 97.93 | 102.03 | 90.31 | 98.04 | 108.97 |

**Fig. S2. Raw hepatocyte stability data.**

| Compound ID | Test Concentration (μM) | Species | $T_{1/2}$ (min) | Remaining Percentage (%) | | | | |
|---|---|---|---|---|---|---|---|---|
| | | | | 0 min | 15 min | 30 min | 60 min | 120 min |
| Propantheline | 5 | Human | 26.82 | 100.00 | 71.47 | 49.67 | 21.40 | 3.41 |
| Propantheline | 5 | Mouse | 38.77 | 100.00 | 78.25 | 61.05 | 36.35 | 11.78 |
| CB1954 | 5 | Human | >511.69 | 100.00 | 112.75 | 106.06 | 101.77 | 104.13 |
| CB1954 | 5 | Mouse | >511.69 | 100.00 | 97.90 | 104.77 | 91.46 | 98.80 |

**Fig. S3. Raw plasma stability data.**

| Compound ID | Concentration (µM) | Replicate | A to B Papp (1E-6.cm/s) | B to A Papp (1E-6.cm/s) | Efflux Ratio | A to B Recovery (%) | B to A Recovery (%) |
|---|---|---|---|---|---|---|---|
| Metoprolol | 10 | Replicate 1 | 26.80 | 28.76 | 1.07 | 103.45 | 104.51 |
| | | Replicate 2 | 23.30 | 24.64 | 1.06 | 105.95 | 93.56 |
| | | Mean | **25.05** | **26.70** | **1.07** | **104.70** | **99.04** |
| Erythromycin | 10 | Replicate 1 | 0.09 | 9.26 | 100.16 | 95.61 | 90.08 |
| | | Replicate 2 | 0.09 | 9.24 | 98.75 | 82.87 | 95.61 |
| | | Mean | **0.09** | **9.25** | **99.46** | **89.24** | **92.85** |
| Cimetidine | 10 | Replicate 1 | 0.76 | 6.28 | 8.27 | 119.47 | 103.10 |
| | | Replicate 2 | 0.53 | 4.21 | 7.91 | 84.59 | 87.00 |
| | | Mean | **0.65** | **5.25** | **8.09** | **102.03** | **95.05** |
| CB1954 | 10 | Replicate 1 | 12.40 | 19.47 | 1.57 | 107.52 | 97.99 |
| | | Replicate 2 | 12.15 | 19.02 | 1.57 | 106.99 | 100.61 |
| | | Mean | **12.27** | **19.25** | **1.57** | **107.25** | **99.30** |

**Fig S4. Raw Caco-2 permeability data.**

| Dose | CB1954 (5 mg/kg, IV) | | | | |
|---|---|---|---|---|---|
| **Time** | **Plasma concentration (ng/mL)** | | | **Mean** | **SD** |
| **(h)** | **Mouse 1** | **Mouse 2** | **Mouse 3** | **(ng/mL)** | **(ng/mL)** |
| 0.083 | 8420 | 8220 | 8480 | 8373 | 136 |
| 0.25 | 7360 | 7570 | 6530 | 7153 | 550 |
| 0.5 | 5630 | 6040 | 4560 | 5410 | 764 |
| 1 | 4520 | 4570 | 3160 | 4083 | 800 |
| 2 | 1930 | 1830 | 1390 | 1717 | 287 |
| 4 | 191 | 253 | 145 | 196 | 54 |
| 8 | 6.57 | 6.70 | 4.97 | 6.08 | 0.96 |
| 24 | BLOQ | BLOQ | BLOQ | NA | NA |

| Dose | CB1954 (25 mg/kg, IP) | | | | |
|---|---|---|---|---|---|
| **Time** | **Plasma concentration (ng/mL)** | | | **Mean** | **SD** |
| **(h)** | **Mouse 1** | **Mouse 2** | **Mouse 3** | **(ng/mL)** | **(ng/mL)** |
| 0.25 | 24700 | 22300 | 22000 | 23000 | 1480 |
| 0.5 | 19800 | 18600 | 18200 | 18867 | 833 |
| 1 | 15700 | 14600 | 13400 | 14567 | 1150 |
| 2 | 7280 | 7910 | 7410 | 7533 | 333 |
| 4 | 1340 | 1200 | 1380 | 1307 | 94.5 |
| 8 | 43.2 | 40.2 | 64.0 | 49.1 | 13.0 |
| 24 | 0.958 | 0.654 | 0.926 | 0.846 | 0.17 |

**Fig. S5. Raw data from single-dose pharmacokinetic study in male C57BL/6 mice.** BLOQ, below level of quantitation.

**Chemistry Materials and Methods.**

General Chemical Experimental Procedures

All reactions were performed in single-neck, oven-dried, round-bottom flasks fitted with rubber septa under a positive pressure of argon unless otherwise stated. Organic solutions were concentrated by rotary evaporation at 23-35 °C. Flash-column chromatography was performed as described by Still et al., employing silica gel ('SiliaFlash® P60', 60 Å, 40-63 $\mu$m particle size) purchased from SiliCycle (Quebec, Canada) [1]. Analytical thin-layered chromatography (TLC) was performed using glass plates pre-coated with silica gel (250 $\mu$m, 60 Å pore size) impregnated with a fluorescent indicator (254 nm). TLC plates were visualized by exposure to ultraviolet light (UV) and/or submersion in aqueous potassium permanganate ($KMNO_4$) or aqueous ceric ammonium molybdate (CAM) followed by brief heating on a hot plate (120 °C, 10-15 s). Structural assignments were made with additional information from qCOSY, qHSQC, and qHMBC experiments.

Chemical Materials

Commercial solvents and reagents were used as received with the following exceptions. Dichloromethane, diethyl ether (ether) and tetrahydrofuran were purified according to the method of Pangborn et al [2]. Triethylamine was distilled from calcium hydride immediately before use. 5-Chloro-2-(methylsulfonyl)-4-nitrobenzamide (**S3**) and 5-chloro-2,4-dinitrobenzamide (**S13**) were prepared according to the published procedures [3,4].

Chemical Instrumentation

Proton nuclear magnetic resonance spectra ($^1$H NMR) were recorded at 400, 500, or 600 megahertz (MHz) at 23 °C. Chemical shifts are expressed in parts per million (ppm, δ scale) downfield from tetramethylsilane and are referenced to residual protium in the NMR solvent ($CD_3S(O)CD_2H$ δ 2.50). Data are represented as follows: chemical shift, multiplicity (s = singlet, d = doublet, t = triplet, q = quarter, m = multiplet and/or multiple resonances), coupling constant in Hertz (Hz), assignment, and integration. Proton-decoupled carbon nuclear magnetic resonance spectra ($^{13}$C{H} NMR) were recorded at 100 or 126 or 151 MHz at 23 °C, unless otherwise noted. Fluorine nuclear magnetic resonance spectra were recorded at 376 or 470 MHz at 23 °C, unless otherwise noted. Chemical shifts are expressed in parts per million (ppm, δ scale) downfield from tetramethylsilane and are referenced to the carbon resonances of the solvent (DMSO-$d_6$, δ 39.5). Herteronuclear single quantum coherence (HSQC) spectra were recorded at 400, 500, or 600 MHz at 23 °C, unless otherwise noted and heteronuclear multiple bond correlation (HMBC) spectra were recorded at 400, 500 or 600 MHz at 23 °C, unless otherwise noted. Proton-proton correlation spectroscopy ($^1$H-$^1$H COSY) spectra were recorded at 600 MHz at 23 °C, unless otherwise noted. High resolution mass spectra (HRMS) were obtained on a Water UPLC/HRMS instrument equipped with a dual API/ESI high-resolution mass spectrometry detector and photodiode array detector. Unless otherwise noted, samples were eluted over a reverse-phase $C_{18}$ column (1.7 $\mu$m particle size, 2.1 x 50 mm) with a linear gradient of 5% acetonitrile-water containing 0.1% formic acid → 95% acetonitrile-water containing 0.1% formic acid for 1 min, at a flow rate of 600 $\mu$L/min.

**Synthetic Procedures.**

*Synthesis of 1-(2,4-dinitrophenyl)aziridine (**1**).*

*Part 1: Synthesis of (chloroethyl)aniline* **S1**.

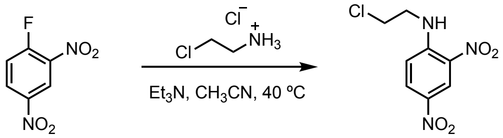


**S1**

Triethylamine (3.00 mL, 21.5 mmol, 2.00 equiv) was added slowly via syringe to a stirred solution of 2-chloroethylamine hydrochloride (2.49 g, 21.9 mmol, 2.00 equiv) and 1-fluoro-2,4-dinitrobenzene (2.00 g, 10.8 mmol, 1 equiv) in anhydrous acetonitrile (54 mL) at 23 °C. The reaction vessel was placed in an oil bath that had been preheated to 40 °C. The reaction mixture was stirred and heated for 2 h at 40 °C. The reaction vessel was then removed from the oil bath, and the product mixture was allowed to cool to 23 °C over 15 min. The cooled product mixture was diluted with ethyl acetate (150 mL). The diluted product mixture was washed sequentially with aqueous hydrogen chloride solution (1M) (1 × 100 mL), saturated aqueous sodium bicarbonate solution (1 × 150 mL) and saturated aqueous sodium chloride solution (2 × 100 mL). The washed organic layer was dried over sodium sulfate. The dried solution was filtered and the filtrate was concentrated. The (chloroethyl)aniline **S1** obtained in this way was unstable toward purification by flash-column chromatography and was used directly in the following step.

*Part 2: Synthesis of 1-(2,4-dinitrophenyl)aziridine (**1**).*

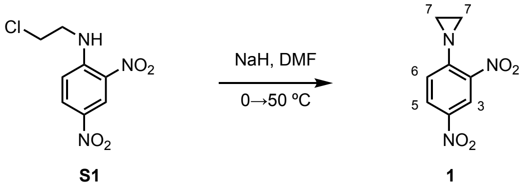


Sodium hydride (60% dispersion in mineral oil, 860 mg, 21.5 mmol, 2.00 equiv) was added portionwise to a stirred solution of the unpurified product obtained in the preceding step (nominally 10.8 mmol, 1 equiv) in anhydrous dimethylformamide (43 mL) at 0 °C. Upon completion of the addition, the cooling bath was removed, and the reaction vessel was placed in an oil bath that had been preheated to 50 °C. The reaction mixture was stirred and heated for 8 h at 50 °C. The reaction vessel was then removed from the oil bath, and the product mixture was allowed to cool to 23 °C over 15 min. The cooled product mixture was diluted with ethyl acetate (150 mL). The diluted product mixture was washed sequentially with saturated aqueous sodium bicarbonate solution (2 × 150 mL) and saturated aqueous sodium chloride solution (2 × 100 mL). The washed organic layer was dried over sodium sulfate. The dried solution was filtered and the filtrate was concentrated. The residue obtained was purified by flash-column chromatography (eluting with 50% ethyl acetate–hexanes) to provide 1-(2,4-dinitrophenyl)aziridine (**1**) as a yellow powder (771 mg, 3.69 mmol, 34% yield over 2 steps).

$R_f$ = 0.50 (50% ethyl acetate–hexanes; UV, CAM). $^1$H NMR (400 MHz, DMSO-$d_6$) $\delta$ 8.74 (d, $J$ = 2.6 Hz, $H_3$, 1H), 8.38 (dd, $J$ = 9.1, 2.0 Hz, $H_5$, 1H), 7.49 (d, $J$ = 9.1 Hz, $H_6$, 1H), 2.46 (s, $H_7$, 4H). $^{13}$C NMR (100 MHz, DMSO-$d_6$) $\delta$ 154.3 (C), 140.44 (C), 140.36 (C), 128.5 (CH), 125.1 (CH), 121.6 (CH), 29.8 (2 × $CH_2$). HRMS-ESI ($m/z$) $[M+H]^+$ calculated for $[C_8H_8N_3O_4]^+$ 210.0509, found 210.0496.

*Synthesis of 5-(aziridin-1-yl)-2,4-dinitroaniline (**2**).*

*Part 1: Synthesis of the (chloroethyl)aniline* **S2**.

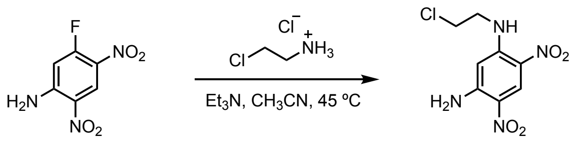

**S2**

Triethylamine (2.77 mL, 19.9 mmol, 2.00 equiv) was added slowly via syringe to a stirred solution of 2-chloroethylamine hydrochloride (2.31 g, 19.9 mmol, 2.00 equiv) and 5-fluoro-2,4-dinitroaniline (2.00 g, 9.95 mmol, 1 equiv) in anhydrous acetonitrile (40 mL) at 23 °C. The reaction vessel was placed in an oil bath that had been preheated to 45 °C. The reaction mixture was stirred and heated for 90 min at 45 °C. The reaction vessel was then removed from the oil bath, and the product mixture was allowed to cool to 23 °C over 15 min. The cooled product mixture was diluted with ethyl acetate (200 mL). The diluted reaction mixture was washed sequentially with saturated aqueous sodium bicarbonate solution (2 × 200 mL) and saturated aqueous sodium chloride solution (2 × 100 mL). The washed organic layer was dried over sodium sulfate. The dried solution was filtered and the filtrate was concentrated. The (chloroethyl)aniline **S2** obtained in this way was unstable toward purification by flash-column chromatography and was used directly in the following step.

*Part 2: Synthesis of 5-(aziridin-1-yl)-2,4-dinitroaniline (**2**).*

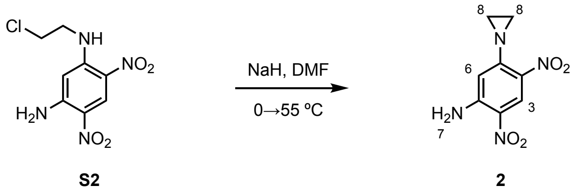

Sodium hydride (60% dispersion in mineral oil, 796 mg, 19.9 mmol, 2.00 equiv) was added portionwise to a stirred solution of the unpurified product obtained in the preceding step (nominally 9.95 mmol, 1 equiv) in anhydrous dimethylformamide (99 mL) at 0 °C. Upon completion of the addition, the cooling bath was removed, and the reaction vessel was placed in an oil bath that had been preheated to 55 °C. The reaction mixture was stirred and heated for 3 h at 55 °C. The reaction vessel was then removed from the oil bath, and the product mixture was allowed to cool to 23 °C over 15 min. The cooled product mixture was diluted with ethyl acetate (150 mL). The diluted product mixture was washed sequentially with saturated aqueous sodium bicarbonate solution (2 × 150 mL) and saturated aqueous sodium chloride solution (3 × 100 mL). The washed organic layer was dried over sodium sulfate. The dried solution was filtered and the filtrate was concentrated. The residue obtained was purified by flash-column chromatography (eluting with 90% ethyl acetate–hexanes) to provide 5-(aziridin-1-yl)-2,4-dinitroaniline (**2**) as a yellow powder (955 mg, 4.26 mmol, 43% yield over 2 steps).

$R_f$ = 0.65 (80% ethyl acetate–hexanes; UV, CAM). $^1$H NMR (400 MHz, DMSO-$d_6$) $\delta$ 8.78 (s, $H_3$, 1H), 7.98 (s, $H_7$, 2H), 6.57 (s, $H_6$, 1H), 2.33 (s, $H_8$, 4H). $^{13}$C NMR (100 MHz, DMSO-$d_6$) $\delta$ 154.7 (C), 149.3 (C), 130.6 (C), 126.7 (CH), 125.1 (C), 109.0 (CH), 29.8 (2 × $CH_2$). HRMS-ESI (*m/z*) $[M+H]^+$ calculated for $[C_8H_9N_4O_4]^+$ 225.0618, found 225.0623.

*Synthesis of N-(5-(aziridin-1-yl)-2,4-dinitrophenyl)acetamide (**3**).*

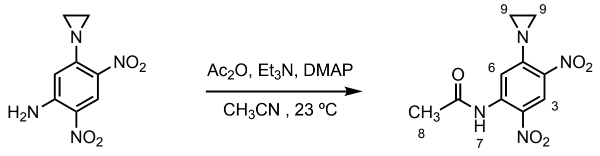


**2** **3**

Triethylamine (248 $\mu$L, 1.78 mmol, 2.00 equiv) was added slowly via syringe to a stirred solution of 5-(aziridin-1-yl)-2,4-dinitroaniline (**2**; 200 mg, 892 $\mu$mol, 1 equiv) and acetic anhydride (168 $\mu$L, 1.78 mmol, 2.00 equiv) in anhydrous acetonitrile (8.9 mL) at 23 °C. Upon completion of the addition, the reaction mixture was stirred for 5 min at 23 °C. After stirring for 5 min at 23 °C, 4-dimethylaminopyridine (10.9 mg, 89.2 $\mu$mol, 0.100 equiv) was added portionwise to the stirred solution. The reaction mixture was stirred for 45 min at 23 °C. The product mixture was diluted with ethyl acetate (25 mL). The diluted product mixture was washed sequentially with saturated aqueous sodium bicarbonate solution (2 × 50 mL) and saturated aqueous sodium chloride solution (1 × 50 mL). The washed organic layer was dried over sodium sulfate. The dried solution was filtered and the filtrate was concentrated. The residue obtained was purified by flash-column chromatography (eluting with 60% ethyl acetate–hexanes) to provide *N*-(5-(aziridin-1-yl)-2,4-dinitrophenyl)acetamide (**3**) as a yellow powder (67.0 mg, 252 $\mu$mol, 28% yield).

$R_f$ = 0.70 (70% ethyl acetate–hexanes; UV, CAM). $^1$H NMR (600 MHz, DMSO-$d_6$) $\delta$ 10.51 (s, $H_7$, 1H), 8.66 (s, $H_3$, 1H), 7.72 (s, $H_6$, 1H), 2.44 (s, $H_9$, 4H), 2.16 (s, $H_8$, 3H). $^{13}$C NMR (151 MHz, DMSO-$d_6$) $\delta$ 169.2 (C), 154.0 (C), 136.9 (C), 135.3 (C), 133.4 (C), 124.4 (CH), 116.8 (CH), 30.0 (2 × $CH_2$), 24.1 ($CH_3$). HRMS-ESI (*m/z*) $[M+H]^+$ calculated for $[C_{10}H_{11}N_4O_5]^+$ 267.0724, found 267.0731.

*Synthesis of 5-(aziridin-1-yl)-2-(methylsulfonyl)-4-nitrobenzamide (**4**).*

*Part 1: Synthesis of the (chloroethyl)aniline* **S4**.

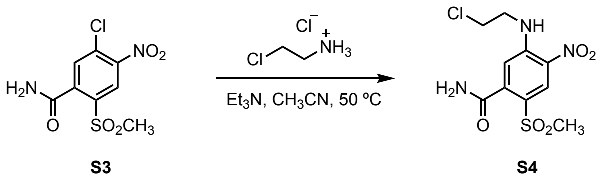

Triethylamine (873 $\mu$L, 6.28 mmol, 5.00 equiv) was added slowly via syringe to a stirred solution of 2-chloroethylamine hydrochloride (728 mg, 6.28 mmol, 5.00 equiv) and 5-chloro-2-(methylsulfonyl)-4-nitrobenzamide (**S3**; 350 mg, 1.26 mmol, 1 equiv) in anhydrous acetonitrile (5.0 mL) at 23 °C. The reaction vessel was placed in an oil bath that had been preheated to 50 °C. The reaction mixture was stirred and heated for 2 h at 50 °C. The reaction vessel was then removed from the oil bath, and the product mixture was allowed to cool to 23 °C over 15 min. The cooled product mixture was diluted with ethyl acetate (100 mL). The diluted product mixture was washed sequentially with saturated aqueous sodium bicarbonate solution (1 × 100 mL) and saturated aqueous sodium chloride solution (2 × 75 mL). The washed organic layer was dried over sodium sulfate. The dried solution was filtered and the filtrate was concentrated. The (chloroethyl)aniline **S4** obtained in this way was unstable toward purification by flash-column chromatography and was used directly in the following step.

*Part 2: Synthesis of 5-(aziridin-1-yl)-2-(methylsulfonyl)-4-nitrobenzamide (***4***).*

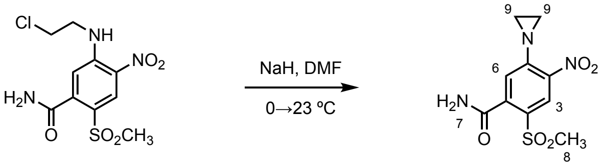


**S4** **4**

Sodium hydride (60% dispersion in mineral oil, 101 mg, 2.52 mmol, 2.00 equiv) was added portionwise to a stirred solution of the unpurified product obtained in the preceding step (nominally 1.26 mmol, 1 equiv) in anhydrous dimethylformamide (13 mL) at 0 °C. Upon completion of the addition, the cooling bath was removed, and the reaction mixture was allowed to warm to 23 °C over 10 min. The warmed reaction mixture was stirred for 3 h at 23 °C. The product mixture was diluted with ethyl acetate (75 mL). The diluted product mixture was washed sequentially with saturated aqueous sodium bicarbonate solution (2 × 100 mL) and saturated aqueous sodium chloride solution (2 × 75 mL). The washed organic layer was dried over sodium sulfate. The dried solution was filtered and the filtrate was concentrated. The residue obtained was purified by flash-column chromatography (eluting with 85% ethyl acetate–hexanes) to provide 5-(aziridin-1-yl)-2-(methylsulfonyl)-4-nitrobenzamide (**4**) as a yellow powder (200 mg, 701 $\mu$mol, 56% yield over 2 steps).

$^{1}$H NMR spectroscopic data for 5-(aziridin-1-yl)-2-(methylsulfonyl)-4-nitrobenzamide (**4**) obtained in this way were identical to those previously reported [3].

$R_f$ = 0.35 (90% ethyl acetate–hexanes; UV, CAM). $^{1}$H NMR (400 MHz, DMSO-$d_6$) $\delta$ 8.46 (s, $H_3$, 1H), 8.23 (s, $H_7$, 1H), 7.89 (s, $H_7$, 1H), 7.43 (s, $H_6$, 1H), 3.41 (s, $H_8$, 3H), 2.47 (s, $H_9$, 4H). $^{13}$C NMR (100 MHz, DMSO-$d_6$) $\delta$ 167.6 (C), 153.3 (C), 142.0 (C), 139.8 (C), 130.6 (C), 127.8 (CH), 124.8 (CH), 44.9 ($CH_3$), 30.0 (2 × $CH_2$). HRMS-ESI (*m/z*) $[M+H]^+$ calculated for $[C_{10}H_{12}N_3O_5S]^+$ 286.0492, found 286.0499.

*Synthesis of 5-chloro-4-nitro-2-(trifluoromethyl)benzoic acid (**S5**).*

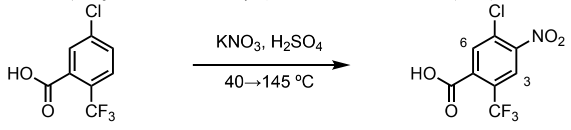


**S5**

Potassium nitrate (4.00 g, 39.6 mmol, 1.19 equiv) was added portionwise over the course of 30 min to a stirred solution of 2-trifluoromethyl-5-chlorobenzoic acid (7.50 g, 33.4 mmol, 1 equiv) in sulfuric acid (75 mL) that had been preheated in an oil bath to 40 °C. Upon completion of the addition, the reaction vessel was brought to 100 °C and additional potassium nitrate (7.00 g, 69.2 mmol, 2.07 eq., 3.26 equiv total) was added portionwise over the course of 20 min. The reaction vessel was brought to 145 °C. The reaction mixture was stirred and heated for 15 min at 145 °C. The reaction vessel was then removed from the oil bath, and the product mixture was allowed to cool to 23 °C over 30 min. The cooled product mixture was poured directly into ice (500 g) and stirred for an additional 30 min. The product was collected by vacuum filtration and washed with water (250 mL). The product was dried on high-vacuum to provide 5-chloro-4-nitro-2-(trifluoromethyl)benzoic acid (**S5**) as a white powder (4.50 g, 16.7 mmol, 50% yield).

$R_f$ = 0.40 (30% methanol–dichloromethane; UV, CAM). $^1$H NMR (400 MHz, DMSO-$d_6$) $\delta$ 8.57 (s, $H_3$, 1H), 8.26 (s, $H_6$, 1H). $^{13}$C NMR (100 MHz, DMSO-$d_6$) $\delta$ 165.3 (C), 148.1 (C), 136.9 (C), 133.0 (CH), 130.0 (C), 126.2 (C) (q, $^2J_{C\text{-}F}$ = 34.0 Hz), 124.7 (CH) (q, $^3J_{C\text{-}F}$ = 5.0 Hz), 122.1 (C) (q, $^1J_{C\text{-}F}$ = 273.0 Hz). $^{19}$F NMR (376 MHz, DMSO-$d_6$) $\delta$ – 58.7 (s). HRMS-ESI (*m/z*) [M-H]$^+$ calculated for $[C_8H_2ClF_3NO_4]^-$ 267.9619/269.9589, found 267.9592/269.9562.

*Synthesis of 5-chloro-4-nitro-2-(trifluoromethyl)benzamide (**S6**). JH-134*

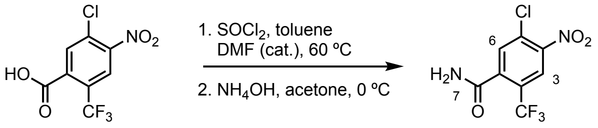


**S5** **S6**

Thionyl chloride (3.65 mL, 50.1 mmol, 3.00 equiv) was added slowly via syringe to a stirred solution of dimethylformamide (129 $\mu$L, 1.67 mmol, 0.100 equiv) and 5-chloro-4-nitro-2-(trifluoromethyl)benzoic acid (**S5**; 4.50 g, 16.7 mmol, 1 equiv) in toluene (67 mL) at 23 °C. The reaction vessel was placed in an oil bath that had been preheated to 60 °C. The reaction mixture was stirred and heated for 6 h at 60 °C. The reaction vessel was then removed from the oil bath, and the product mixture was allowed to cool to 23 °C over 20 min. The cooled product mixture was concentrated. The unpurified acid chloride obtained in this way was used directly in the next step without further purification.

The unpurified acid chloride obtained in the preceding step (nominally 16.7 mmol, 1 equiv) was added slowly via syringe as a solution in acetone (40 mL) to a stirred solution of ammonium hydroxide (28–30% $NH_3$, 20 mL) at 0 °C. The reaction mixture was stirred at 0 °C for 30 min. The reaction mixture was then poured into ice water (500 g) and stirred for an additional 45 min. The product was collected by vacuum filtration and washed with water (200 mL). The product was dried on high-vacuum to provide 5-chloro-4-nitro-2-(trifluoromethyl)benzamide (**S6**) as an off-yellow powder (3.10 g, 11.5 mmol, 69% yield over 2 steps).

$R_f$ = 0.25 (70% ethyl acetate–hexanes; UV, CAM). $^1$H NMR (500 MHz, DMSO-$d_6$) $\delta$ 8.54 (s, $H_3$, 1H), 8.19 (s, $H_7$, 1H), 8.06 (s, $H_6$, 1H), 7.97 (s, $H_7$, 1H). $^{13}$C NMR (126 MHz, DMSO-$d_6$) $\delta$ 165.8 (C), 147.4 (C), 140.9 (C), 131.7 (CH), 129.4 (C), 125.6 (C) (q, $^2J_{C\text{-}F}$ = 32.8 Hz), 124.2 (CH) (q, $^3J_{C\text{-}F}$ = 5.0 Hz), 122.2 (C) (q, $^1J_{C\text{-}F}$ = 274.7 Hz). $^{19}$F NMR (470 MHz, DMSO-$d_6$) $\delta$ – 58.6 (s). HRMS-ESI ($m/z$) $[M+H]^+$ calculated for $[C_8H_5ClF_3N_2O_3]^+$ 268.9935/270.9906, found 268.9941/270.9914.

*Synthesis of 5-(aziridin-1-yl)-4-nitro-2-(trifluoromethyl)benzamide (**5**).*

*Part 1: Synthesis of (chloroethyl)aniline* **S7**:

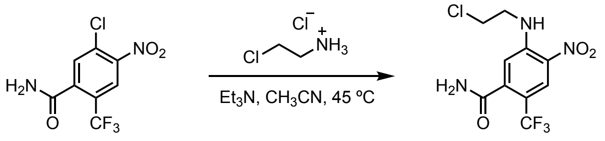


**S6** **S7**

Triethylamine (3.11 mL, 22.3 mmol, 6.00 equiv) was added slowly via syringe to a stirred solution of 2-chloroethylamine hydrochloride (2.59 g, 22.3 mmol, 6.00 equiv) and 5-chloro-4-nitro-2-(trifluoromethyl)benzamide (**S6**; 1.00 g, 3.72 mmol, 1 equiv) in anhydrous acetonitrile (15 mL) at 23 °C. The reaction vessel was placed in an oil bath that had been preheated to 45 °C. The reaction mixture was stirred and heated for 8 h at 45 °C. The reaction vessel was then removed from the oil bath, and the product mixture was allowed to cool to 23 °C over 15 min. The cooled product mixture was diluted with ethyl acetate (100 mL). The diluted product mixture was washed sequentially with saturated aqueous sodium bicarbonate solution (1 × 150 mL) and saturated aqueous sodium chloride solution (2 × 100 mL). The washed organic layer was dried over sodium sulfate. The dried solution was filtered and the filtrate was concentrated. The (chloroethyl)aniline **S7** obtained in this way was unstable toward purification by flash-column chromatography and was used directly in the following step.

*Step 2: Synthesis of 5-(aziridin-1-yl)-4-nitro-2-(trifluoromethyl)benzamide (**5**).*

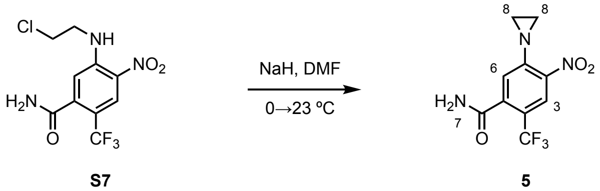


Sodium hydride (60% dispersion in mineral oil, 298 mg, 7.45 mmol, 2.00 equiv) was added portionwise to a stirred solution of the unpurified product obtained in the preceding step (nominally 3.72 mmol, 1 equiv) in anhydrous dimethylformamide (37 mL) at 0 °C. Upon completion of the addition, the cooling bath was removed, and the reaction mixture was allowed to warm to 23 °C. The warmed reaction mixture was stirred for 2 h at 23 °C. The product mixture was diluted with ethyl acetate (150 mL). The diluted product mixture was washed sequentially with saturated aqueous sodium bicarbonate solution (2 × 100 mL) and saturated aqueous sodium chloride solution (2 × 100 mL). The washed organic layer was dried over sodium sulfate. The dried solution was filtered and the filtrate was concentrated. The residue obtained was purified by flash-column chromatography (eluting with 90% ethyl acetate–hexanes) to provide 5-(aziridin-1-yl)-4-nitro-2-(trifluoromethyl)benzamide (**5**) as a yellow powder (390 mg, 1.42 mmol, 38% yield over 2 steps).

$R_f$ = 0.50 (90% ethyl acetate–hexanes; UV, CAM). $^1$H NMR (600 MHz, DMSO-$d_6$) $\delta$ 8.28 (s, $H_3$, 1H), 8.11 (s, $H_7$, 1H), 7.83 (s, $H_7$, 1H), 7.42 (s, $H_6$, 1H), 2.41 (s, $H_8$, 4H). $^{13}$C NMR (151 MHz, DMSO-$d_6$) $\delta$ 167.1 (C), 151.9 (C), 141.1 (C), 140.6 (C), 124.3 (CH), 124.2 (CH) (q, $^3J_{C\text{-}F}$ = 4.5 Hz), 122.9 (C) (q, $^1J_{C\text{-}F}$ = 273.3 Hz), 118.8 (C) (q, $^2J_{C\text{-}F}$ = 33.2 Hz), 29.6 (2 × $CH_2$). $^{19}$F (376 MHz, DMSO-$d_6$) $\delta$ – 57.7 (s). HRMS-ESI (*m/z*) [M+H]$^+$ calculated for $[C_{10}H_9F_3N_3O_3]^+$ 276.0591, found 276.0599.

*Synthesis of 3-(aziridin-1-yl)-4-nitrobenzamide (**6**).*

*Step 1: Synthesis of the (chloroethyl)aniline* **S8**.

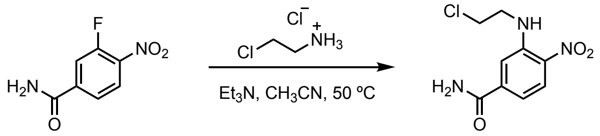

**S8**

Triethylamine (1.51 mL, 10.9 mmol, 6.00 equiv) was added slowly via syringe to a stirred solution of 2-chloroethylamine hydrochloride (1.26 g, 10.9 mmol, 6.00 equiv) and 3-fluoro-4-nitrobenzamide (333 mg, 1.81 mmol, 1 equiv) in anhydrous acetonitrile (7 mL) at 23 °C. The reaction vessel was placed in an oil bath that had been preheated to 50 °C. The reaction mixture was stirred and heated for 16 h at 50 °C. The reaction vessel was then removed from the oil bath, and the product mixture was allowed to cool to 23 °C over 15 min. The cooled product mixture was diluted with ethyl acetate (150 mL). The diluted product mixture was washed sequentially with aqueous hydrogen chloride solution (1M) (1 × 75 mL), saturated aqueous sodium bicarbonate solution (2 × 75 mL) and saturated aqueous sodium chloride solution (2 × 75 mL). The washed organic layer was dried over sodium sulfate. The dried solution was filtered and the filtrate was concentrated. The (chloroethyl)aniline **S8** obtained in this way was unstable toward purification by flash-column chromatography and was used directly in the following step.

*Step 2: Synthesis of 3-(aziridin-1-yl)-4-nitrobenzamide (**6**).*

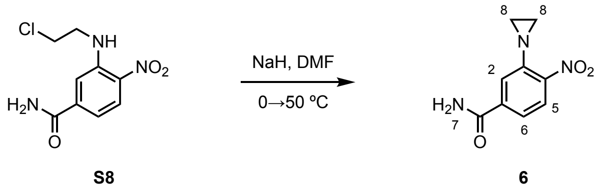

Sodium hydride (60% dispersion in mineral oil, 181 mg, 4.52 mmol, 2.50 equiv) was added portionwise to a stirred solution of the crude residue obtained in the preceding step (nominally 1.81 mmol, 1 equiv) in anhydrous dimethylformamide (7 mL) at 0 °C. Upon completion of the addition, the cooling bath was removed, and the reaction vessel was then placed in an oil bath that had been preheated to 50 °C. The reaction mixture was stirred and heated for 3 h at 50 °C. The reaction vessel was removed from the oil bath, and the product mixture was allowed to cool to 23 °C over 15 min. The cooled product mixture was diluted with ethyl acetate (75 mL). The diluted product mixture was washed sequentially with saturated aqueous sodium bicarbonate solution (2 × 75 mL) and saturated aqueous sodium chloride solution (3 × 50 mL). The washed organic layer was dried over sodium sulfate. The dried solution was filtered and the filtrate was concentrated. The residue obtained was purified by flash-column chromatography (eluting with 10% triethylamine–ethyl acetate) to provide (3-(aziridin-1-yl)-4-nitrobenzamide (**6**) as a yellow powder (145 mg, 700 $\mu$mol, 39% yield over 2 steps).

$^1$H NMR spectroscopic data for 3-(aziridin-1-yl)-4-nitrobenzamide (**6**) obtained in this way were identical to those previously reported [4].

$R_f$ = 0.30 (10% triethylamine–ethyl acetate; UV, CAM). $^1$H NMR (600 MHz, DMSO-$d_6$) $\delta$ 8.20 (s, $H_7$, 1H), 7.99 (d, $J$ = 8.3 Hz, $H_5$, 1H), 7.68 (d, $J$ = 1.7 Hz, $H_2$, 1H), 7.66 (s, $H_7$, 1H), 7.56 (dd, $J$ = 8.5, 1.6 Hz, $H_6$, 1H), 2.30 (s, $H_8$, 4H). $^{13}$C NMR (151 MHz, DMSO-$d_6$) $\delta$ 166.1 (C), 148.7 (C), 143.3 (C), 139.1 (C), 125.3 (CH), 122.9 (CH), 121.0 (CH), 29.2 (2 × $CH_2$). HRMS-ESI (*m/z*) $[M+H]^+$ calculated for $[C_9H_{10}N_3O_3]^+$ 208.0717, found 208.0706.

*Synthesis of 5-chloro-2-nitro-4-(trifluoromethyl)benzoic acid (**S9**).*

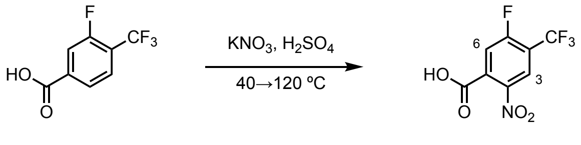


**S9**

Potassium nitrate (7.30 g, 72.1 mmol, 2.50 equiv) was added portionwise over the course of 10 min to a stirred solution of 3-fluoro-4-(trifluoromethyl)benzoic acid (6.00 g, 28.8 mmol, 1 equiv) in sulfuric acid (35 mL) that had been preheated in an oil bath to 40 °C. Upon completion of the addition, the reaction vessel was brought to 120 °C. The reaction mixture was stirred and heated for 60 min at 120 °C. The reaction vessel was then removed from the oil bath, and the product mixture was allowed to cool to 23 °C over 30 min. The cooled product mixture was poured into ice (500 g) and stirred for an additional 30 min. The product was collected by vacuum filtration and washed with water (250 mL). The product was dried on high-vacuum to provide 5-fluoro-2-nitro-4-(trifluoromethyl)benzoic acid (**S9**) as a white powder (3.10 g, 12.3 mmol, 43% yield).

$R_f$ = 0.40 (20% methanol–dichloromethane; UV, CAM). $^{1}$H NMR (600 MHz, DMSO-$d_6$) $\delta$ 8.50 (d, $J$ = 5.9 Hz, $H_3$, 1H), 8.06 (d, $J$ = 9.9 Hz, $H_6$, 1H). $^{13}$C NMR (151 MHz, DMSO-$d_6$) $\delta$ 164.2 (C), 160.6 (C) (d, $^{1}J_{C\text{-}F}$ = 264.3 Hz), 143.5 (C) (d, $^{4}J_{C\text{-}F}$ = 3.0 Hz), 134.9 (C) (d, $^{3}J_{C\text{-}F}$ = 9.0 Hz), 124.6 (CH) (apparent m), 121.2 (C) (q, $^{1}J_{C\text{-}F}$ = 273.3 Hz), 119.1 (C) (qd, $^{2}J_{C\text{-}F}$ = 34.7 Hz, $^{2}J_{C\text{-}F}$ = 13.6 Hz), 119.0 (CH) (d, $^{2}J_{C\text{-}F}$ = 24.2 Hz). $^{19}$F NMR (376 MHz, DMSO-$d_6$) $\delta$ -60.8 (d, $J$ = 12.7 Hz), -106.4 (apparent m). HRMS-ESI (*m/z*) [M-H]$^+$ calculated for [$C_8H_2F_4NO_4$]$^-$ 251.9914, found 251.9888.

*Synthesis of 5-fluoro-2-nitro-4-(trifluoromethyl)benzamide (**S10**).*

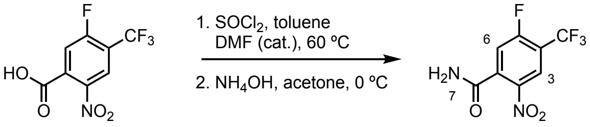


**S9** **S10**

Thionyl chloride (1.66 mL, 22.7 mmol, 2.50 equiv) was added slowly via syringe to a stirred solution of dimethylformamide (70 $\mu$L, 909 $\mu$mol, 0.100 equiv) and 5-fluoro-2-nitro-4-(trifluoromethyl)benzoic acid (**S9**; 2.30 g, 9.09 mmol, 1 equiv) in toluene (45 mL) at 23 °C. The reaction vessel was placed in an oil bath that had been preheated to 60 °C. The reaction mixture was stirred and heated for 5 h at 60 °C. The reaction vessel was then removed from the oil bath, and the product mixture was allowed to cool to 23 °C over 20 min. The cooled product mixture was concentrated. The unpurified acid chloride obtained in this way was used directly in the next step without further purification.

The unpurified acid chloride obtained in the preceding step (nominally 9.09 mmol, 1 equiv) was added slowly via syringe as a solution in acetone (25 mL) to a stirred solution of ammonium hydroxide (28–30% $NH_3$, 15 mL) at 0 °C. The reaction was stirred for 45 min at 0 °C. The product mixture was then poured into ice (500 g) and stirred for an additional 45 min. The product was collected by vacuum filtration and washed with water (200 mL). The product was dried on high-vacuum to provide 5-fluoro-2-nitro-4-(trifluoromethyl)benzamide (**S10**) as a white powder (1.57 g, 6.23 mmol, 69% yield over 2 steps).

$R_f$ = 0.35 (50% ethyl acetate–hexanes; UV, CAM). $^1$H NMR (600 MHz, DMSO-$d_6$) $\delta$ 8.48 (d, $J$ = 6.0 Hz, $H_3$, 1H), 8.29 (s, $H_7$, 1H), 8.01 (s, $H_7$, 1H), 7.92 (d, $J$ = 10.0 Hz, $H_6$, 1H). $^{13}$C NMR (151 MHz, DMSO-$d_6$) $\delta$ 164.7 (C), 160.5 (d, $^1J_{C\text{-}F}$ = 261.2 Hz), 142.9 (C) (d, $^4J_{C\text{-}F}$ = 3.0 Hz), 139.4 (C) (d, $^3J_{C\text{-}F}$ = 7.6 Hz), 124.6 (CH) (apparent m), 121.2 (q, $^1J_{C\text{-}F}$ = 273.3 Hz), 118.4 (d, $^2J_{C\text{-}F}$ = 24.2 Hz), 117.8 (qd, $^2J_{C\text{-}F}$ = 34.3 Hz, $^2J_{C\text{-}F}$ = 15.1 Hz). $^{19}$F NMR (470 MHz, DMSO-$d_6$) $\delta$ -60.6 (d, $J$ = 14.1 Hz), -106.9 (apparent m). HRMS-ESI (*m/z*) $[M+H]^+$ calculated for $[C_8H_5F_4N_2O_3]^+$ 253.0231, found 253.0233.

~

*Synthesis of 5-(aziridin-1-yl)-2-nitro-4-(trifluoromethyl)benzamide (7).*

*Step 1: Synthesis of the (chloroethyl)aniline* **S11**:

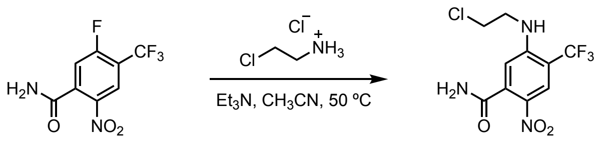


**S10** **S11**

Triethylamine (1.54 mL, 11.1 mmol, 4.00 equiv) was added slowly via syringe to a stirred solution of 2-chloroethylamine hydrochloride (1.29 g, 11.1 mmol, 4.00 equiv) and 5-fluoro-2-nitro-4-(trifluoromethyl)benzamide (**S10**; 700 mg, 2.78 mmol, 1 equiv) in anhydrous acetonitrile (11 mL) at 23 °C. The reaction vessel was placed in an oil bath that had been preheated to 50 °C. The reaction mixture was stirred and heated for 2 h at 50 °C. The reaction vessel was then removed from the oil bath, and the product mixture was allowed to cool to 23 °C over 15 min. The cooled product mixture was diluted with ethyl acetate (100 mL). The diluted product mixture was washed sequentially with saturated aqueous sodium bicarbonate solution (2 × 100 mL) and saturated aqueous sodium chloride solution (2 × 100 mL). The washed organic layer was dried over sodium sulfate. The dried solution was filtered and the filtrate was concentrated. The (chloroethyl)aniline **S11** obtained in this way was unstable toward purification by flash-column chromatography and was used directly in the following step.

*Step 2: Synthesis of 5-(aziridin-1-yl)-2-nitro-4-(trifluoromethyl)benzamide (7).*

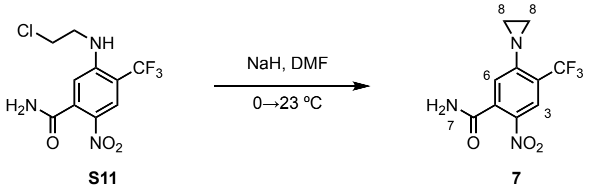


Sodium hydride (60% dispersion in mineral oil, 278 mg, 6.94 mmol, 2.50 equiv) was added portionwise to a stirred solution of the unpurified product obtained in the preceding step (nominally 2.78 mmol, 1 equiv) in anhydrous dimethylformamide (28 mL) at 0 °C. Upon completion of the addition, the cooling bath was removed and the reaction vessel was warmed to 23 °C and stirred for 2 h. The warmed product mixture was diluted with ethyl acetate (100 mL). The diluted product mixture was washed sequentially with saturated aqueous sodium bicarbonate solution (2 × 100 mL) and saturated aqueous sodium chloride solution (2 × 100 mL). The washed organic layer was dried over sodium sulfate. The dried solution was filtered and the filtrate was concentrated. The residue obtained was purified by flash column chromatography on silica (eluting with a mixture of 70:15:5:10 dichloromethane–ethyl acetate–methanol–triethylamine) to provide 5-(aziridin-1-yl)-2-nitro-4-(trifluoromethyl)benzamide (**7**) as a white powder (180 mg, 654 μmol, 24% yield over 2 steps).

$R_f$ = 0.45 (70:15:5:10 dichloromethane–ethyl acetate–methanol–triethylamine; UV, CAM). $^{1}$H NMR (600 MHz, DMSO-$d_6$) δ 8.23 (s, $H_3$, 1H), 8.13 (s, $H_7$, 1H), 7.83 (s, $H_7$, 1H), 7.29 (s, $H_6$, 1H), 2.46 (s, $H_8$, 4H). $^{13}$C NMR (151 MHz, DMSO-$d_6$) δ 166.2 (C), 156.6 (C), 139.5 (C), 138.0 (C), 123.8 (CH) (q, $^{3}J_{C\text{-}F}$ = 4.5 Hz), 122.9 (C) (q, $^{1}J_{C\text{-}F}$ = 273.3 Hz), 122.3 (CH), 120.8 (q, $^{2}J_{C\text{-}F}$ = 31.7 Hz), 29.3 (2 × $CH_2$). $^{19}$F NMR (376 MHz, DMSO-$d_6$) δ -60.4 (s). HRMS-ESI (*m/z*) $[M+H]^+$ calculated for $[C_{10}H_9F_3N_3O_3]^+$ 276.0590, found 276.0588.

*Synthesis of 5-(aziridin-1-yl)-2-nitrobenzamide (**8**).*

*Step 1: Synthesis of the (chloroethyl)aniline* **S12**.

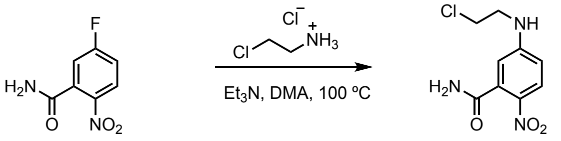

**S12**

Triethylamine (7.55 mL, 54.3 mmol, 5.00 equiv) was added slowly via syringe to a stirred solution of 2-chloroethylamine hydrochloride (6.30 g, 54.3 mmol, 5.00 equiv) and 5-fluoro-2-nitrobenzamide (2.00 g, 10.9 mmol, 1 equiv) in anhydrous dimethylacetamide (27 mL) at 23 °C. The reaction vessel was then placed in an oil bath that had been preheated to 100 °C. The reaction mixture was stirred and heated for 5 h at 100 °C. The reaction vessel was then removed from the oil bath, and the product mixture was allowed to cool to 23 °C over 30 min. The cooled product mixture was diluted with ethyl acetate (100 mL). The diluted product mixture was washed sequentially with aqueous hydrogen chloride solution (1M) (1 × 150 mL), saturated aqueous sodium bicarbonate solution (2 × 150 mL) and saturated aqueous sodium chloride solution (2 × 150 mL). The washed organic layer was dried over sodium sulfate. The dried solution was filtered and the filtrate was concentrated. The (chloroethyl)aniline **S12** obtained in this way was unstable toward purification by flash-column chromatography and was used directly in the following step.

*Step 2: Synthesis of 5-(aziridin-1-yl)-2-nitrobenzamide (**8**).*

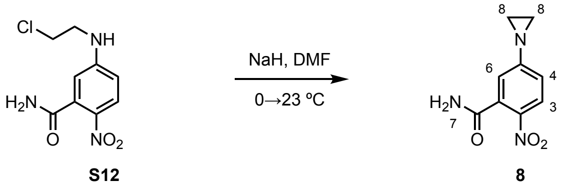

Sodium hydride (60% dispersion in mineral oil, 1.09 g, 27.2 mmol, 2.50 equiv) was added portionwise to a stirred solution of the unpurified product obtained in the preceding step (nominally 10.9 mmol, 1 equiv) in anhydrous dimethylformamide (109 mL) at 0 °C. Upon completion of the addition, the cooling bath was removed, and the reaction mixture was allowed to warm to 23 °C over 10 min. The warmed reaction mixture was then stirred for 1.5 h at 23 °C. The product mixture was diluted with ethyl acetate (250 mL). The diluted product mixture was washed sequentially with saturated aqueous sodium bicarbonate solution (2 × 200 mL) and saturated aqueous sodium chloride solution (2 × 200 mL). The washed organic layer was dried over sodium sulfate. The dried solution was filtered and the filtrate was concentrated. The residue obtained was purified by flash-column chromatography (eluting with 5% methanol–ethyl acetate) to provide 5-(aziridin-1-yl)-2-nitrobenzamide (**8**) as a yellow powder (260 mg, 1.26 mmol, 12% yield over 2 steps).

$^1$H NMR spectroscopic data for 5-(aziridin-1-yl)-2-nitrobenzamide (**8**) obtained in this way were identical to those previously reported [3].

$R_f$ = 0.35 (5% methanol–ethyl acetate; UV, CAM). $^1$H NMR (400 MHz, DMSO-$d_6$) $\delta$ 8.02 (s, $H_7$, 1H), 7.93 (d, $J$ = 8.7 Hz, $H_3$, 1H), 7.62 (s, $H_7$, 1H), 7.17 (dd, $J$ = 8.8, 2.4 Hz, $H_4$, 1H), 7.10 (d, $J$ = 2.3 Hz, $H_6$, 1H), 2.24 (s, $H_8$, 4H). $^{13}$C NMR (100 MHz, DMSO-$d_6$) $\delta$ 167.4 (C), 159.9 (C), 140.4 (C), 134.9 (C), 125.7 (CH), 121.5 (CH), 120.5 (CH), 27.8 (2 × $CH_2$). HRMS-ESI ($m/z$) $[M+H]^+$ calculated for $[C_9H_{10}N_3O_3]^+$ 208.0717, found 208.0717.

*Synthesis of 5-(dimethylamino)-2,4-dinitrobenzamide (**9**).*

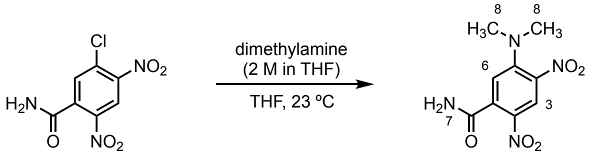


A solution of dimethylamine in tetrahydrofuran (2 M, 8.14 mL, 16.3 mmol, 4.00 equiv) was added slowly via syringe to a stirred solution of 5-chloro-2,4-dinitrobenzamide (**S13**; 1.00 g, 4.07 mmol, 1 equiv) in anhydrous tetrahydrofuran (20 mL) at 23 °C. The reaction mixture was stirred for 2 h at 23 °C. The product mixture was then diluted with ethyl acetate (200 mL). The diluted product mixture was washed sequentially with saturated aqueous sodium bicarbonate solution (2 × 200 mL) and saturated aqueous sodium chloride solution (2 × 200 mL). The washed organic layer was dried over sodium sulfate. The dried solution was filtered, and the filtrate was concentrated to provide 5-(dimethylamino)-2,4-dinitrobenzamide (**9**) as a yellow powder (415 mg, 1.63 mmol, 40% yield).

$R_f$ = 0.10 (100% ethyl acetate; UV, $KMNO_4$). $^{1}$H NMR (600 MHz, DMSO-$d_6$) $\delta$ 8.52 (s, $H_3$, 1H), 8.04 (s, $H_7$, 1H), 7.76 (s, $H_7$, 1H), 7.13 (s, $H_6$, 1H), 3.01 (s, $H_8$, 6H). $^{13}$C NMR (151 MHz, DMSO-$d_6$) $\delta$ 166.7 (C), 147.9 (C), 137.8 (C), 134.0 (C), 133.1 (C), 125.1 (CH), 116.8 (CH), 42.3 (2 × $CH_3$). $[M+H]^+$ calculated for $[C_9H_{11}N_4O_5]^+$ 255.0724, found 255.0722.

*Synthesis of 5-((2-chloroethyl)amino)-2,4-dinitrobenzamide (**10**).*

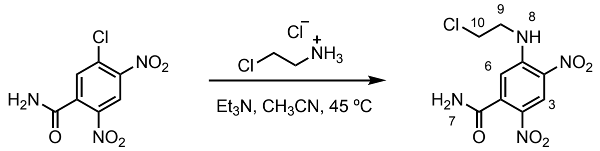


**S13** **10**

Triethylamine (1.13 mL, 8.14 mmol, 2.00 equiv) was added slowly via syringe to a stirred solution of 2-chloroethylamine hydrochloride (945 mg, 8.14 mmol, 2.00 equiv) and 5-chloro-2,4-dinitrobenzamide (**S13**; 1.00 g, 4.07 mmol, 1 equiv) in anhydrous acetonitrile (20 mL) at 23 °C. The reaction vessel was then placed in an oil bath that had been preheated to 45 °C. The reaction mixture was stirred and heated for 2 h at 45 °C. The reaction vessel was then removed from the oil bath, and the product mixture was allowed to cool to 23 °C over 15 min. The cooled product mixture was diluted with ethyl acetate (200 mL). The diluted product mixture was triturated with cold ethyl acetate (75 mL) and the product collected by vacuum filtration. The product obtained was dried on high vacuum to provide 5-((2-chloroethyl)amino-2,4-dinitrobenzamide (**10**) as a yellow powder (365 mg, 1.27 mmol, 31% yield).

Note: Partial decomposition of 5-((2-chloroethyl)amino)-2,4-dinitrobenzamide (**10**) was observed upon purification by flash-column chromatography.

$^{1}$H NMR spectroscopic data for 5-(aziridin-1-yl)-2-(methylsulfonyl)-4-nitrobenzamide (**10**) obtained in this way were identical to those previously reported [5].

$R_f$ = 0.25 (100% ethyl acetate; UV, CAM). $^{1}$H NMR (600 MHz, DMSO-$d_6$) $\delta$ 8.92 (t, $J$ = 5.4 Hz, $H_8$, 1H), 8.77 (s, $H_3$, 1H), 8.12 (s, $H_7$, 1H), 7.81 (s, $H_7$, 1H), 7.20 (s, $H_6$, 1H), 3.93–3.84 (m, $H_9$, $H_{10}$, 4H). $^{13}$C NMR (151 MHz, DMSO-$d_6$) $\delta$ 166.4 (C), 147.1 (C), 140.0 (C), 133.4 (C), 129.3 (C), 124.8 (CH), 114.5 (CH), 44.2 ($CH_2$), 42.8 ($CH_2$). HRMS-ESI (*m/z*) $[M+H]^+$ calculated for $[C_9H_{10}ClN_4O_5]^+$, 289.0334/291.0305; found 289.0334/291.0317.

### Catalog of Nuclear Magnetic Resonance Spectra.

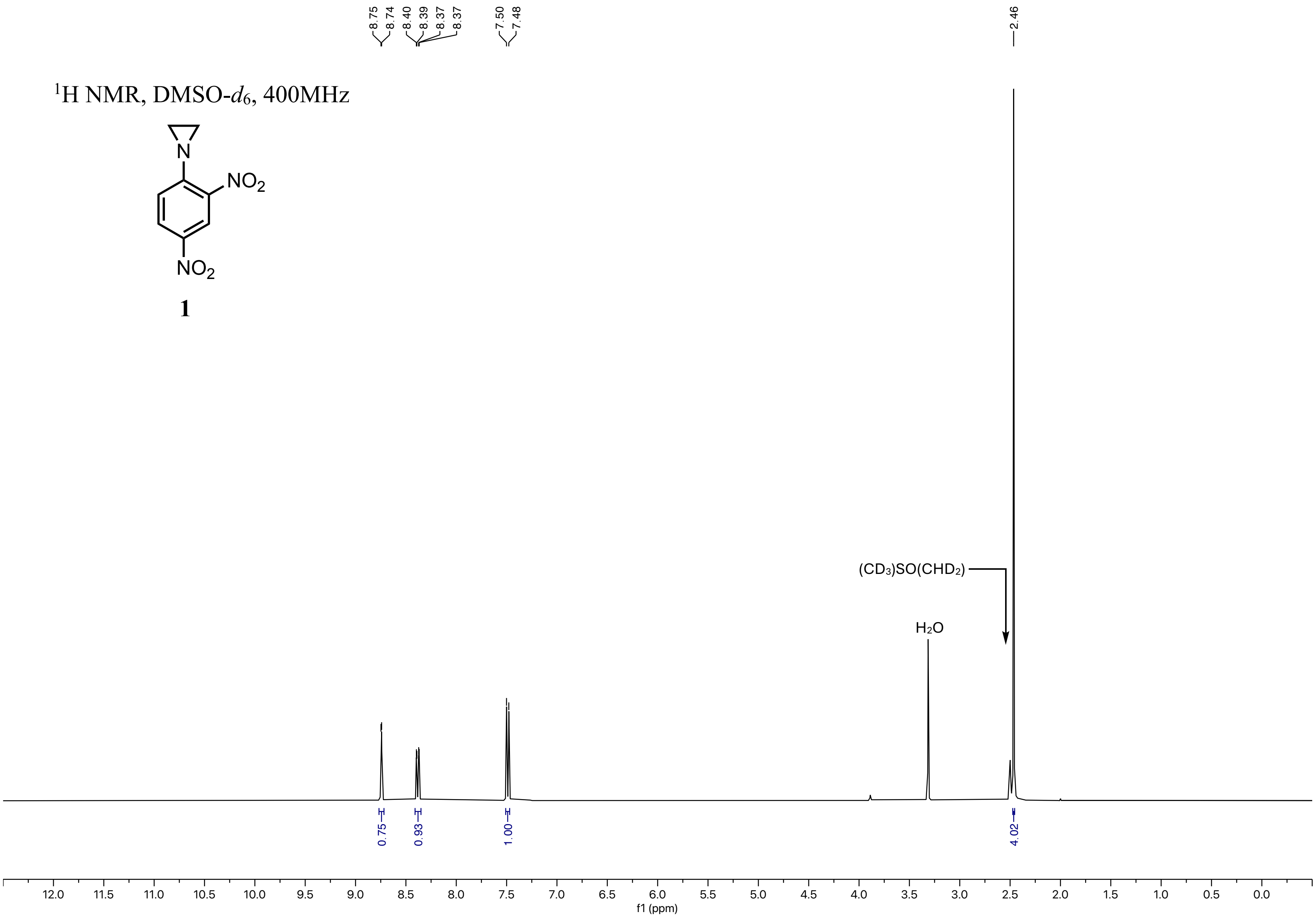

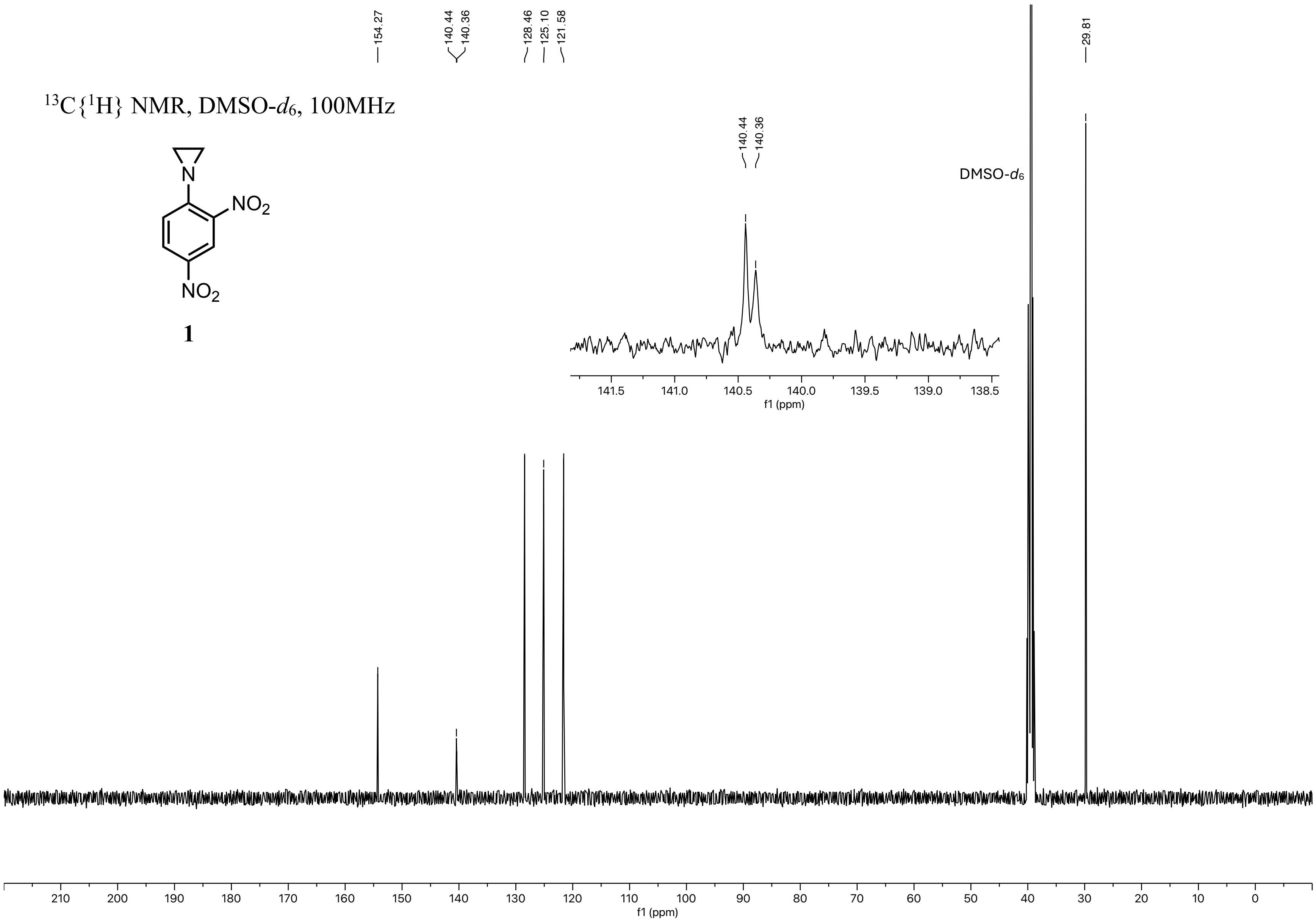

13C{1H} NMR, DMSO-d6, 100MHz
NO2
NO2
N
1
154.27
140.44
140.36
128.46
125.10
121.58
29.81
DMSO-d6
141.5
141.0
140.5
140.0
139.5
139.0
138.5
f1 (ppm)
210
200
190
180
170
160
150
140
130
120
110
100
90
80
70
60
50
40
30
20
10
0
f1 (ppm)

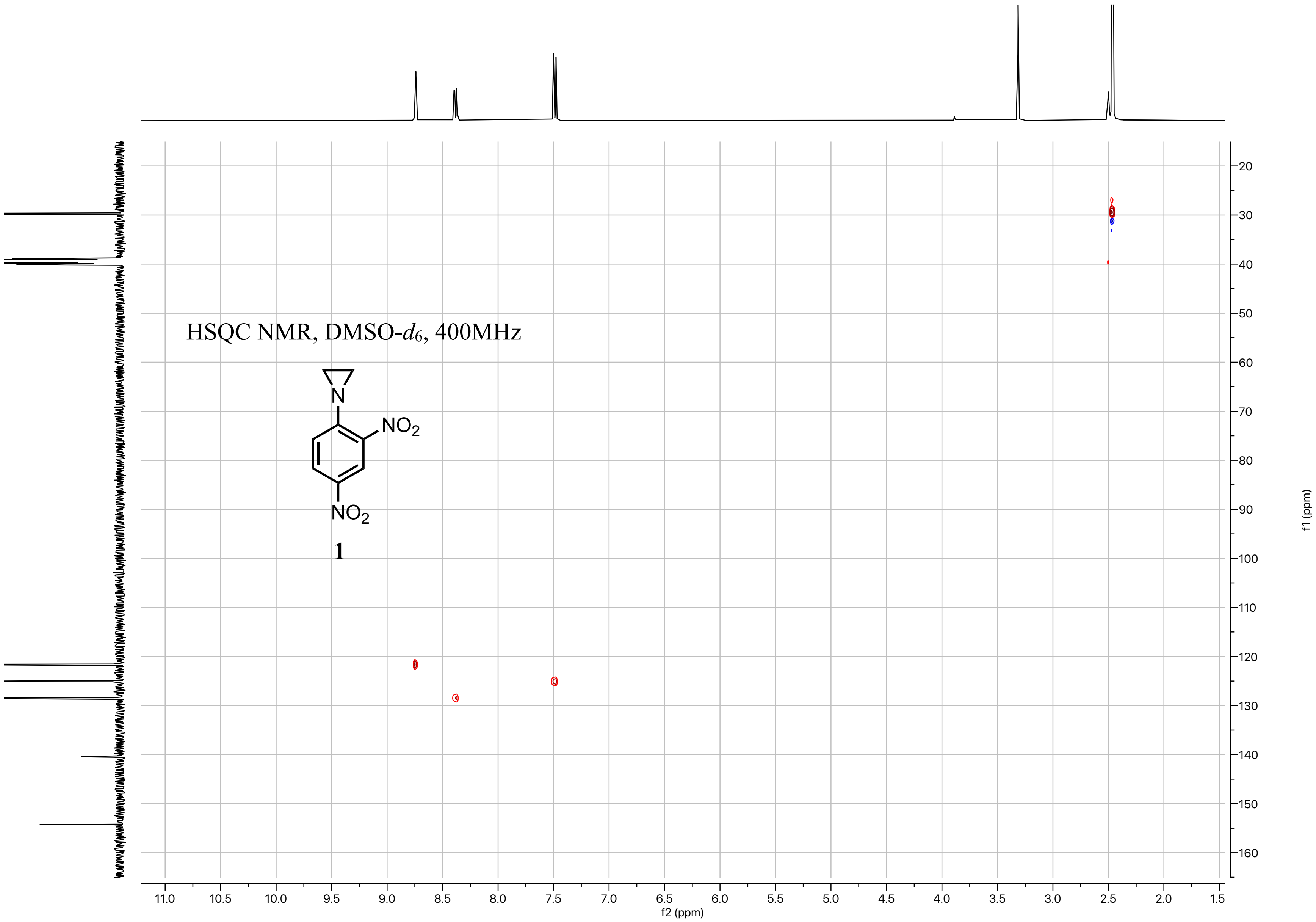

HSQC NMR, DMSO-$d_6$, 400MHz
N
NO2
NO2
1
f2 (ppm)
f1 (ppm)
11.0
10.5
10.0
9.5
9.0
8.5
8.0
7.5
7.0
6.5
6.0
5.5
5.0
4.5
4.0
3.5
3.0
2.5
2.0
1.5
20
30
40
50
60
70
80
90
100
110
120
130
140
150
160

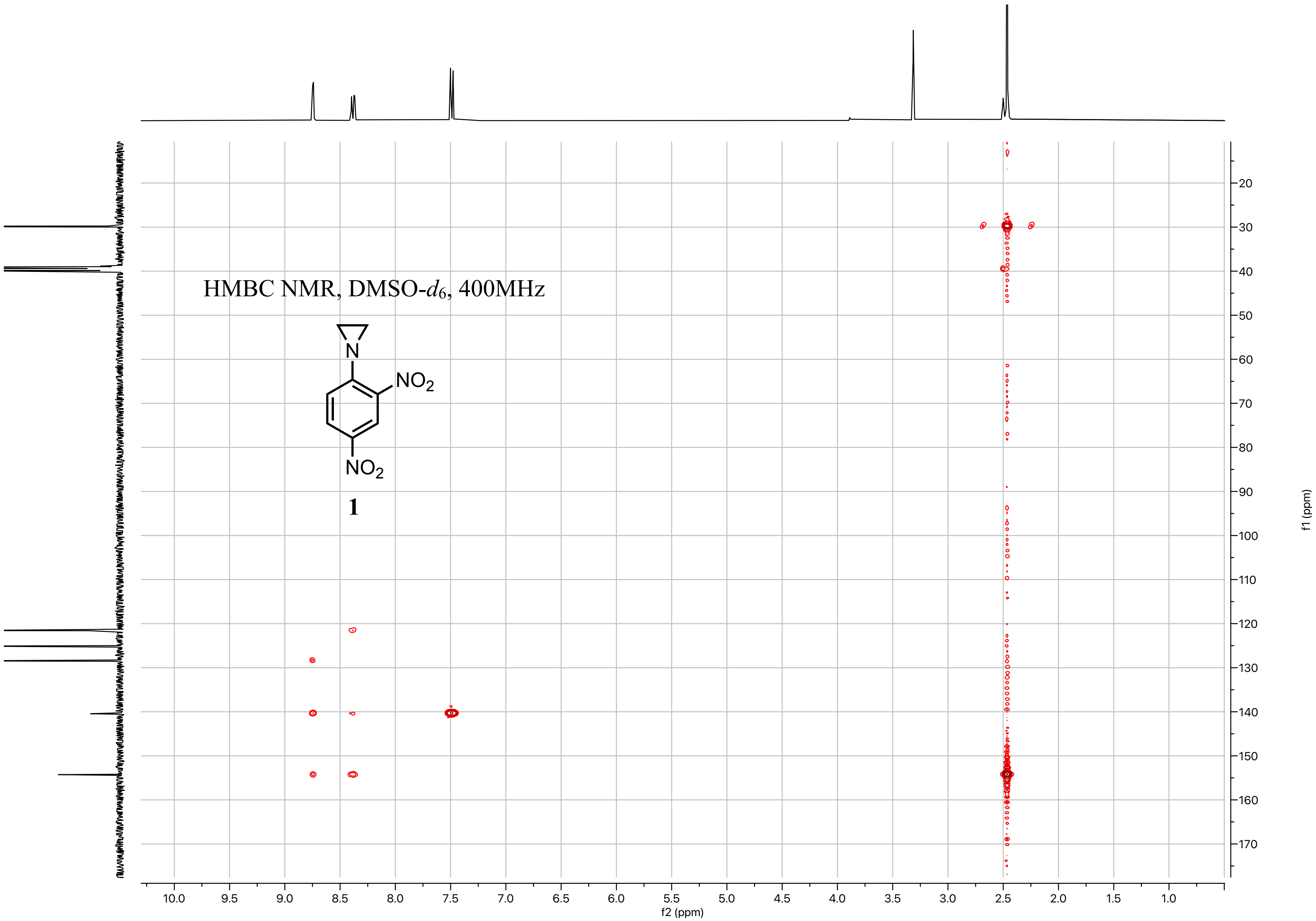
HMBC NMR, DMSO-d6, 400MHz
N
NO2
NO2
1
10.0 9.5 9.0 8.5 8.0 7.5 7.0 6.5 6.0 5.5 5.0 4.5 4.0 3.5 3.0 2.5 2.0 1.5 1.0
f2 (ppm)
20 30 40 50 60 70 80 90 100 110 120 130 140 150 160 170
f1 (ppm)

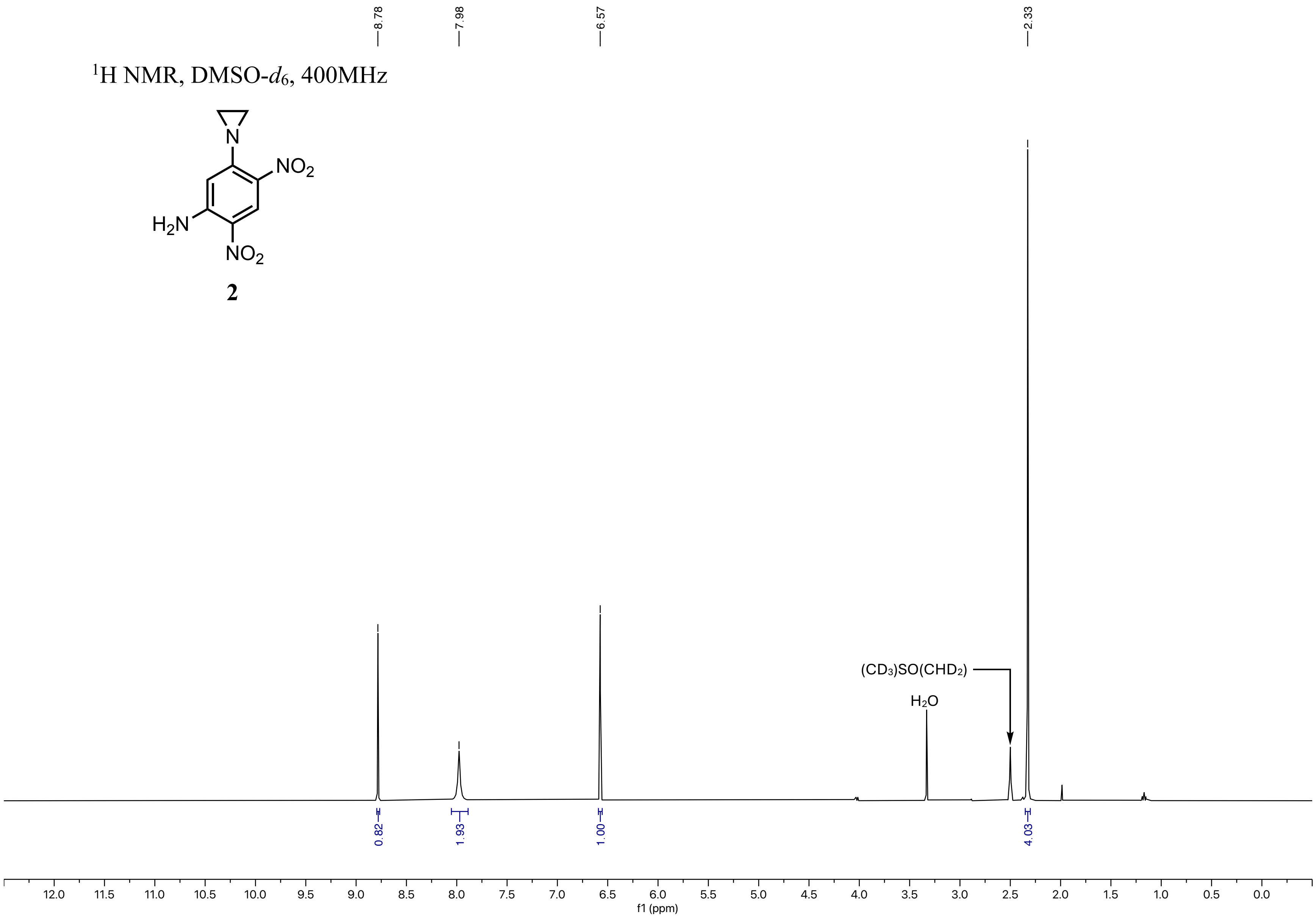

1H NMR, DMSO-d6, 400MHz
NO2
H2N
NO2
N
2
8.78
7.98
6.57
2.33
(CD3)SO(CHD2)
H2O
0.82
1.93
1.00
4.03
12.0 11.5 11.0 10.5 10.0 9.5 9.0 8.5 8.0 7.5 7.0 6.5 6.0 5.5 5.0 4.5 4.0 3.5 3.0 2.5 2.0 1.5 1.0 0.5 0.0
f1 (ppm)

$^{13}C\{^1H\}$ NMR, DMSO-$d_6$, 100MHz

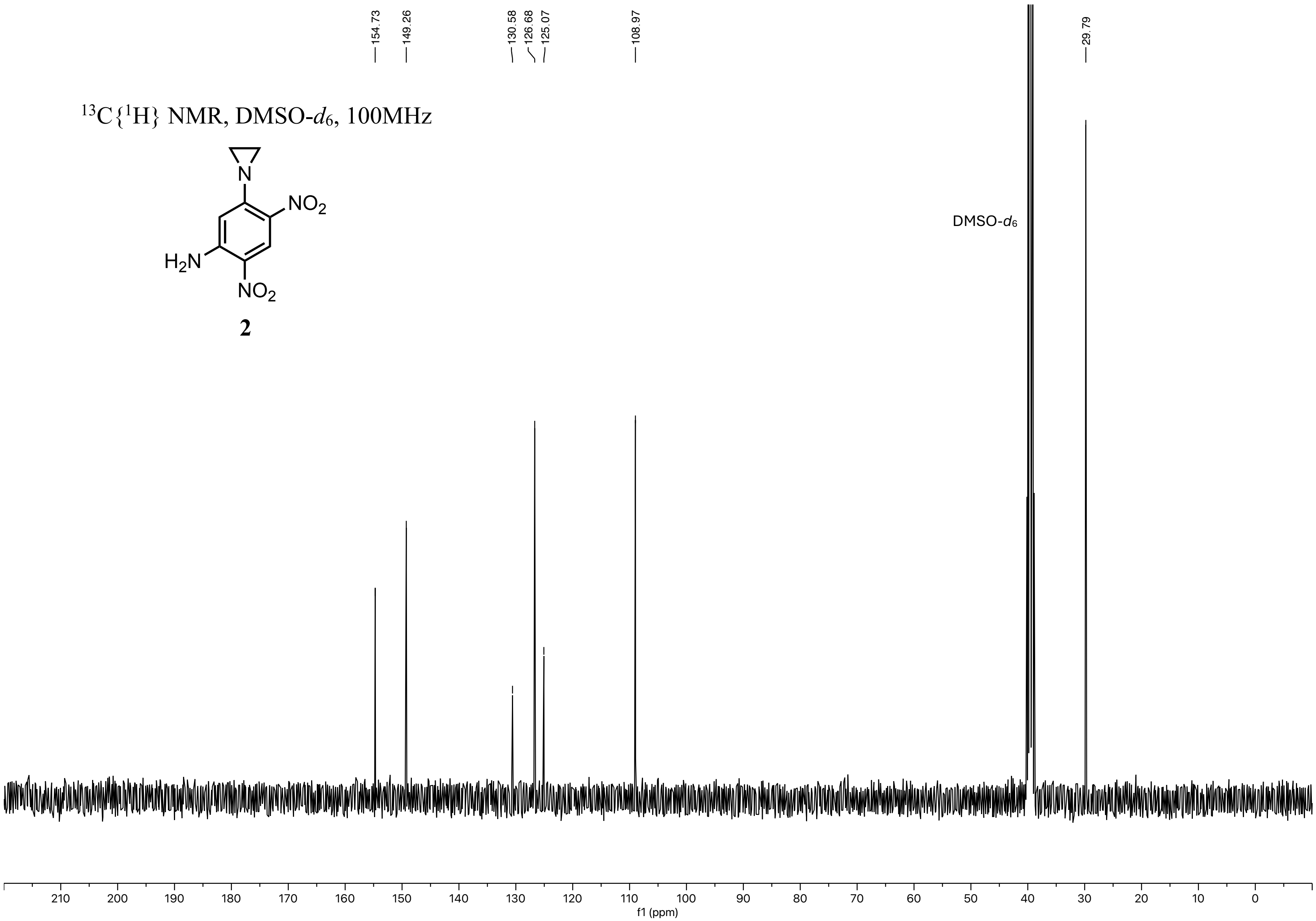


**2**

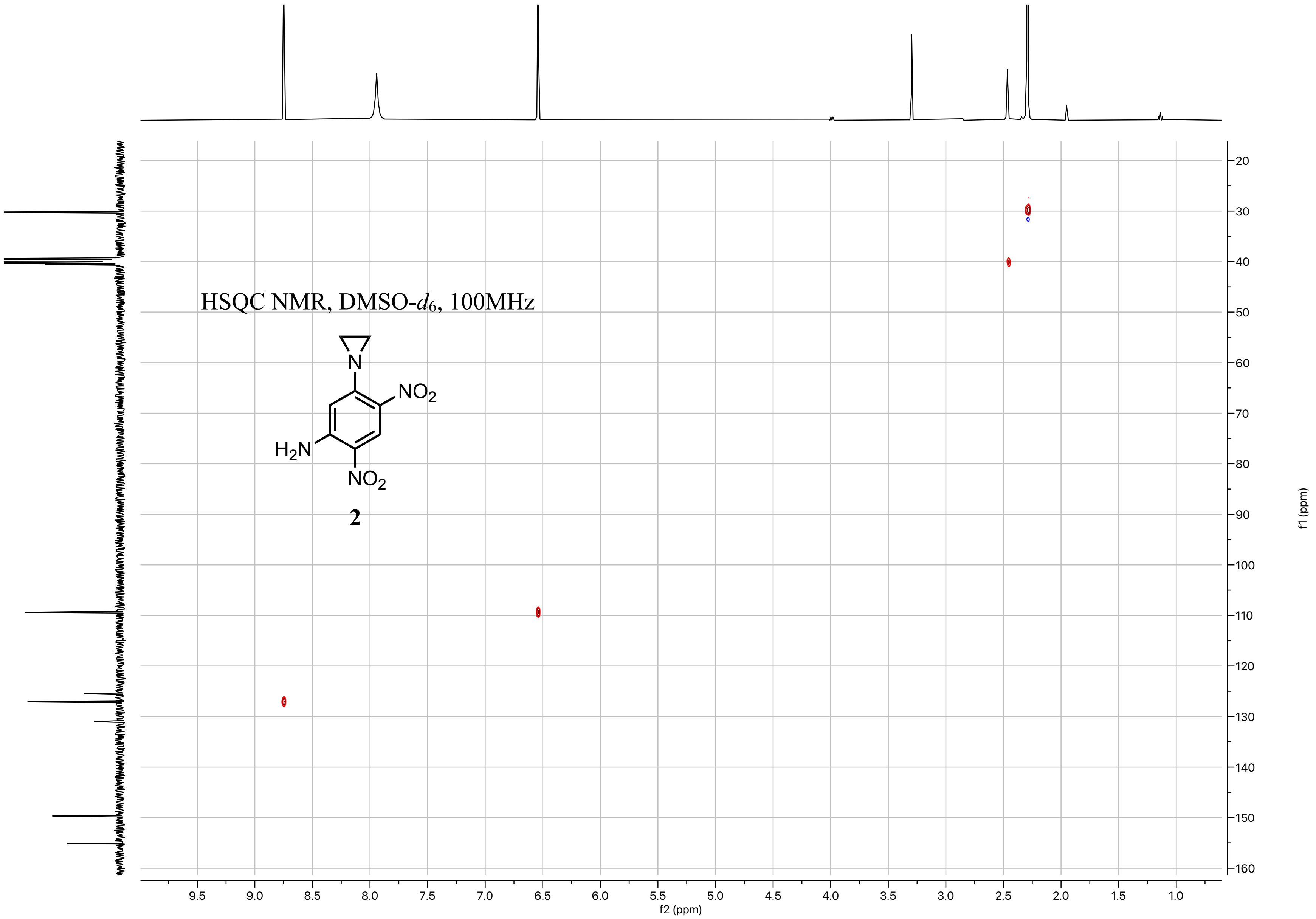
HSQC NMR, DMSO-$d_6$, 100MHz
NO2
H2N
NO2
N
2
f2 (ppm)
f1 (ppm)

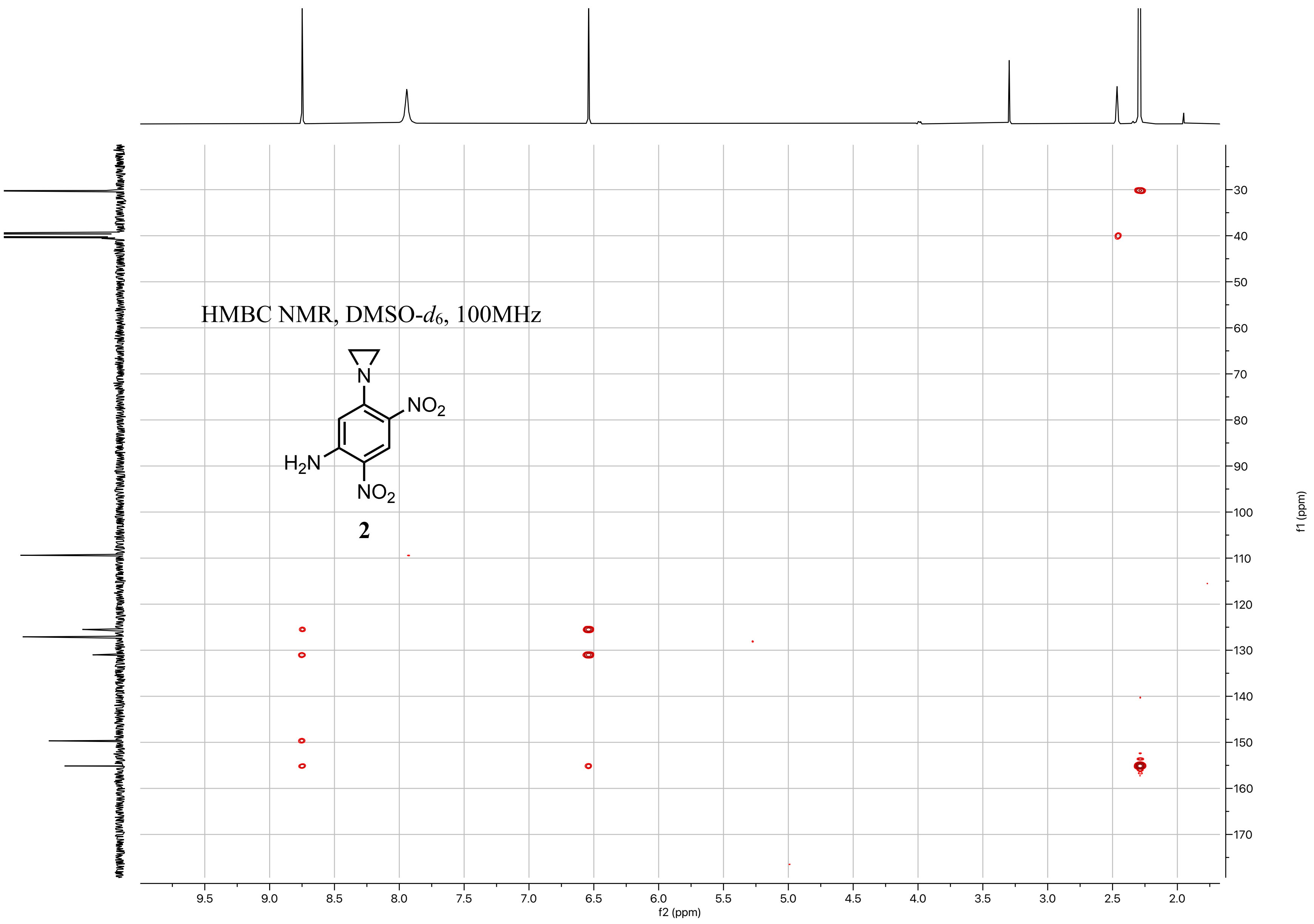
HMBC NMR, DMSO-$d_6$, 100MHz
N
$NO_2$
$H_2N$
$NO_2$
2
f2 (ppm)
f1 (ppm)
9.5
9.0
8.5
8.0
7.5
7.0
6.5
6.0
5.5
5.0
4.5
4.0
3.5
3.0
2.5
2.0
30
40
50
60
70
80
90
100
110
120
130
140
150
160
170

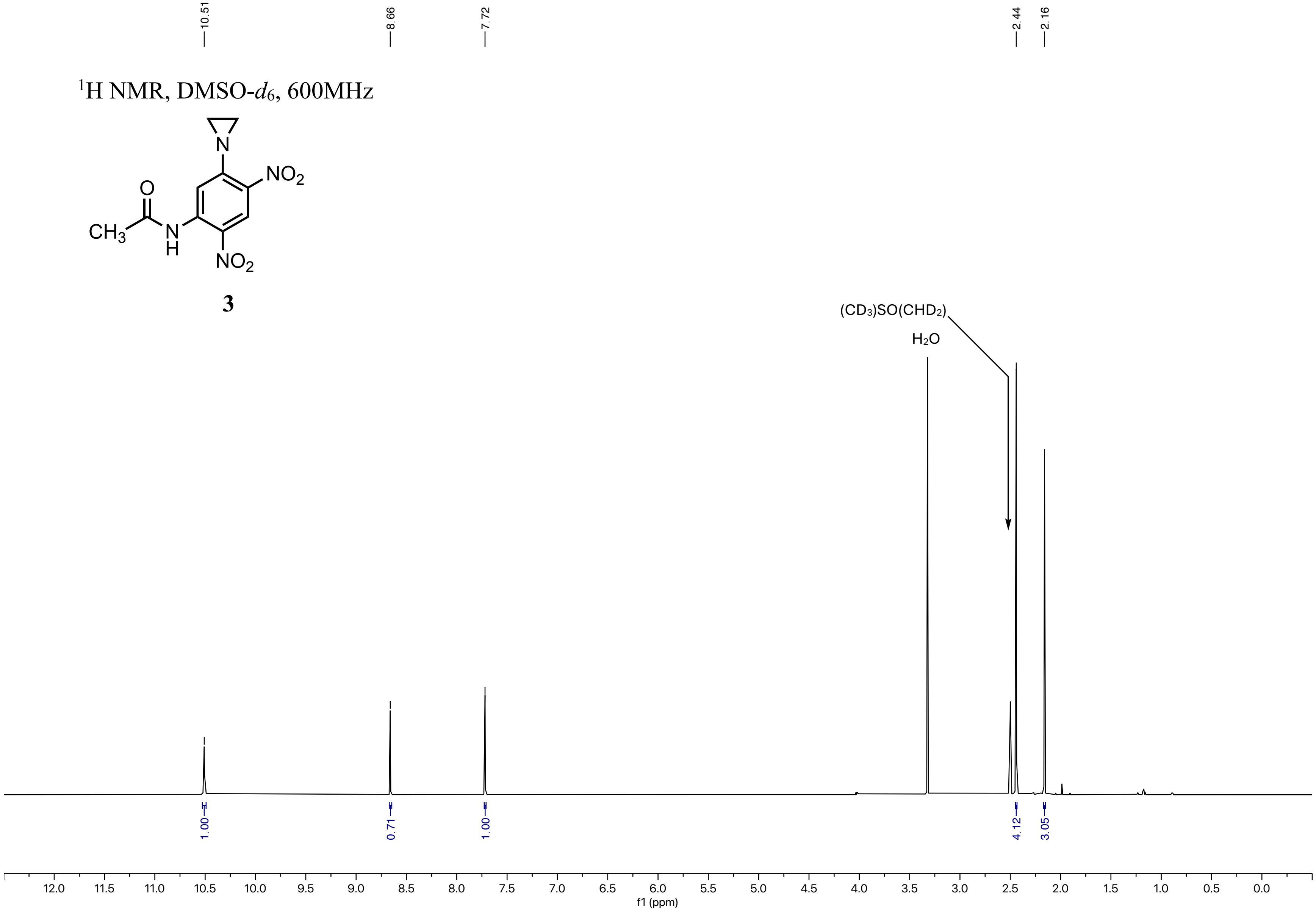
10.51
8.66
7.72
2.44
2.16
1H NMR, DMSO-d6, 600MHz
NO2
CH3
O
N
H
NO2
3
(CD3)SO(CHD2)
H2O
1.00
0.71
1.00
4.12
3.05
12.0 11.5 11.0 10.5 10.0 9.5 9.0 8.5 8.0 7.5 7.0 6.5 6.0 5.5 5.0 4.5 4.0 3.5 3.0 2.5 2.0 1.5 1.0 0.5 0.0
f1 (ppm)

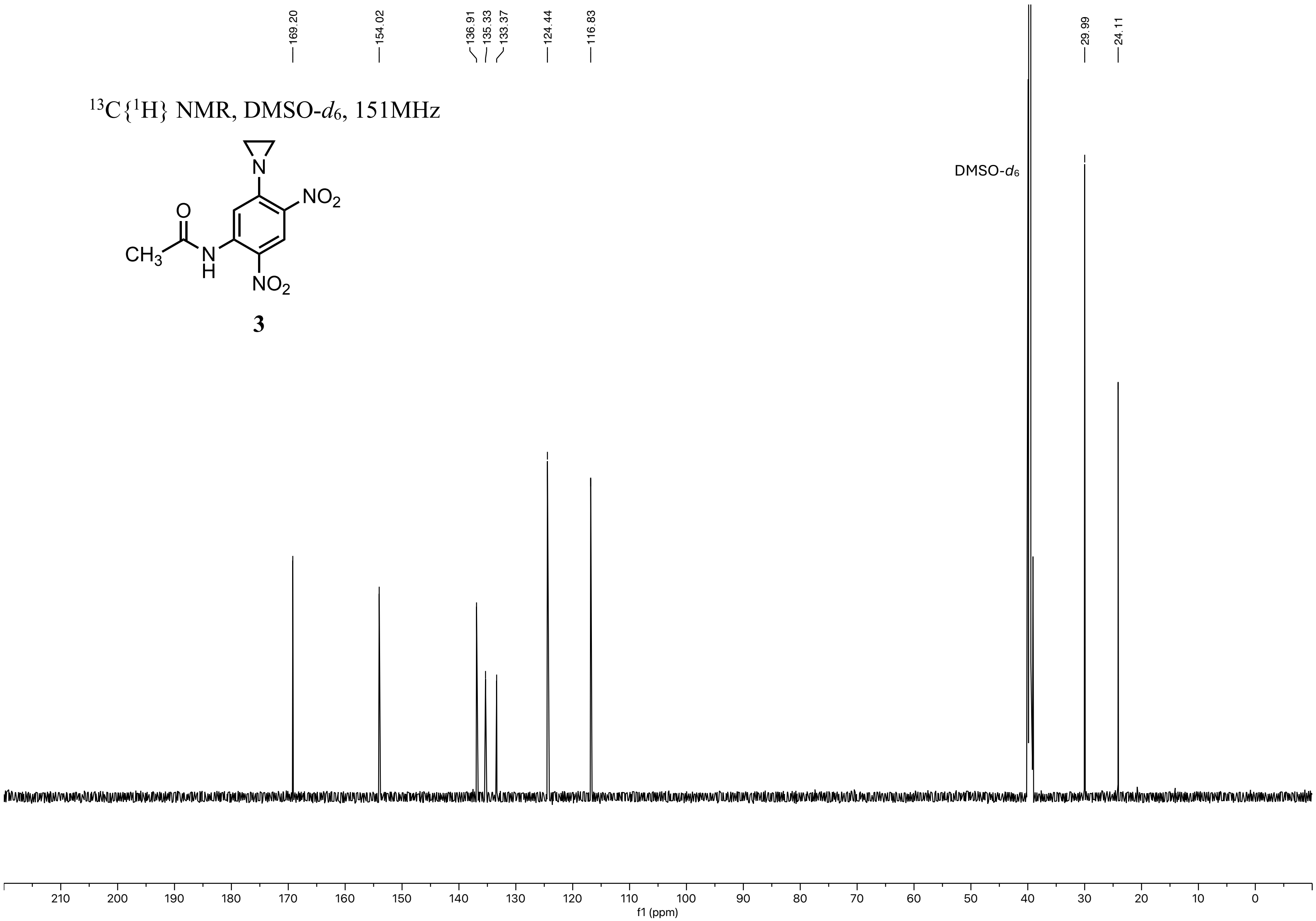
169.20
154.02
136.91
135.33
133.37
124.44
116.83
29.99
24.11
13C{1H} NMR, DMSO-d6, 151MHz
NO2
O
CH3
N
H
NO2
3
DMSO-d6
210
200
190
180
170
160
150
140
130
120
110
100
90
80
70
60
50
40
30
20
10
0
f1 (ppm)

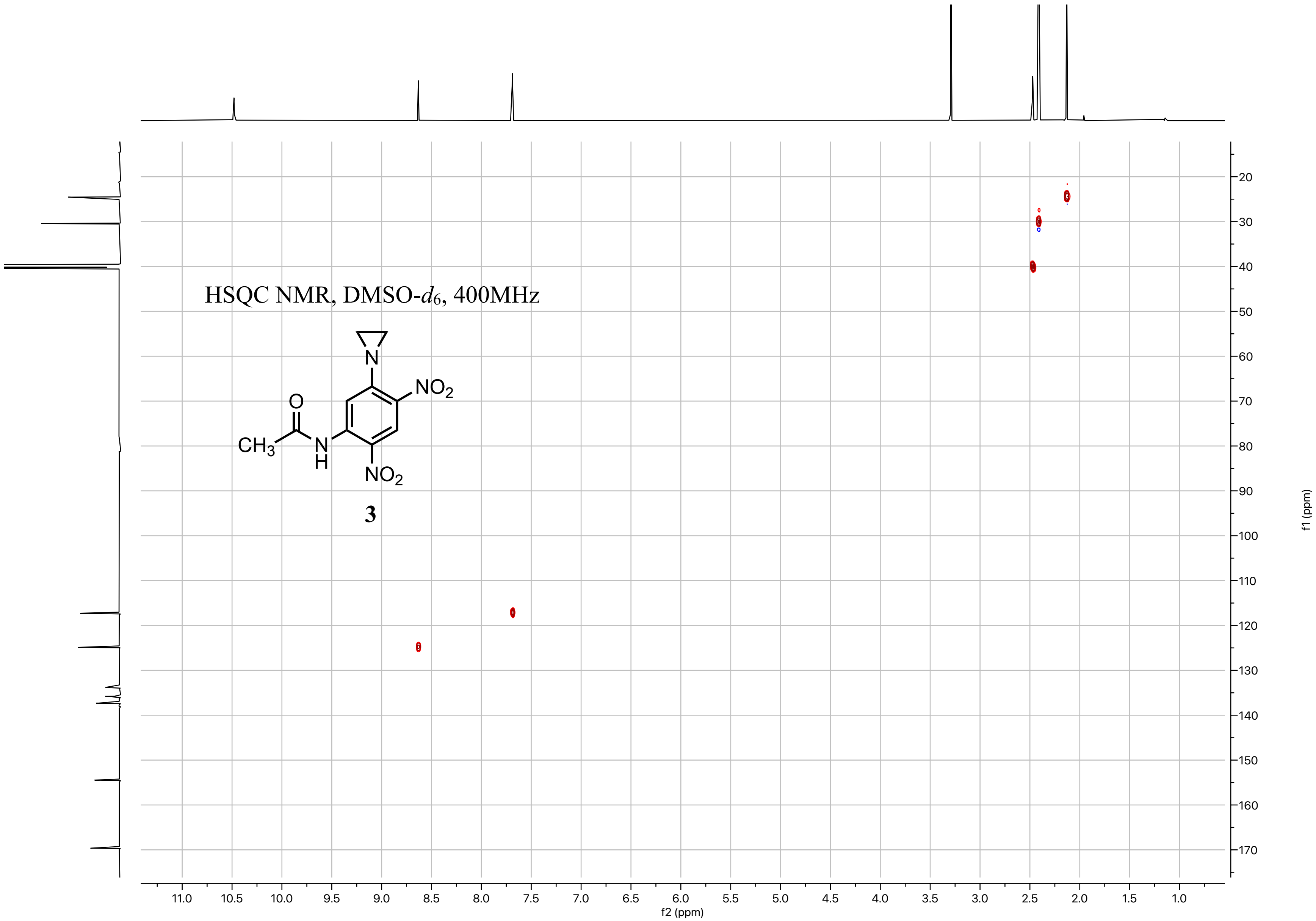

HSQC NMR, DMSO-d6, 400MHz
O
CH3
N
H
N
NO2
NO2
3
f2 (ppm)
f1 (ppm)

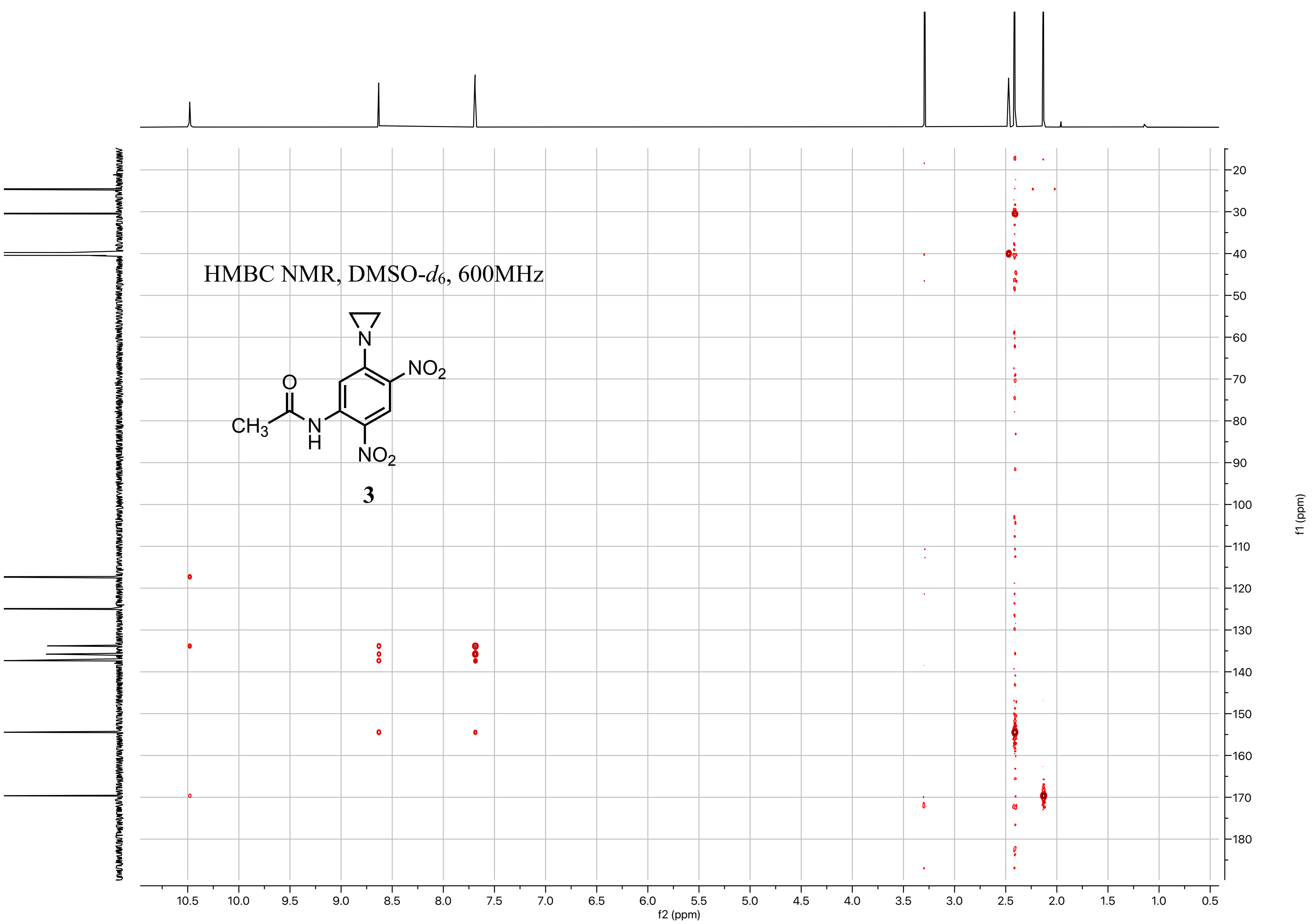
HMBC NMR, DMSO-$d_6$, 600MHz
N
$NO_2$
O
$CH_3$
N
H
$NO_2$
3
10.5
10.0
9.5
9.0
8.5
8.0
7.5
7.0
6.5
6.0
5.5
5.0
4.5
4.0
3.5
3.0
2.5
2.0
1.5
1.0
0.5
f2 (ppm)
20
30
40
50
60
70
80
90
100
110
120
130
140
150
160
170
180
f1 (ppm)

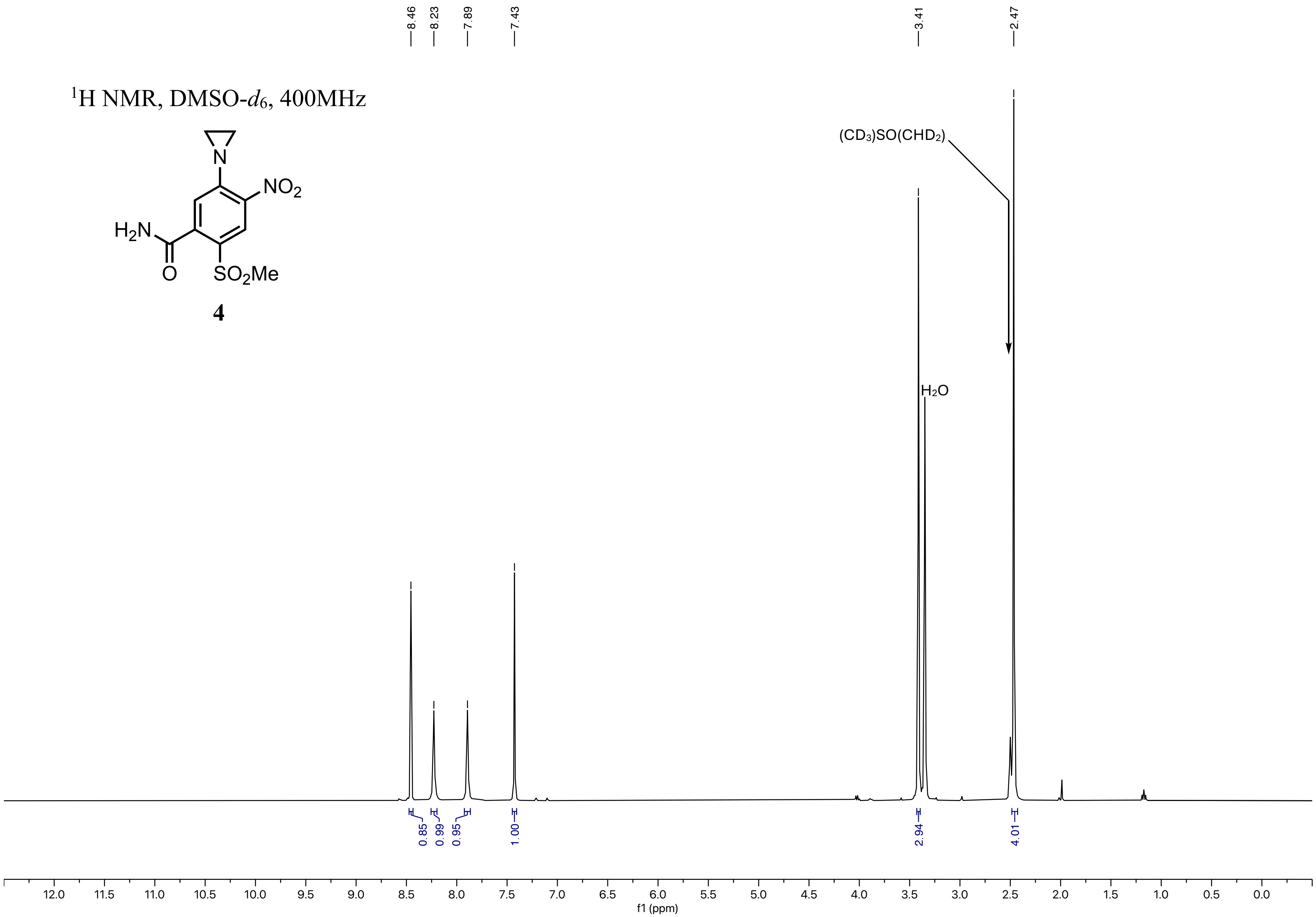

$^1H$ NMR, DMSO-$d_6$, 400MHz
NO2
H2N
O
SO2Me
N
4
8.46
8.23
7.89
7.43
3.41
2.47
(CD3)SO(CHD2)
H2O
0.85
0.99
0.95
1.00
2.94
4.01
f1 (ppm)

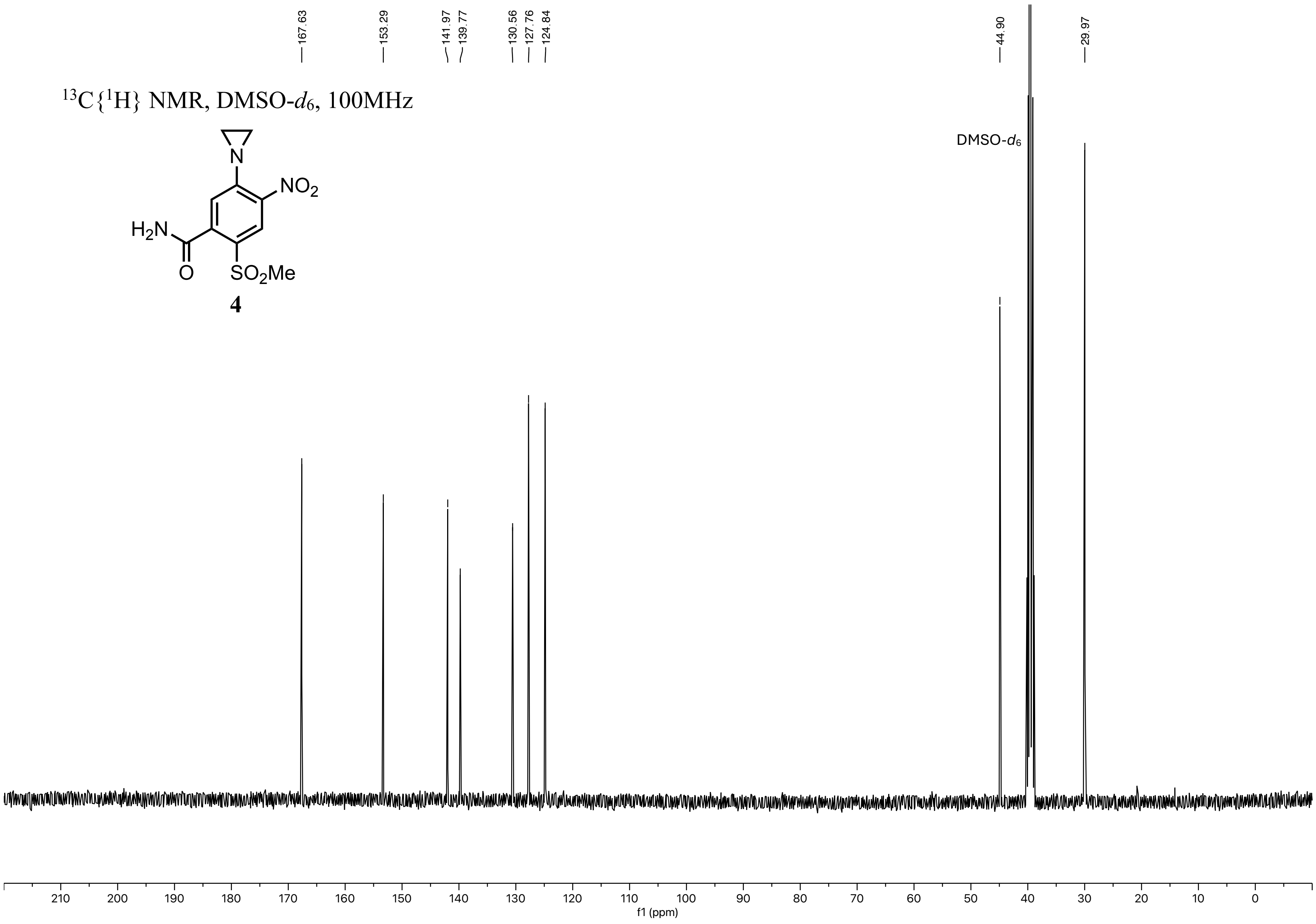

$^{13}C\{^{1}H\}$ NMR, DMSO-$d_6$, 100MHz
H2N
O
N
NO2
SO2Me
4
167.63
153.29
141.97
139.77
130.56
127.76
124.84
44.90
29.97
DMSO-$d_6$
210
200
190
180
170
160
150
140
130
120
110
100
90
80
70
60
50
40
30
20
10
0
f1 (ppm)

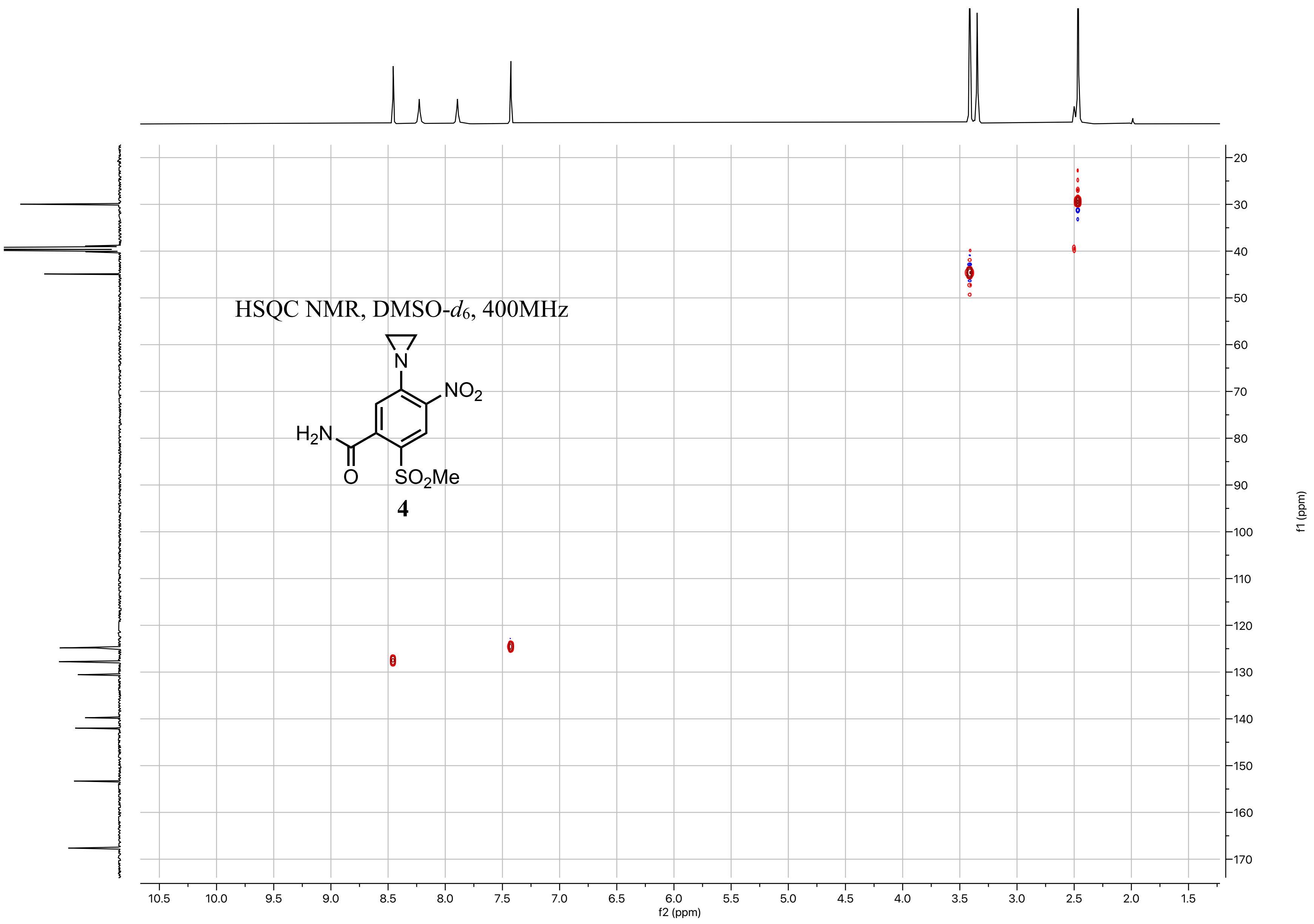

HSQC NMR, DMSO-d6, 400MHz
N
NO2
H2N
O
SO2Me
4
f2 (ppm)
f1 (ppm)

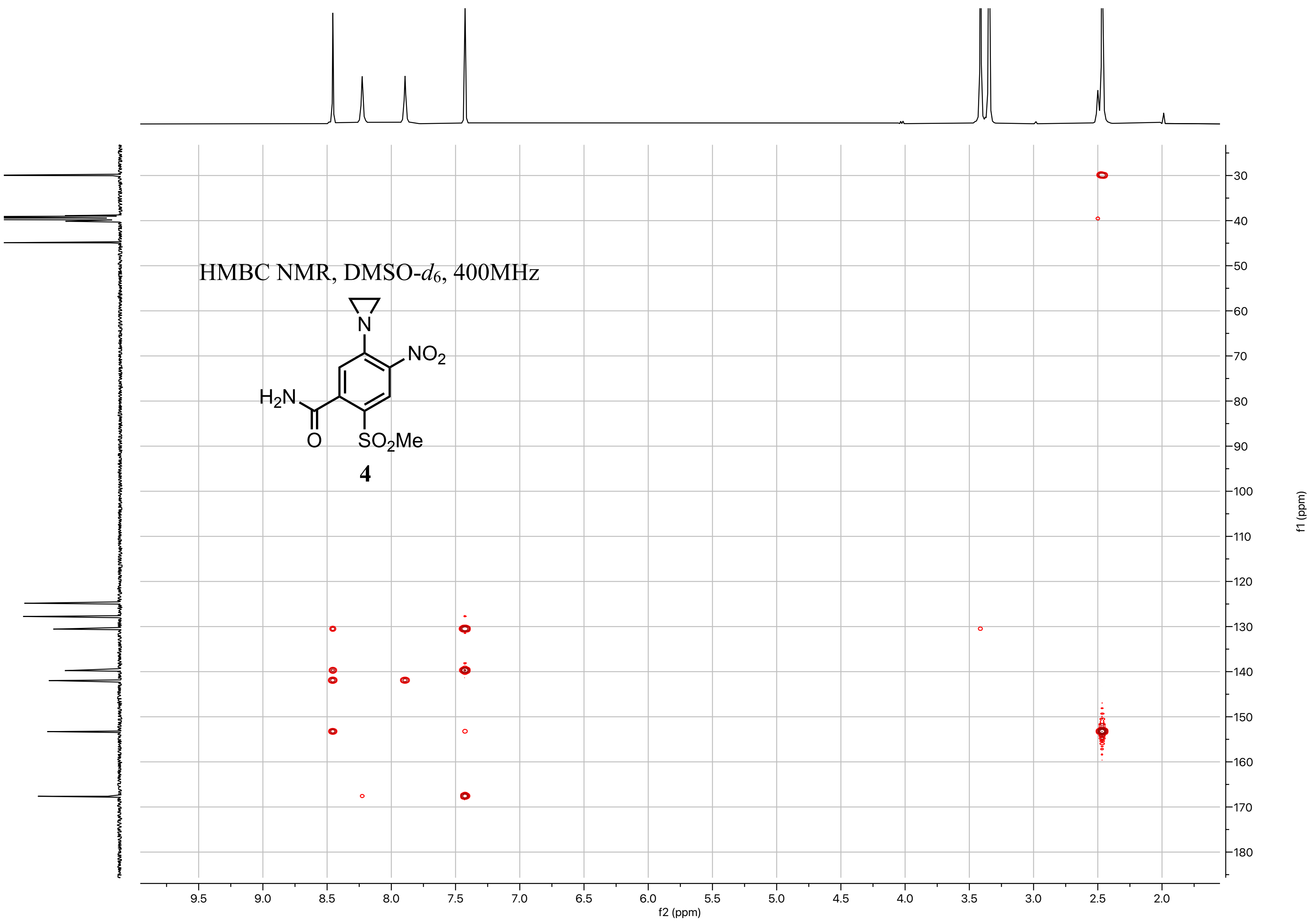
HMBC NMR, DMSO-d6, 400MHz
NO2
H2N
O
SO2Me
N
4
f2 (ppm)
f1 (ppm)

$^{1}H$ NMR, DMSO-$d_6$, 400MHz

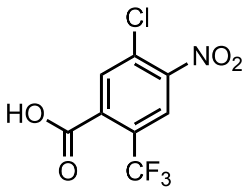


**S5**

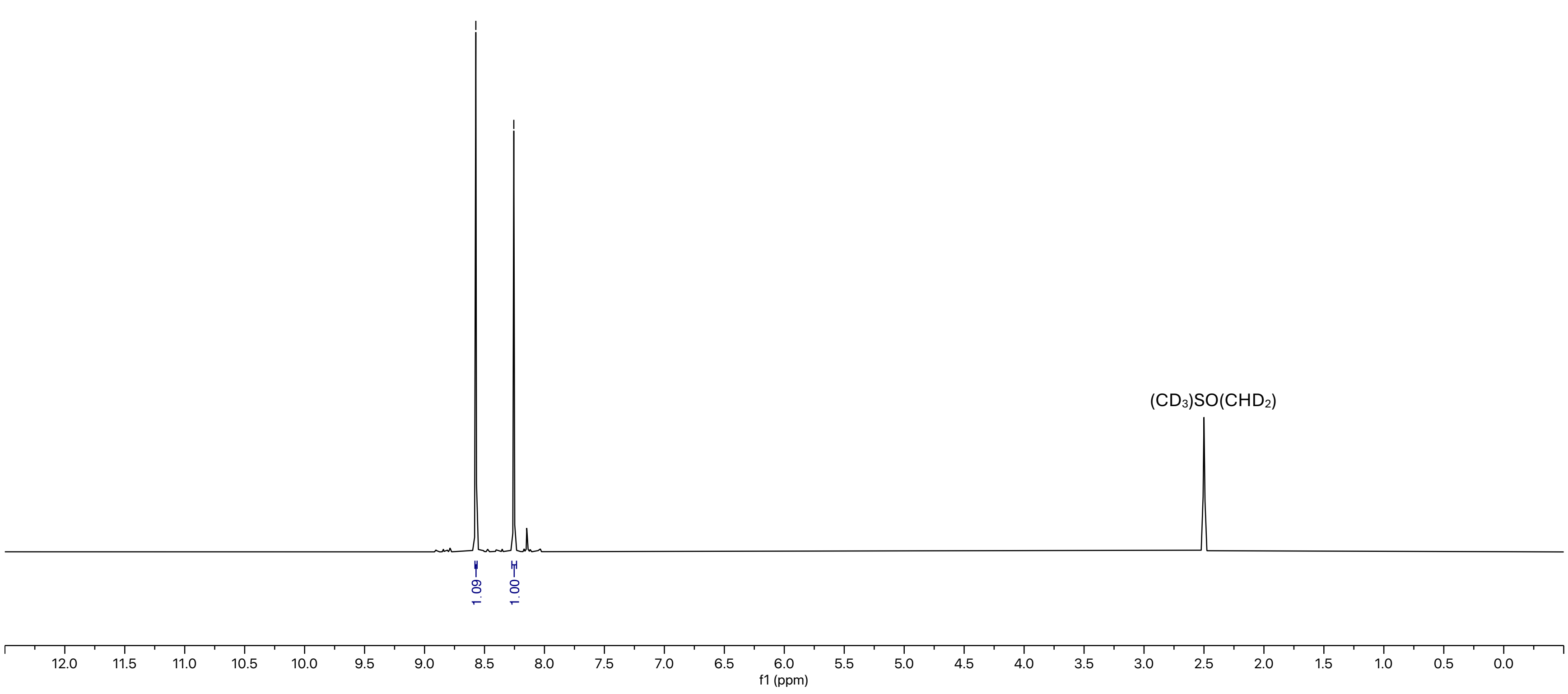

$^{13}C\{^1H\}$ NMR, DMSO-$d_6$, 100MHz

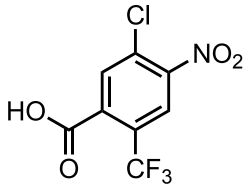


**S5**

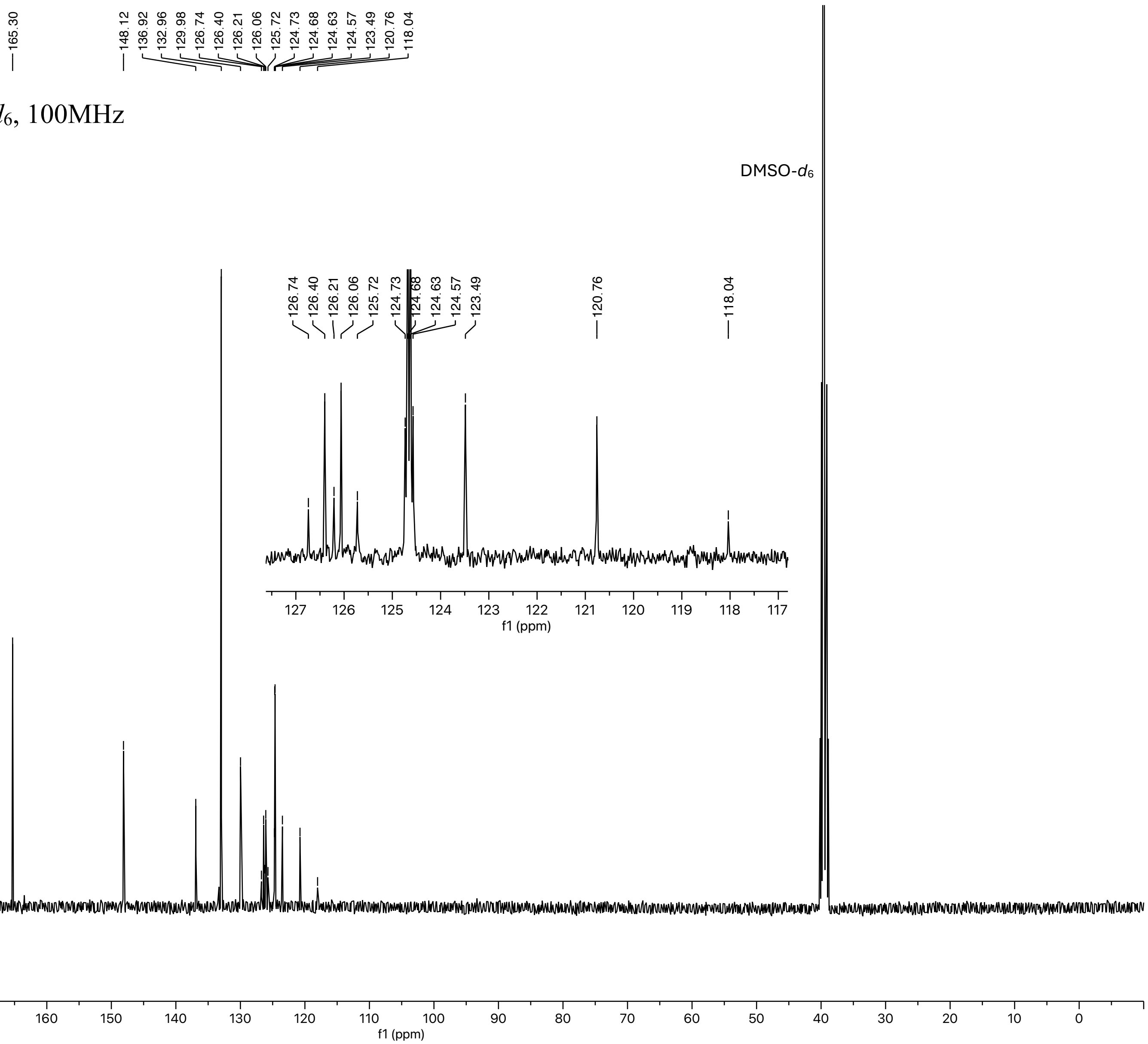

$^{19}F$ NMR, DMSO-$d_6$, 376MHz

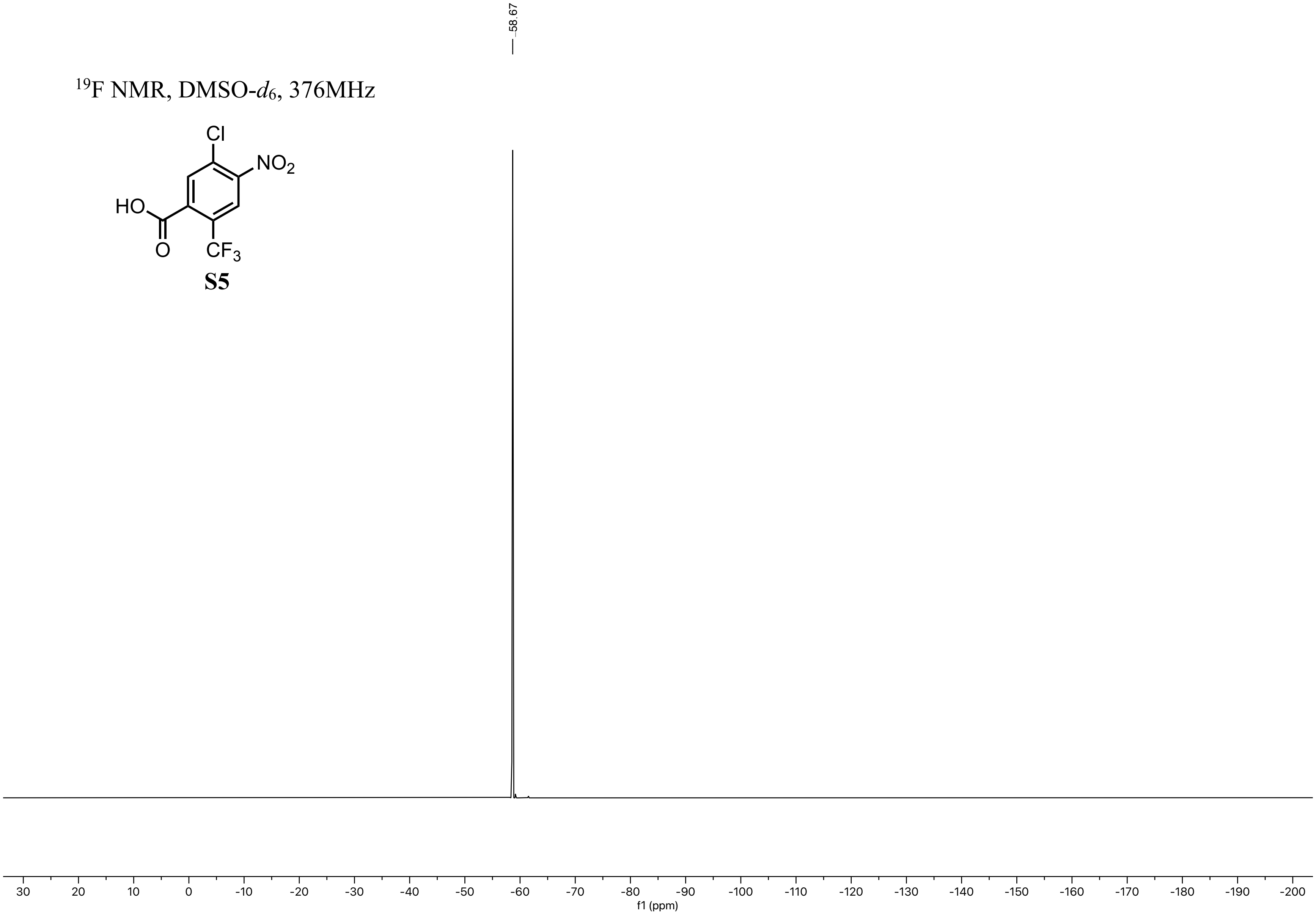

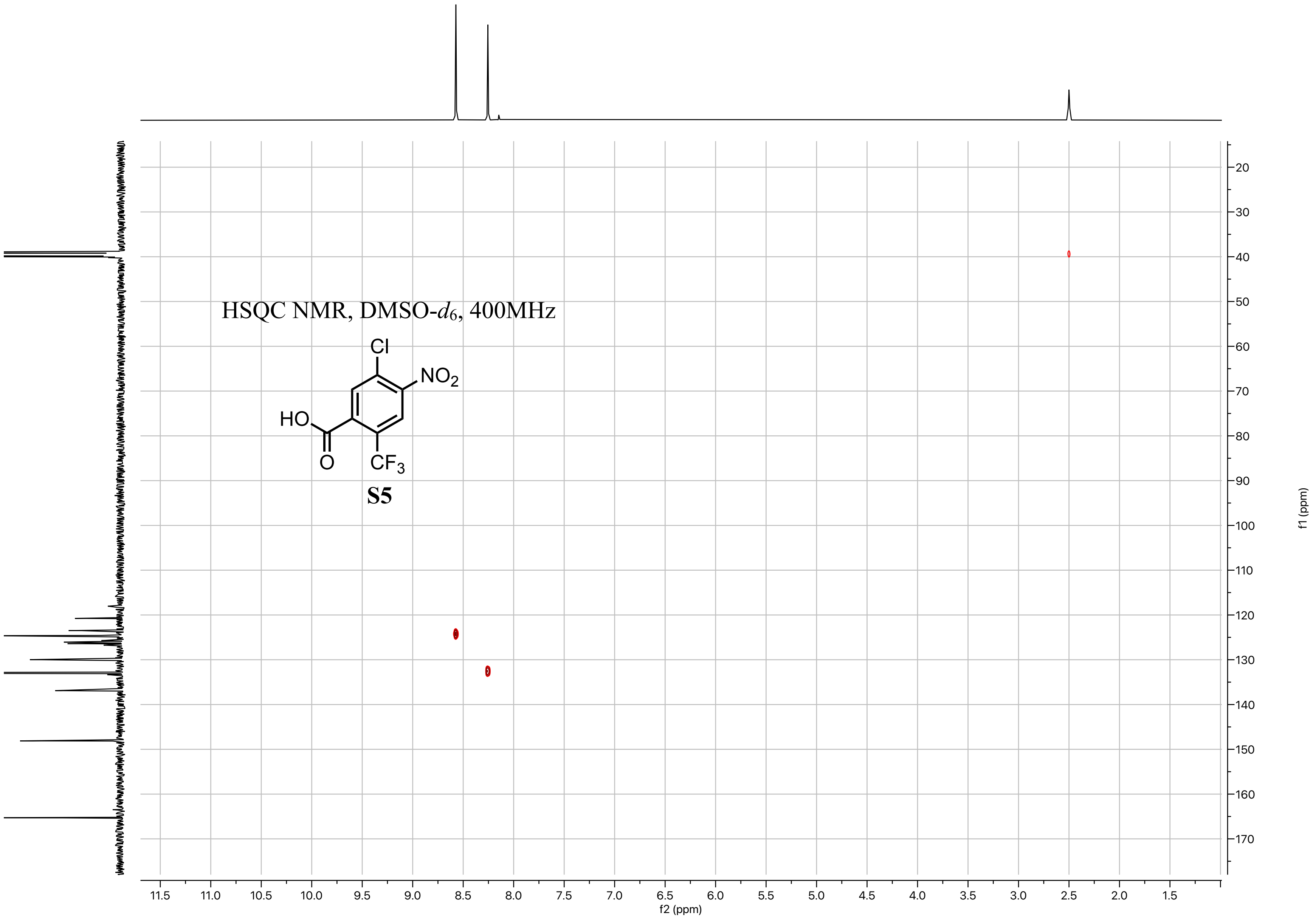
HSQC NMR, DMSO-d6, 400MHz
Cl
NO2
HO
O
CF3
S5
f2 (ppm)
f1 (ppm)

$^{1}H$ NMR, DMSO-$d_6$, 500MHz

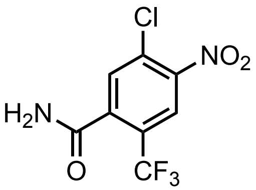

**S6**

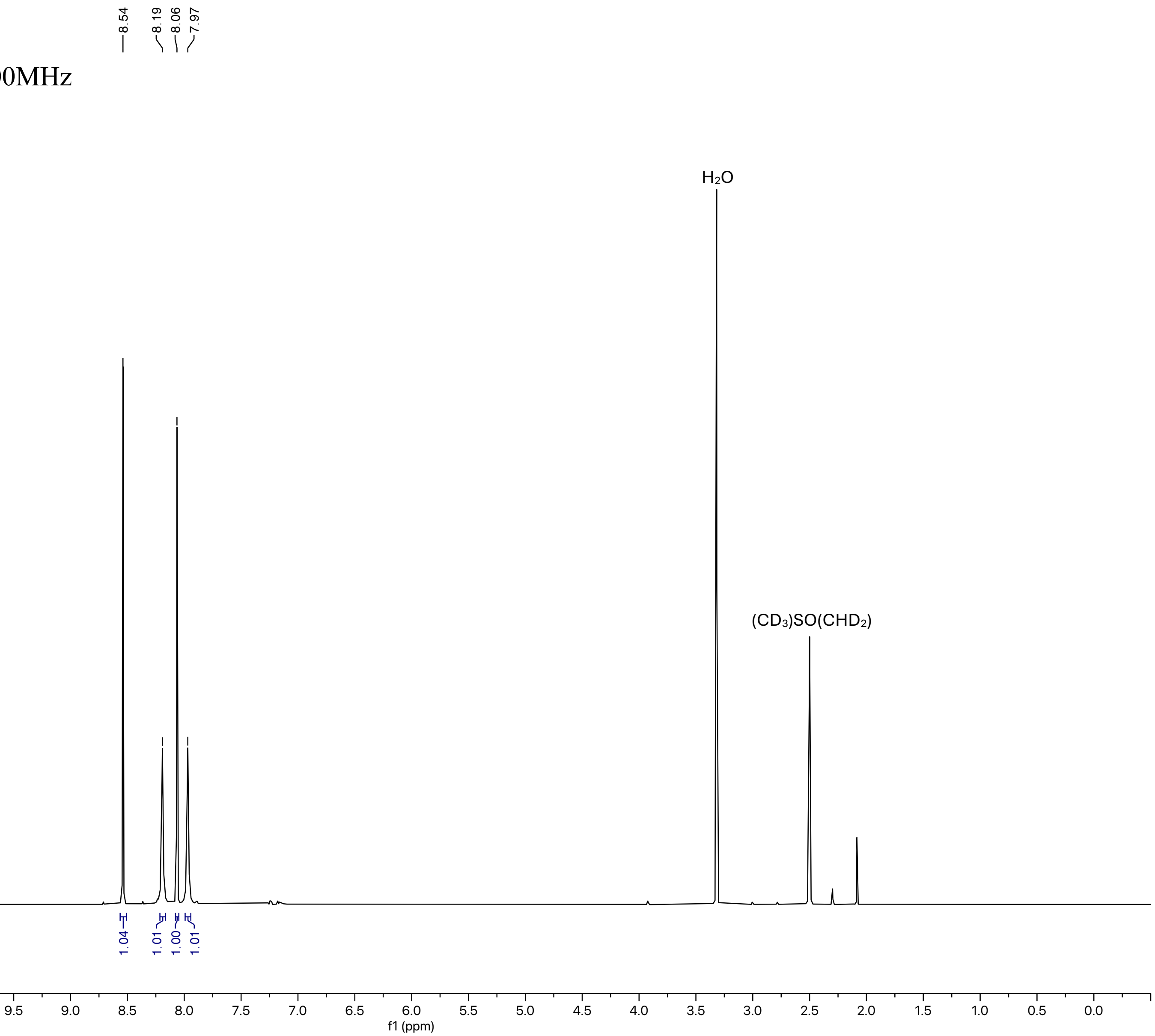

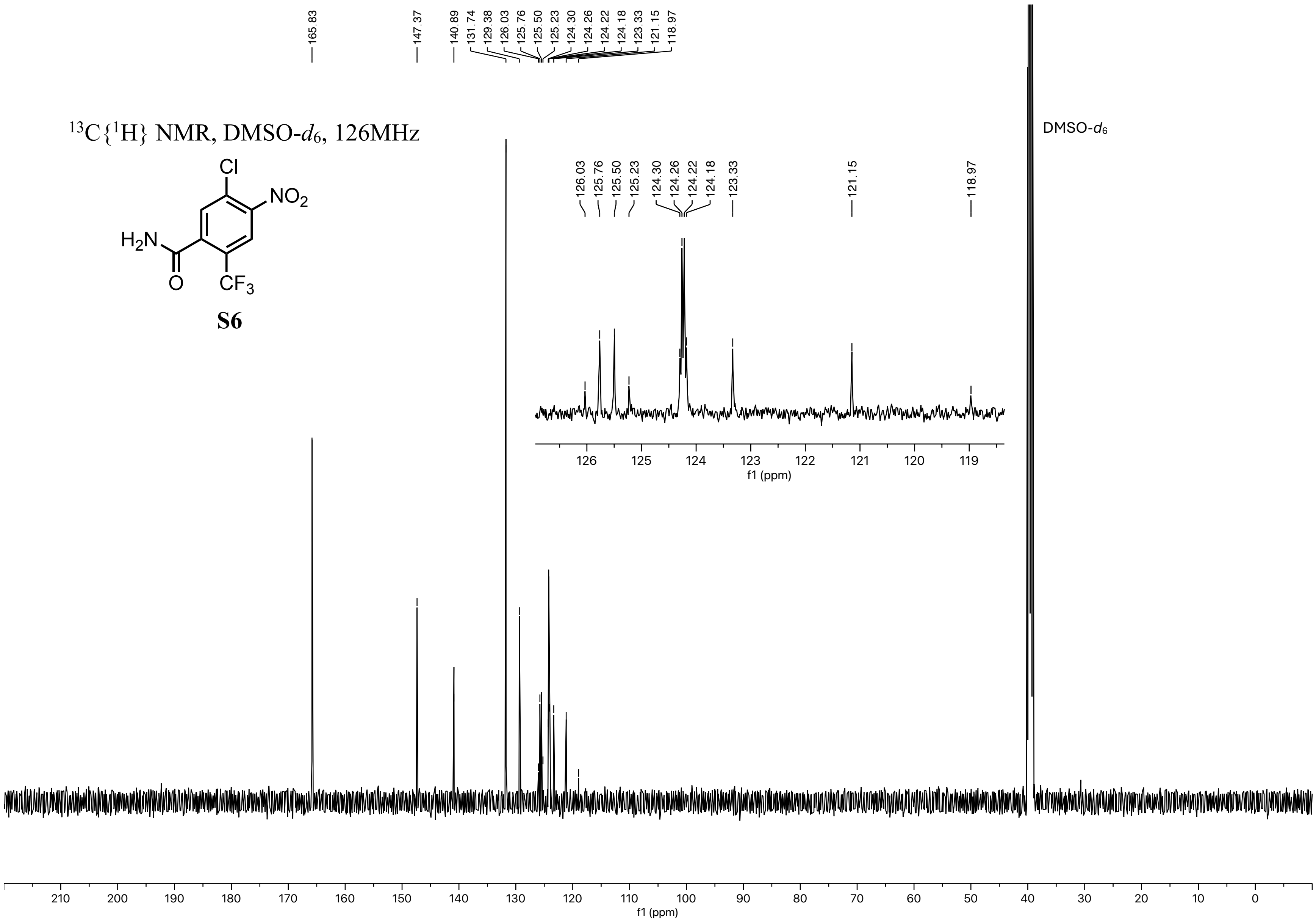

165.83
147.37
140.89
131.74
129.38
126.03
125.76
125.50
125.23
124.30
124.26
124.22
124.18
123.33
121.15
118.97
$^{13}C\{^{1}H\}$ NMR, DMSO-$d_6$, 126MHz
Cl
$NO_2$
$H_2N$
O
$CF_3$
S6
DMSO-$d_6$
f1 (ppm)

$^{19}F$ NMR, DMSO-$d_6$, 470MHz

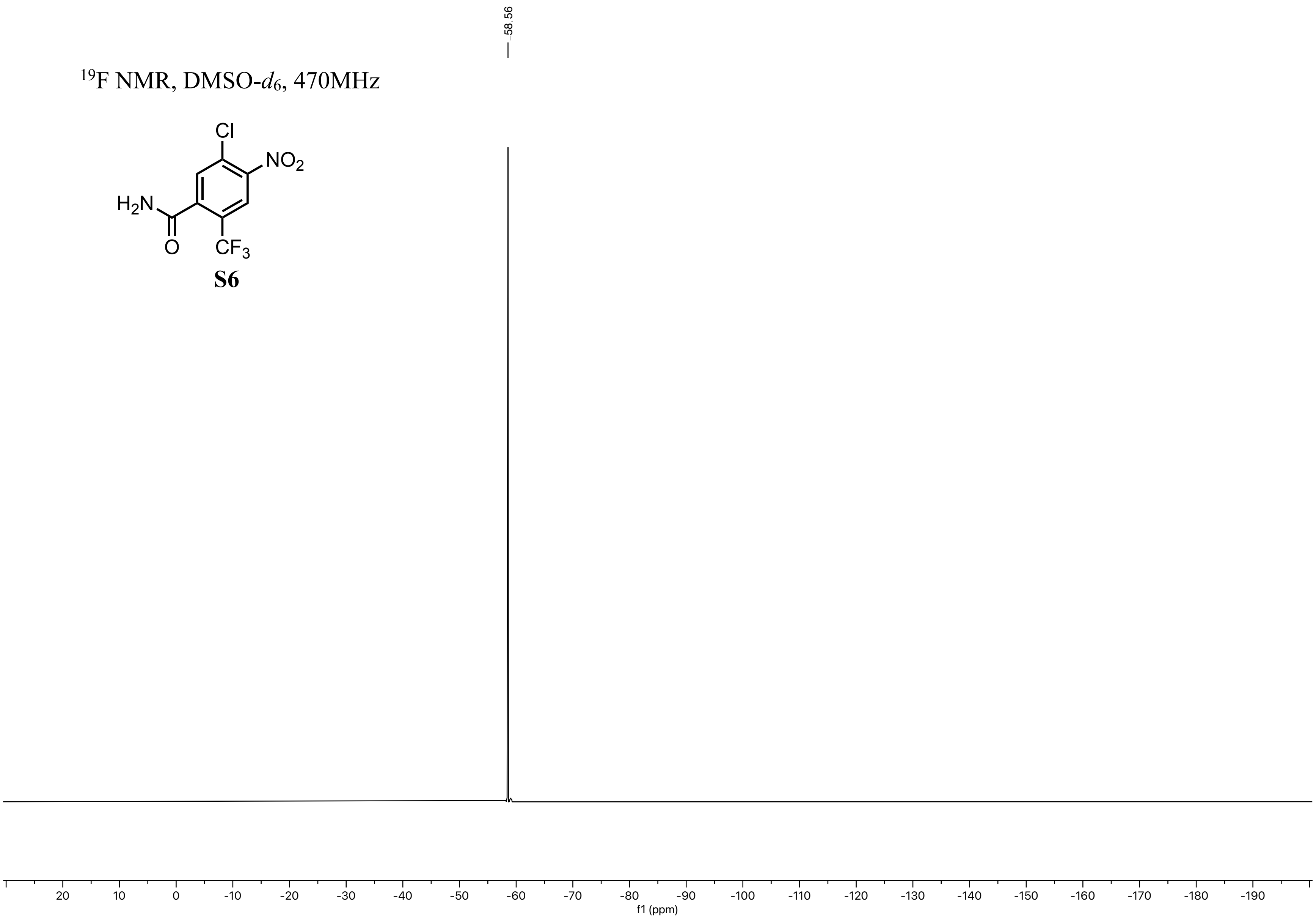


**S6**

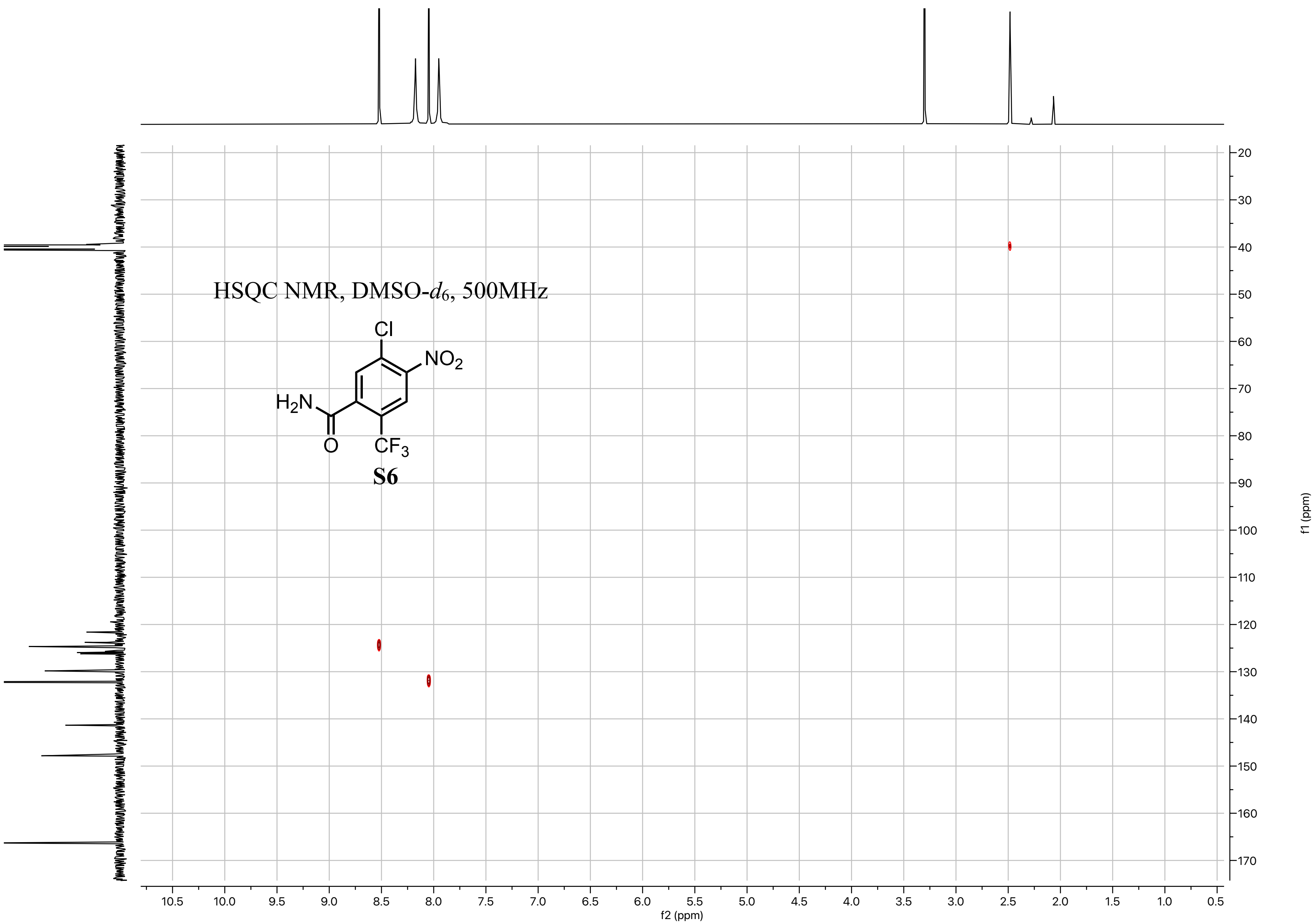

HSQC NMR, DMSO-d6, 500MHz
Cl
NO2
H2N
O
CF3
S6
f1 (ppm)
f2 (ppm)

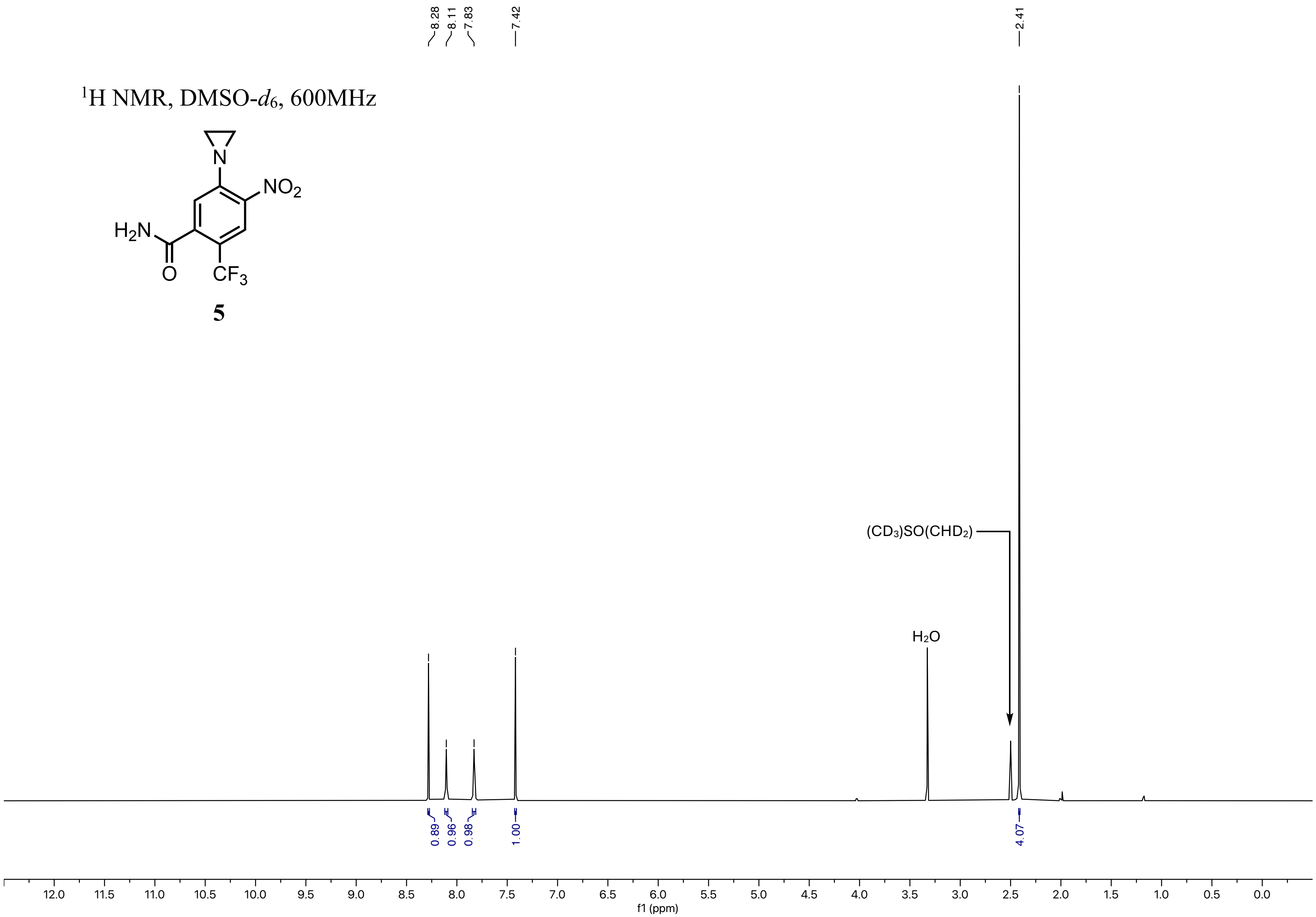

$^1$H NMR, DMSO-$d_6$, 600MHz
NO2
H2N
O
CF3
N
5
8.28
8.11
7.83
7.42
2.41
(CD3)SO(CHD2)
H2O
0.89
0.96
0.98
1.00
4.07
12.0
11.5
11.0
10.5
10.0
9.5
9.0
8.5
8.0
7.5
7.0
6.5
6.0
5.5
5.0
4.5
4.0
3.5
3.0
2.5
2.0
1.5
1.0
0.5
0.0
f1 (ppm)

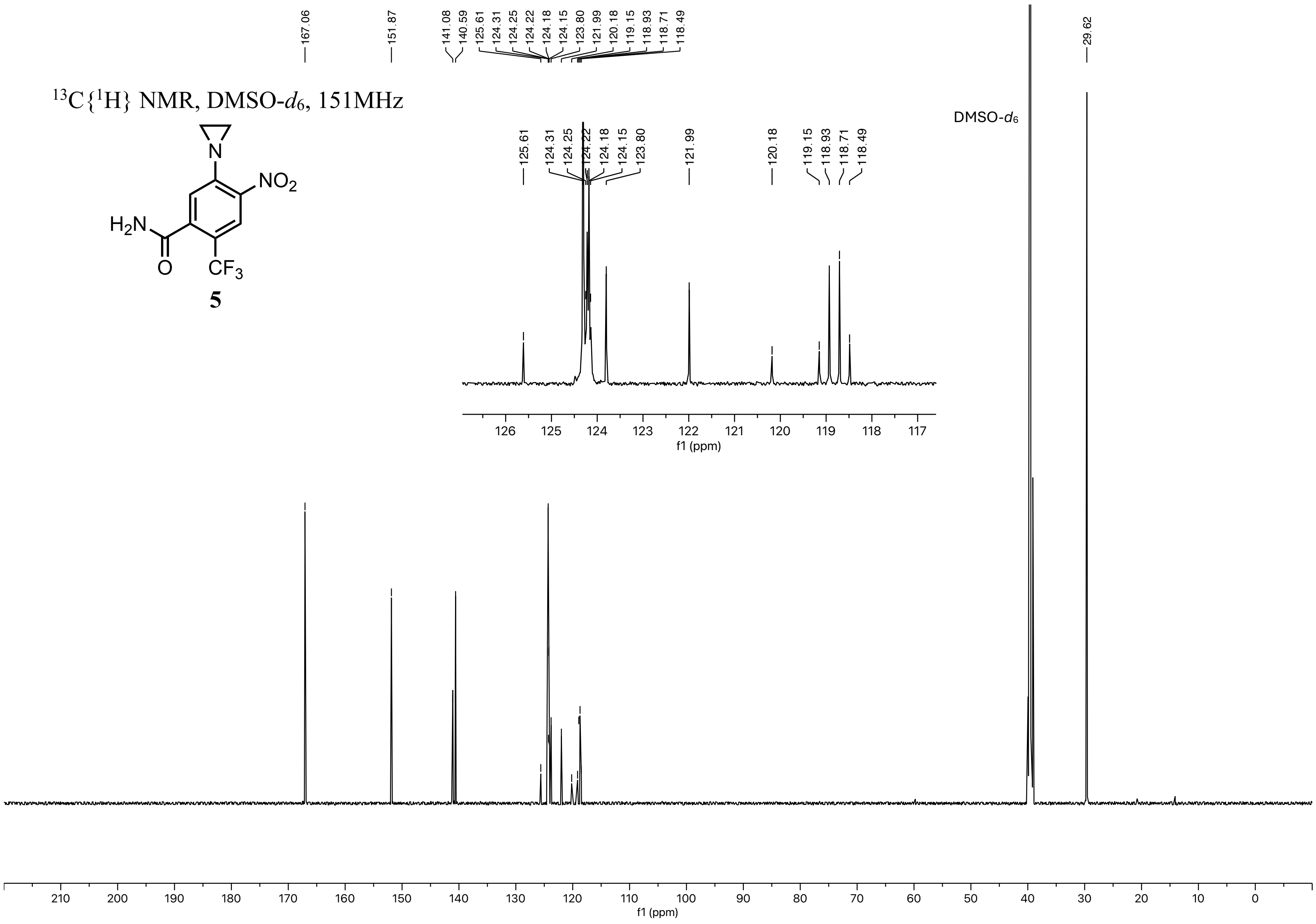

13C{1H} NMR, DMSO-d6, 151MHz
NO2
H2N
O
CF3
N
5
167.06
151.87
141.08
140.59
125.61
124.31
124.25
124.22
124.18
124.15
123.80
121.99
120.18
119.15
118.93
118.71
118.49
29.62
DMSO-d6
f1 (ppm)

$^{19}F$ NMR, DMSO-$d_6$, 376MHz

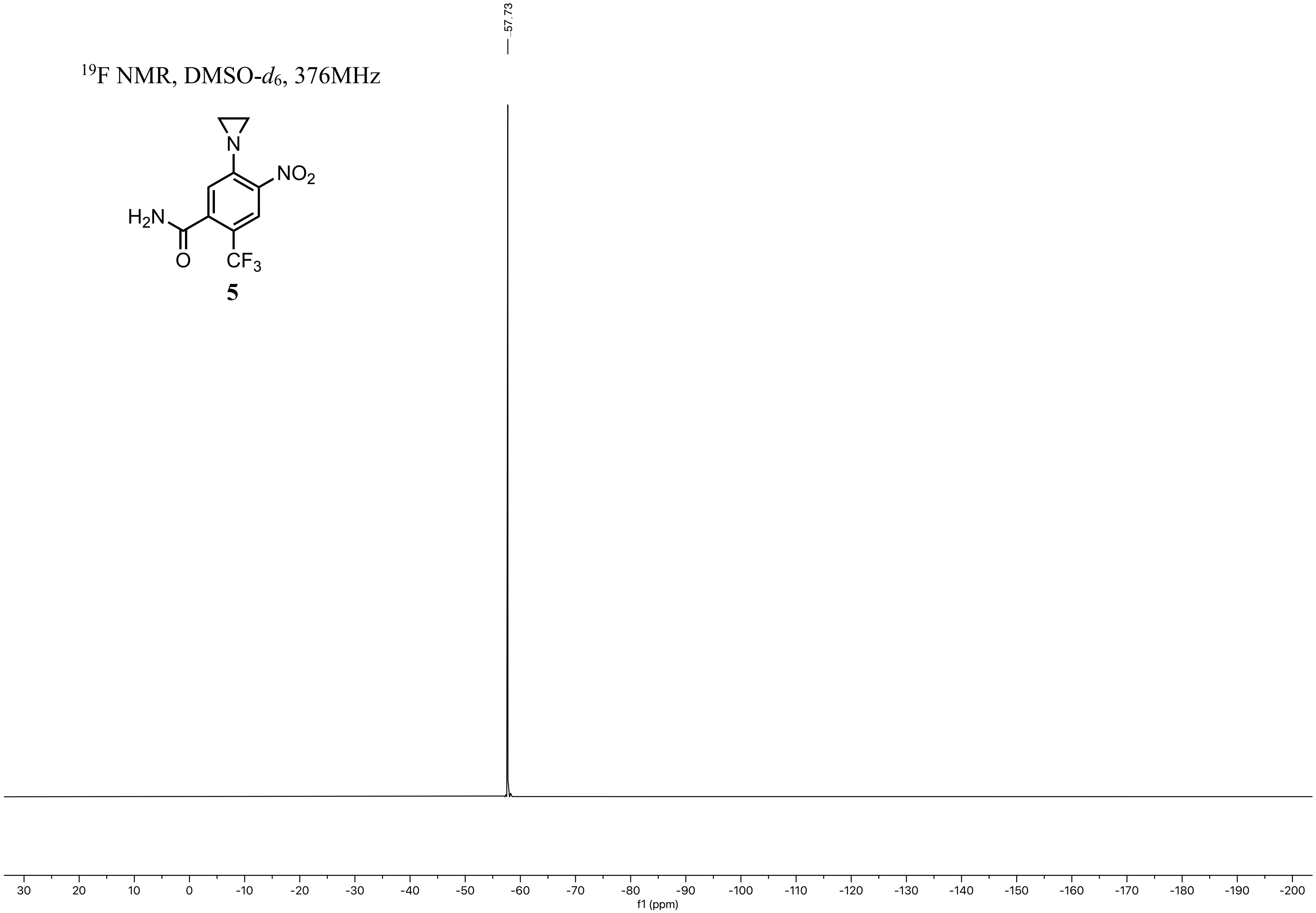


**5**

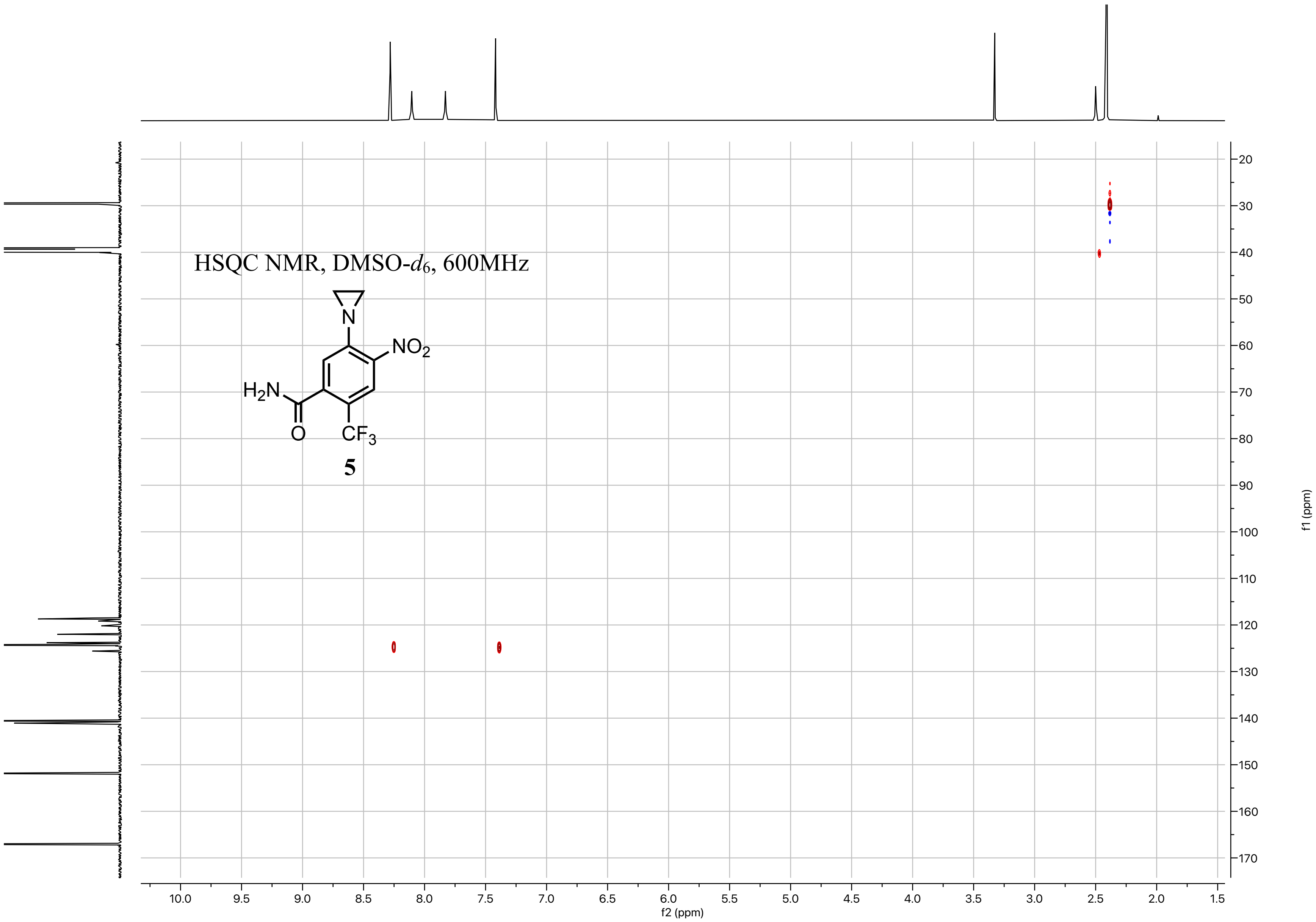

HSQC NMR, DMSO-d6, 600MHz
N
NO2
H2N
O
CF3
5
f2 (ppm)
f1 (ppm)

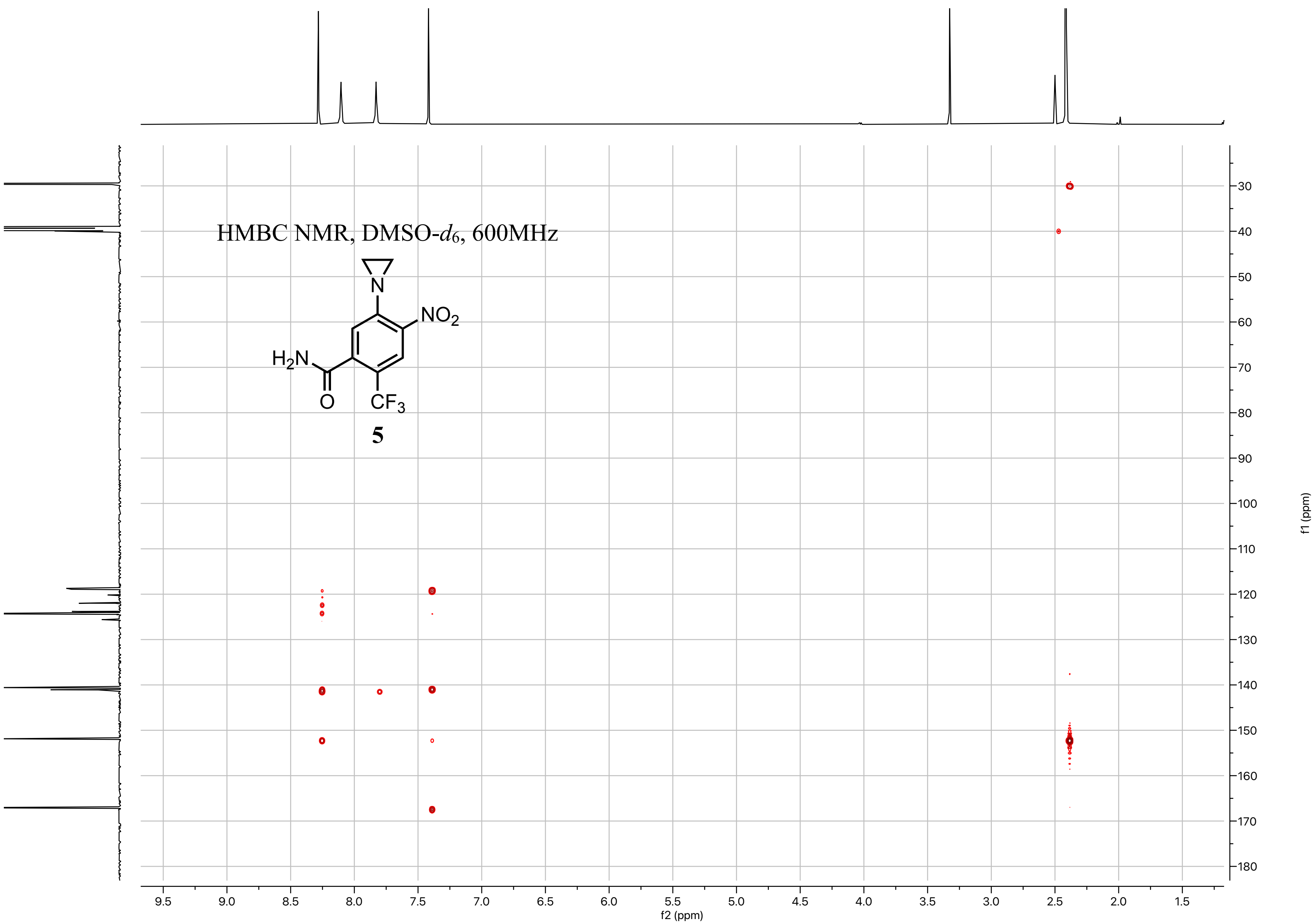
HMBC NMR, DMSO-d6, 600MHz
NO2
H2N
O
CF3
N
5
f2 (ppm)
f1 (ppm)

$^1H$ NMR, DMSO-$d_6$, 600MHz

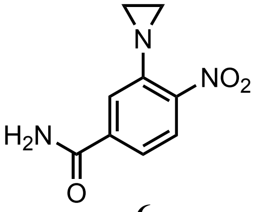


**6**

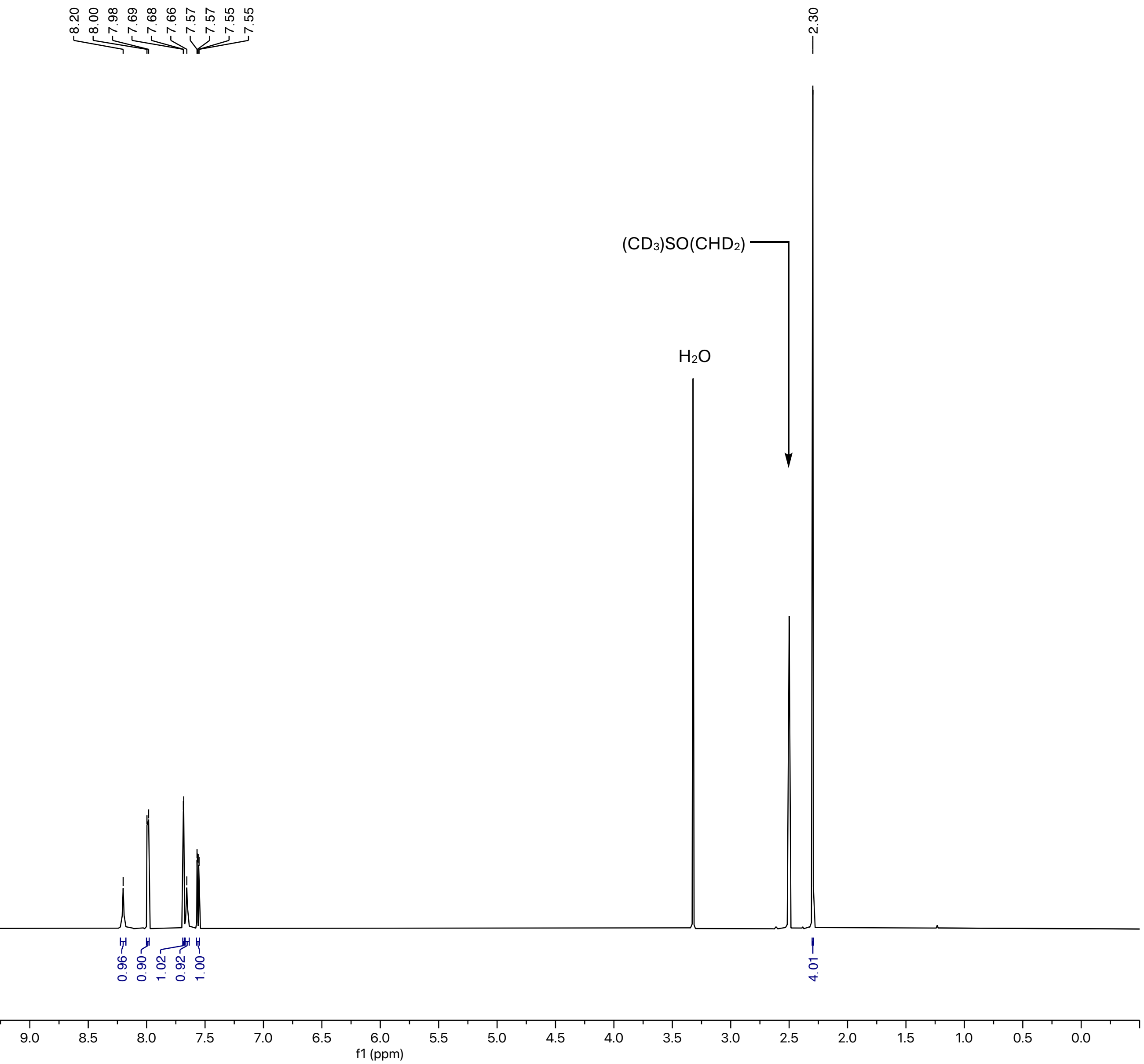

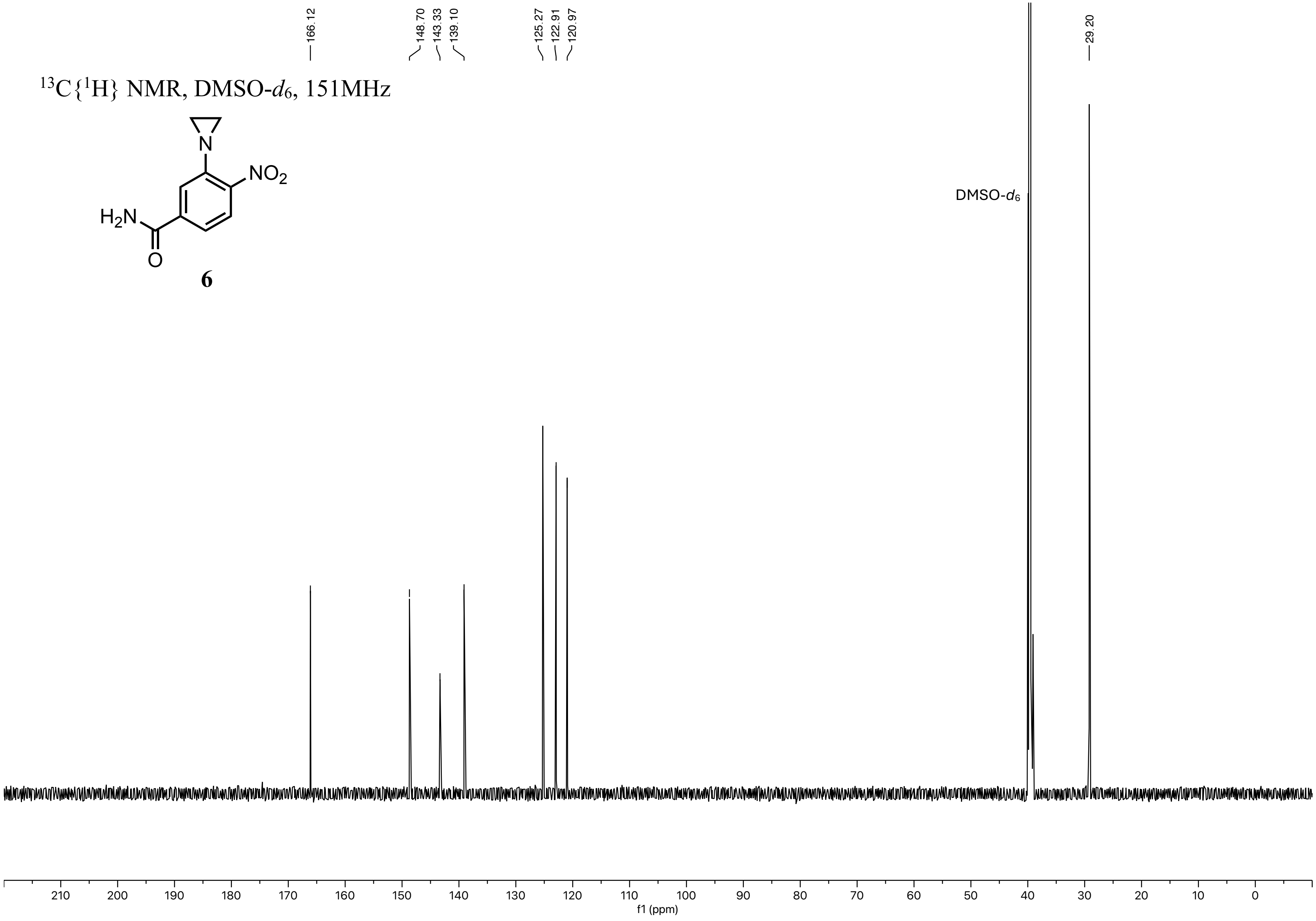

166.12
148.70
143.33
139.10
125.27
122.91
120.97
29.20
13C{1H} NMR, DMSO-d6, 151MHz
N
NO2
H2N
O
6
DMSO-d6
210
200
190
180
170
160
150
140
130
120
110
100
90
80
70
60
50
40
30
20
10
0
f1 (ppm)

HSQC NMR, DMSO-$d_6$, 400MHz

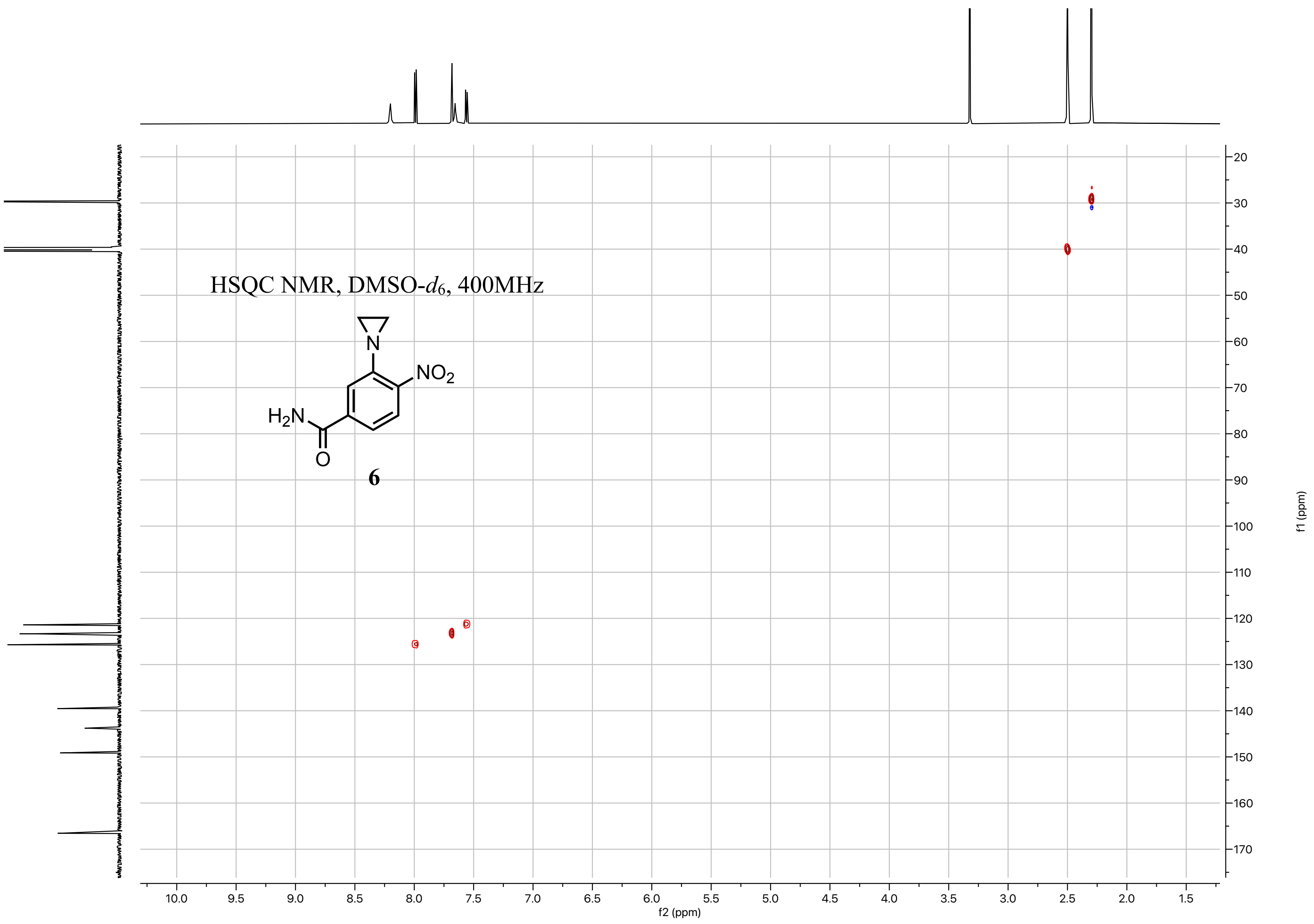


**6**

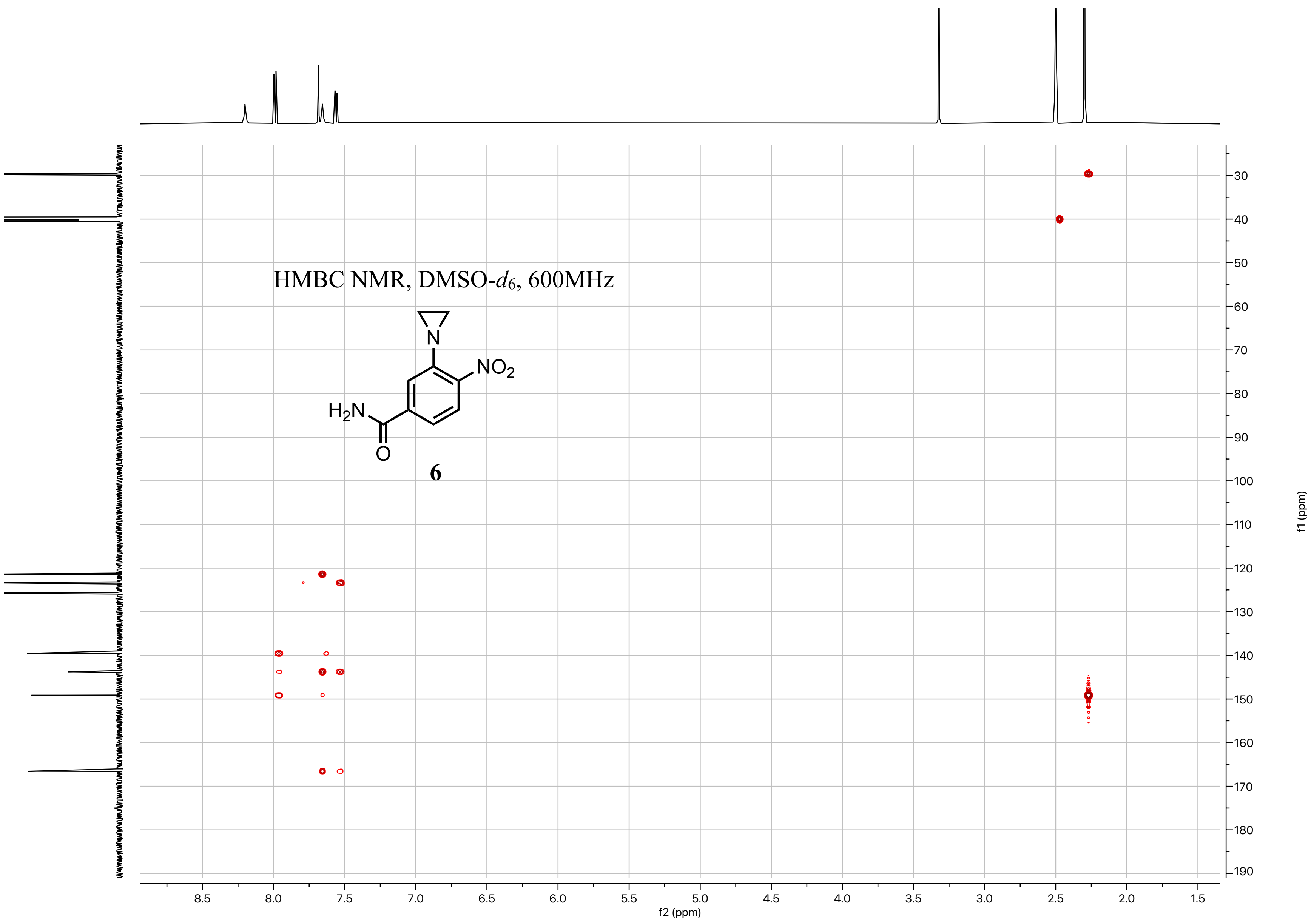
HMBC NMR, DMSO-$d_6$, 600MHz
N
$NO_2$
$H_2N$
O
6
f2 (ppm)
f1 (ppm)

[1]H NMR, DMSO-$d_6$, 600MHz

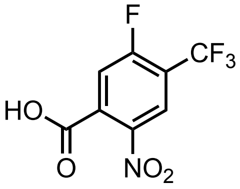

**S9**

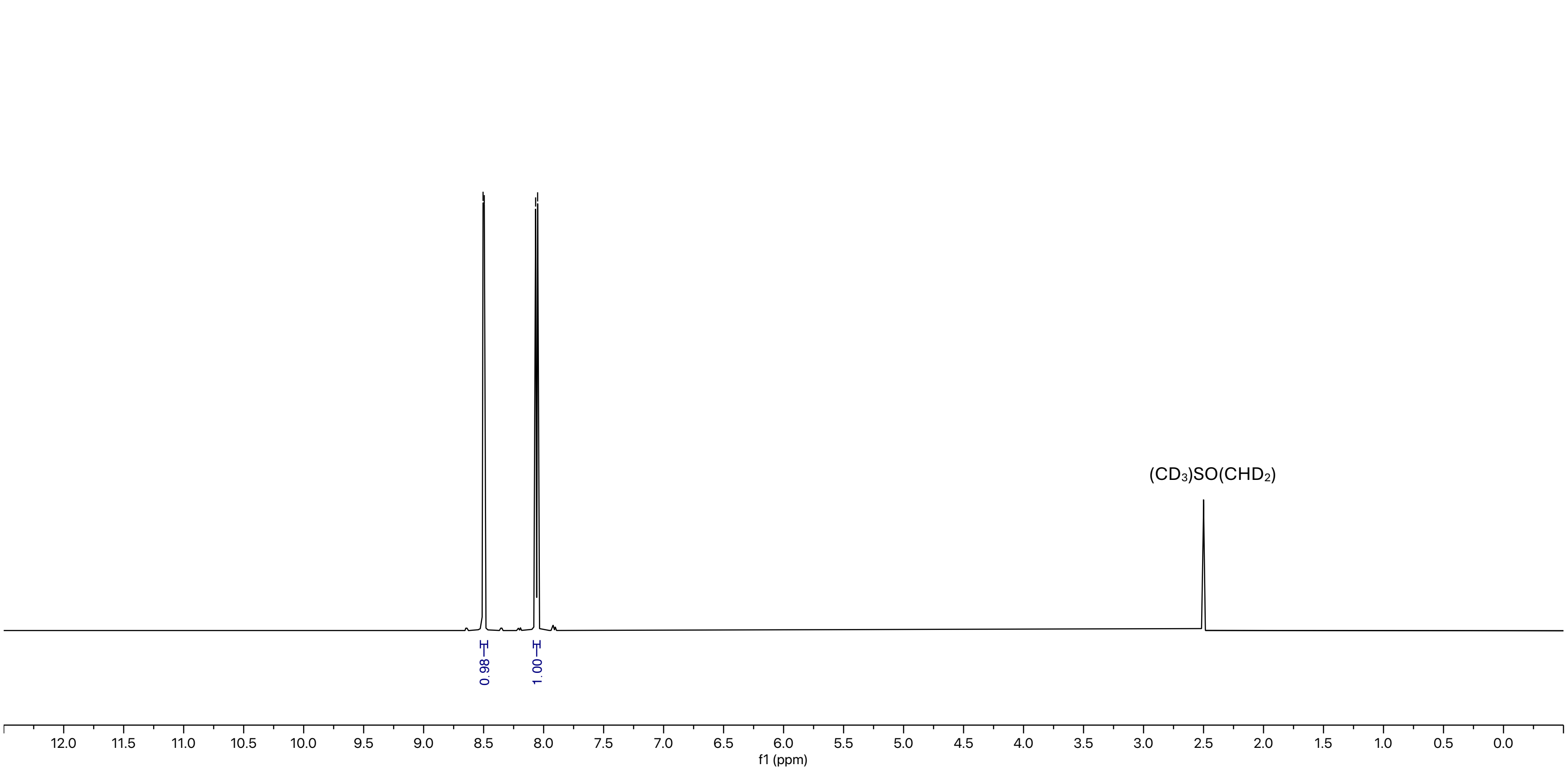

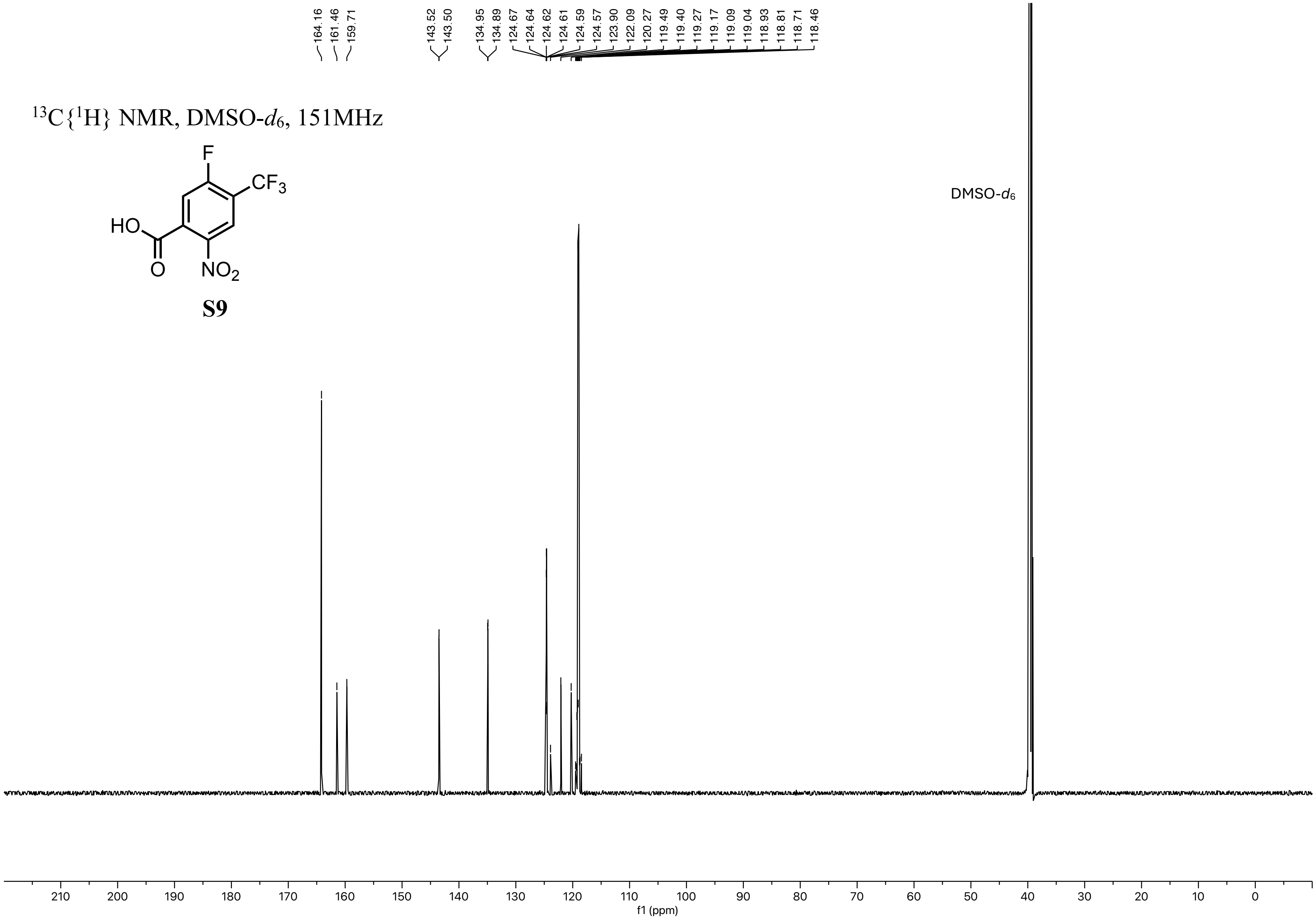

164.16
161.46
159.71
143.52
143.50
134.95
134.89
124.67
124.64
124.62
124.61
124.59
124.57
123.90
122.09
120.27
119.49
119.40
119.27
119.17
119.09
119.04
118.93
118.81
118.71
118.46
$^{13}C\{^1H\}$ NMR, DMSO-$d_6$, 151MHz
F
$CF_3$
HO
O
$NO_2$
S9
DMSO-$d_6$
f1 (ppm)

$^{19}$F NMR, DMSO-$d_6$, 376MHz

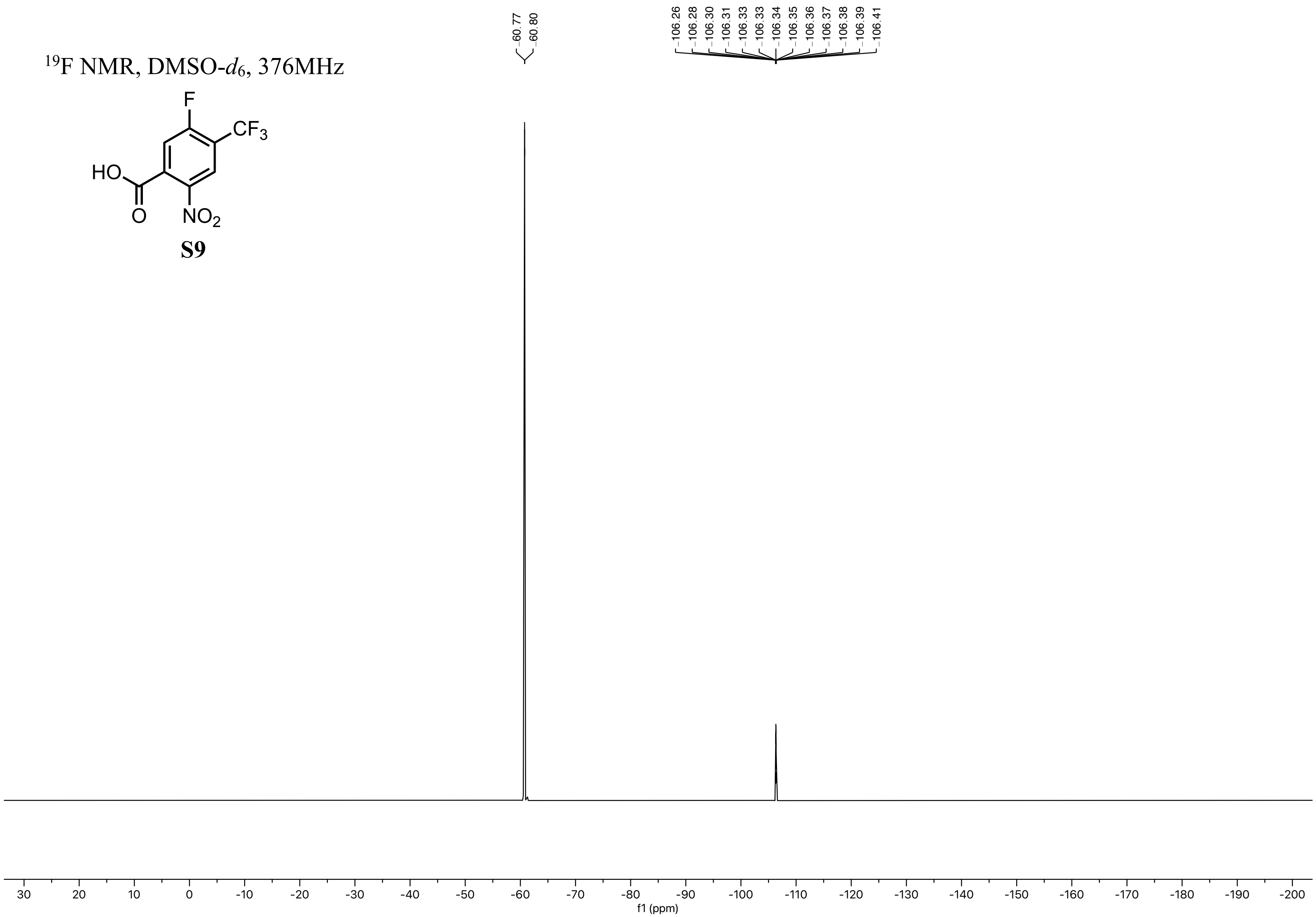

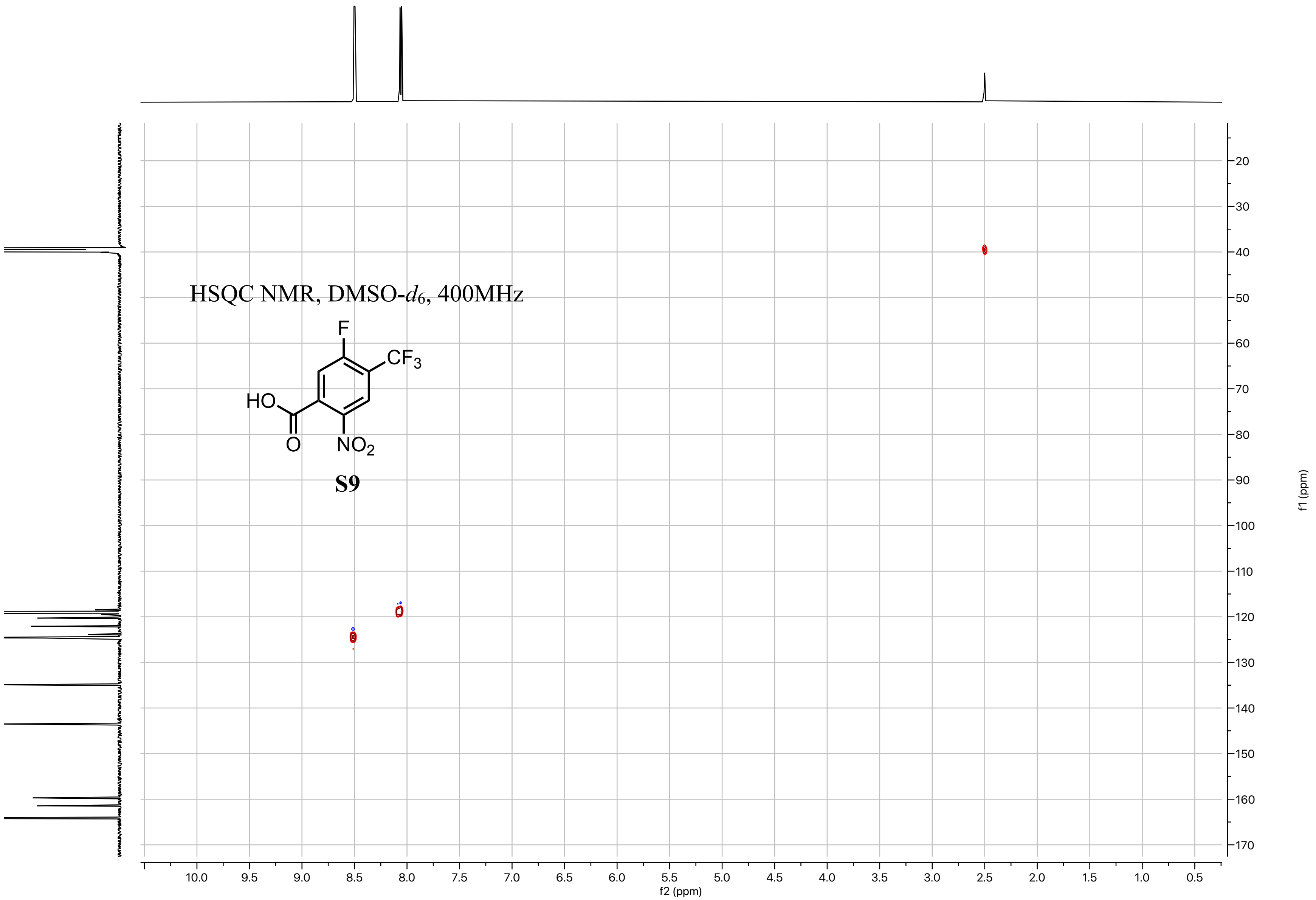
HSQC NMR, DMSO-$d_6$, 400MHz
F
$CF_3$
HO
O
$NO_2$
S9
f2 (ppm)
f1 (ppm)

[1]H NMR, DMSO-$d_6$, 600MHz

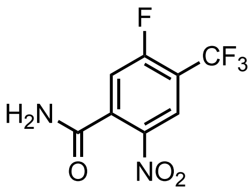


**S10**

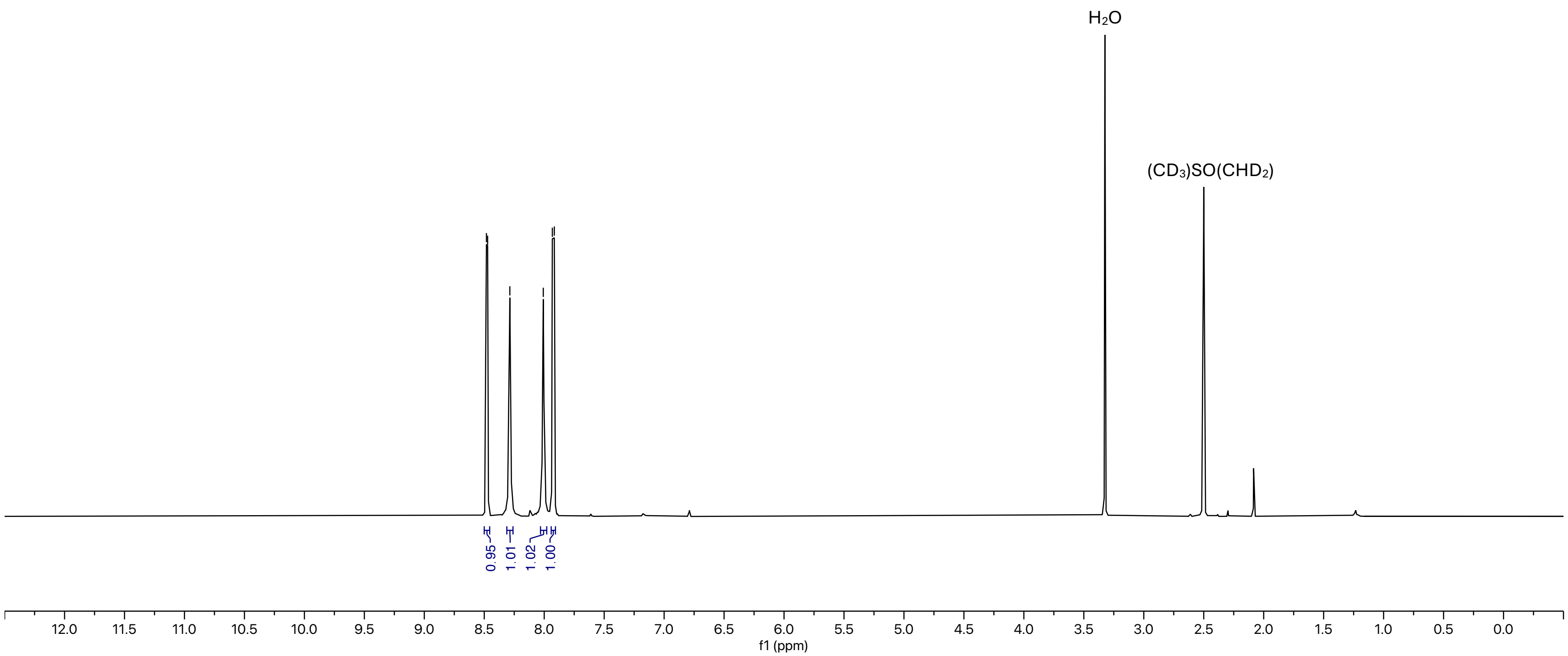

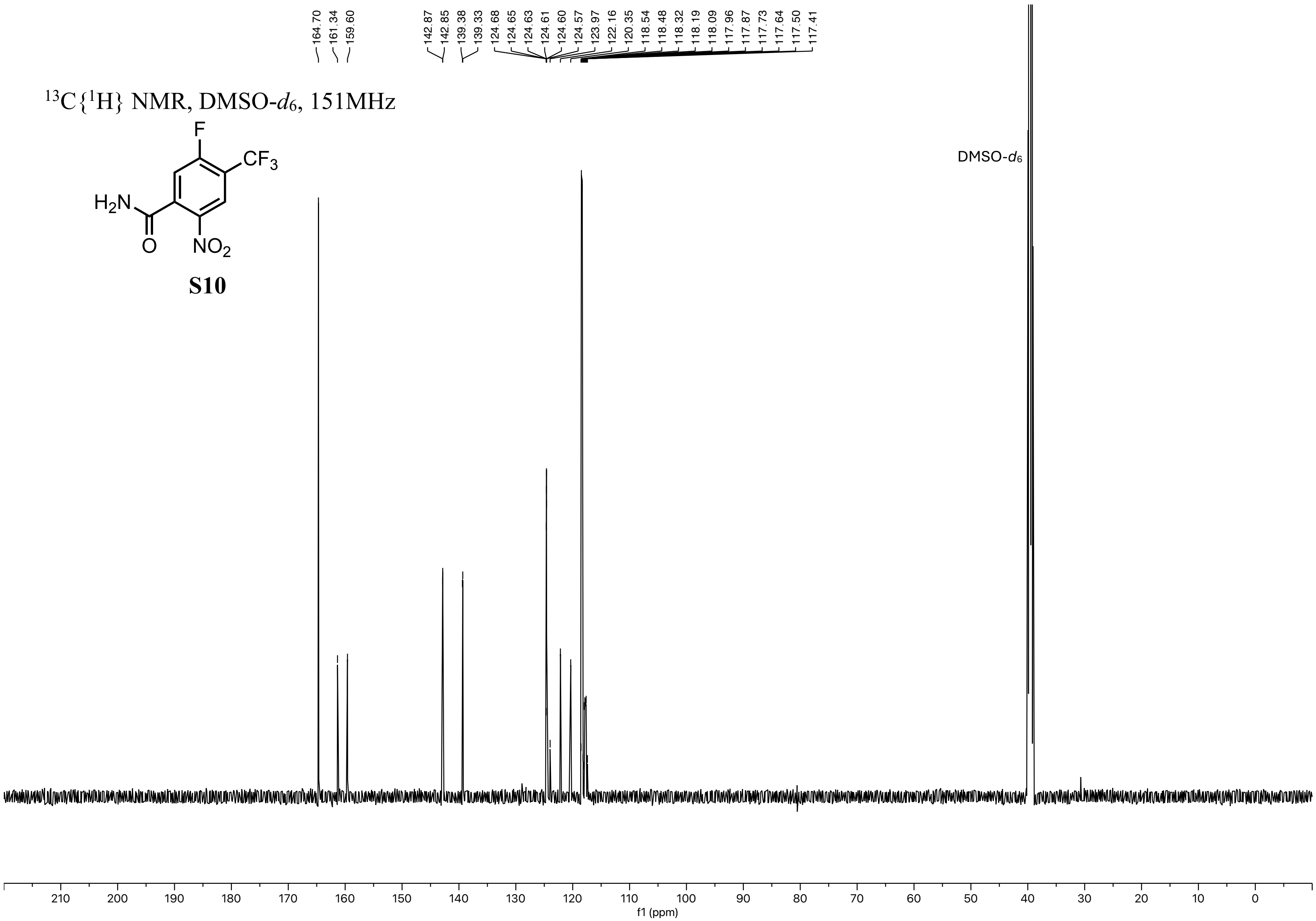
164.70
161.34
159.60
142.87
142.85
139.38
139.33
124.68
124.65
124.63
124.61
124.60
124.57
123.97
122.16
120.35
118.54
118.48
118.32
118.19
118.09
117.96
117.87
117.73
117.64
117.50
117.41
$^{13}C\{^1H\}$ NMR, DMSO-$d_6$, 151MHz
F
$CF_3$
$H_2N$
O
$NO_2$
S10
DMSO-$d_6$
210 200 190 180 170 160 150 140 130 120 110 100 90 80 70 60 50 40 30 20 10 0
f1 (ppm)

$^{19}$F NMR, DMSO-$d_6$, 470MHz

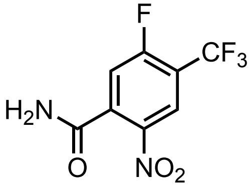

**S10**

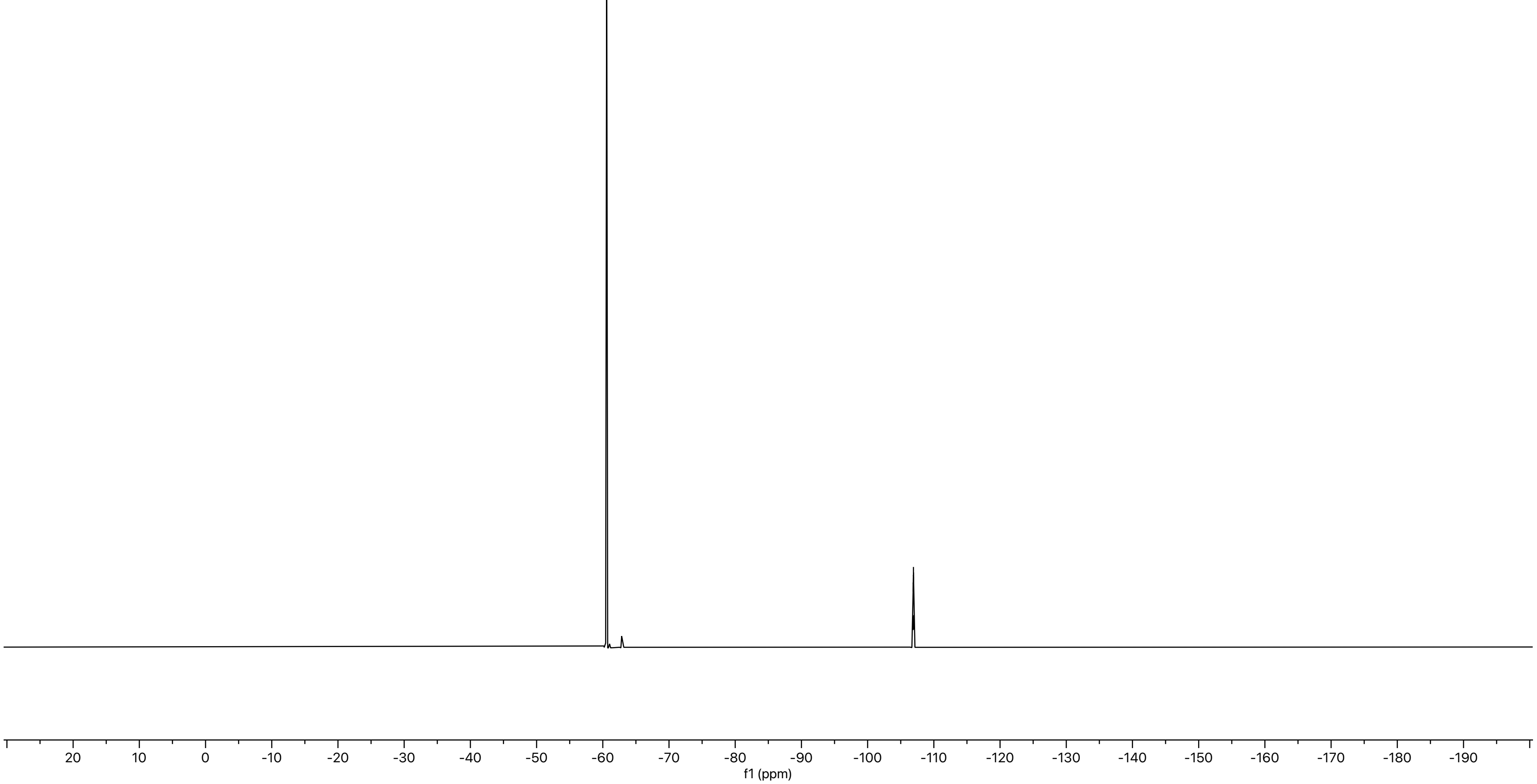

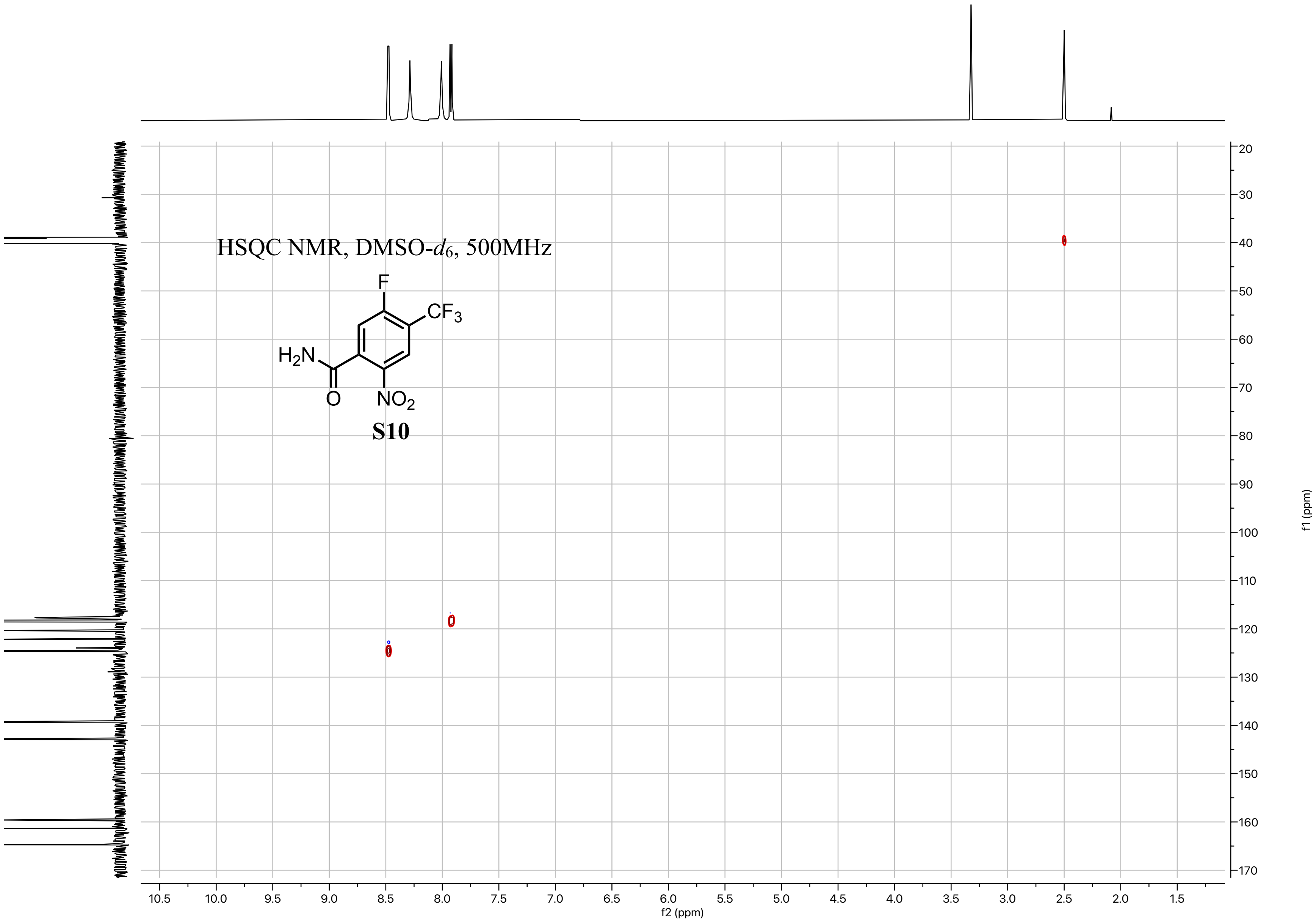
HSQC NMR, DMSO-$d_6$, 500MHz
F
$CF_3$
$H_2N$
O
$NO_2$
S10
f2 (ppm)
f1 (ppm)

$^{1}H$ NMR, DMSO-$d_6$, 600MHz

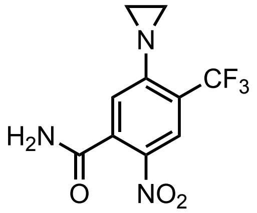

**7**

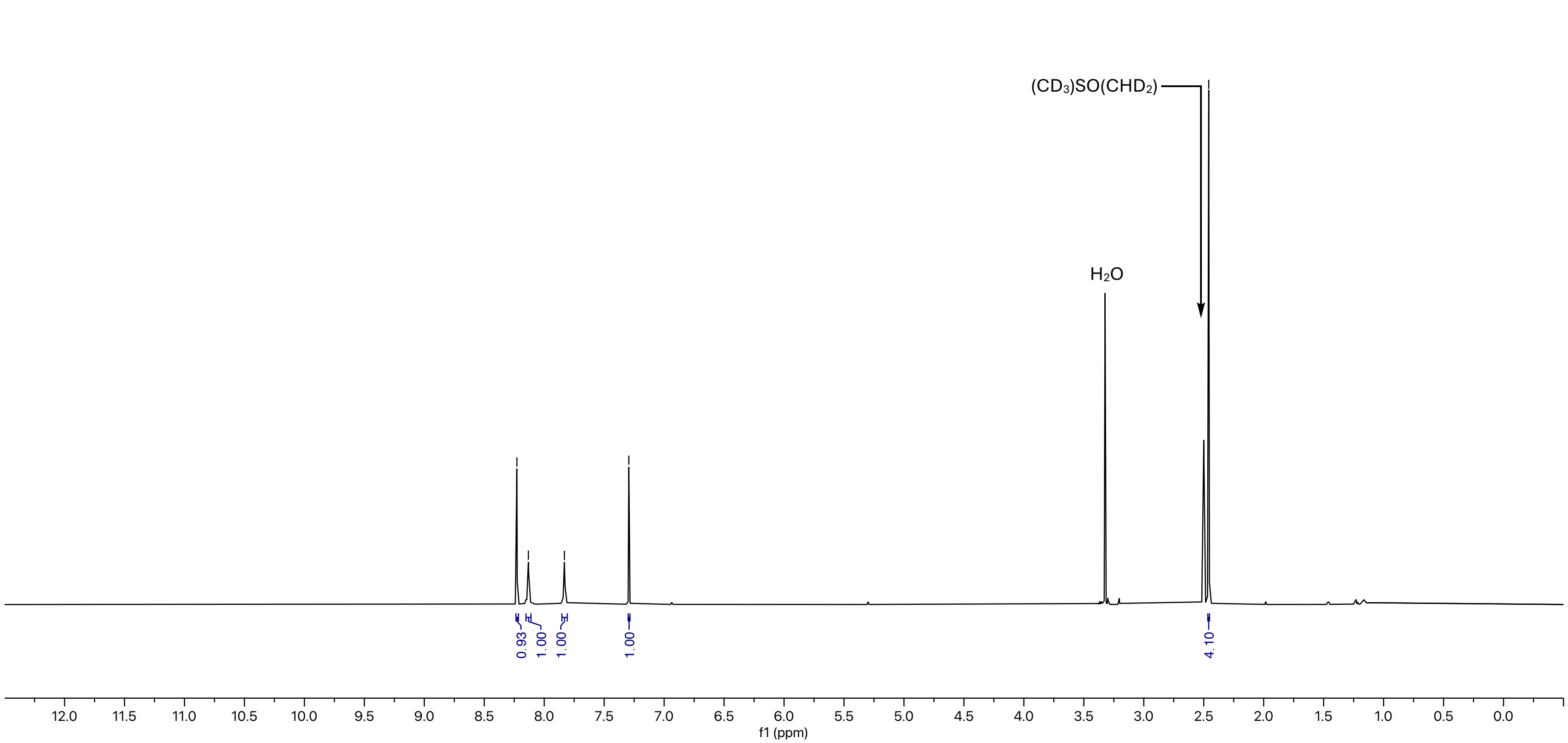

$^{13}C\{^{1}H\}$ NMR, DMSO-$d_6$, 151MHz

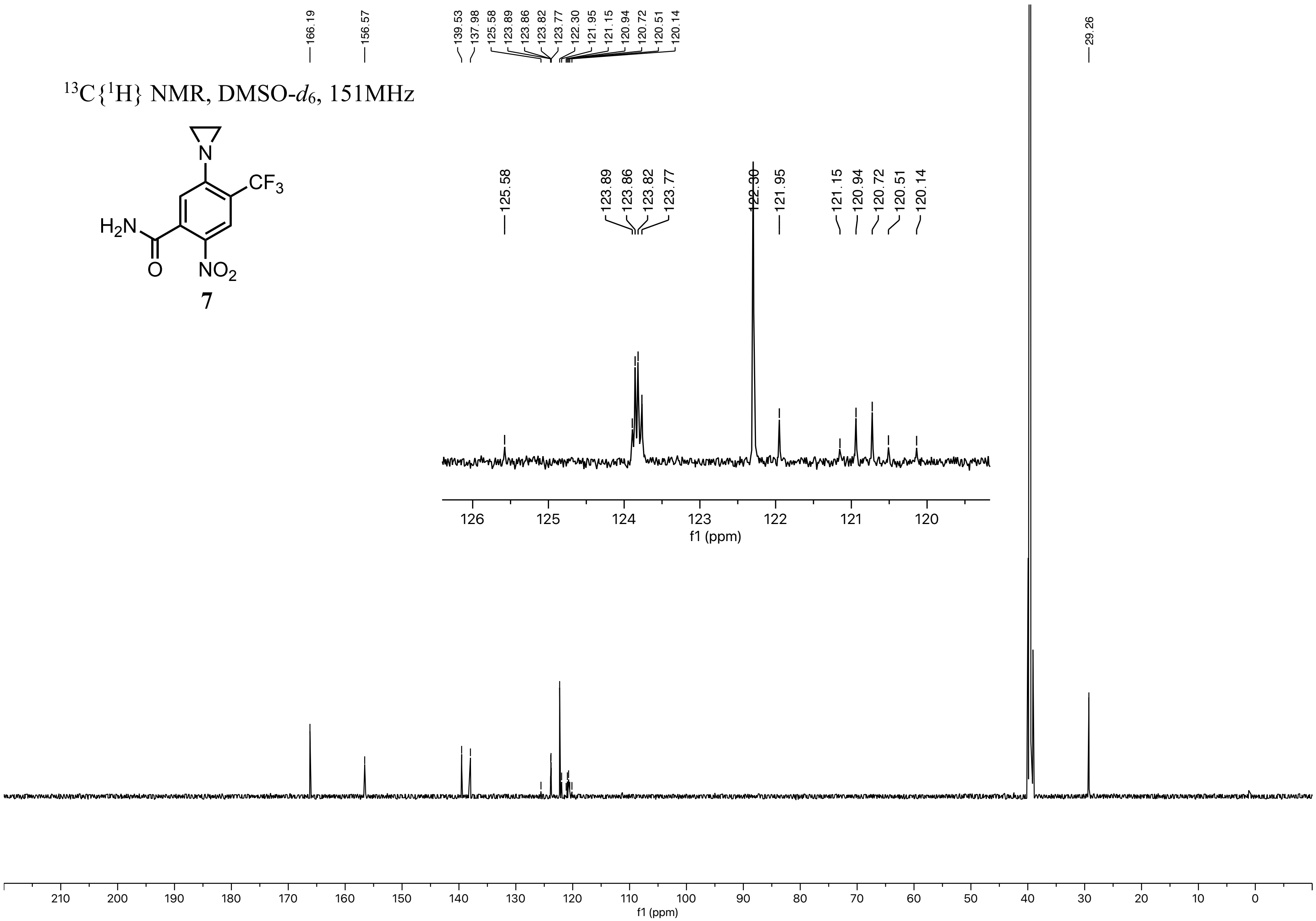


**7**

$^{19}F$ NMR, DMSO-$d_6$, 376MHz

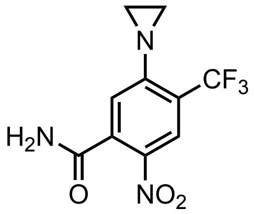


**7**

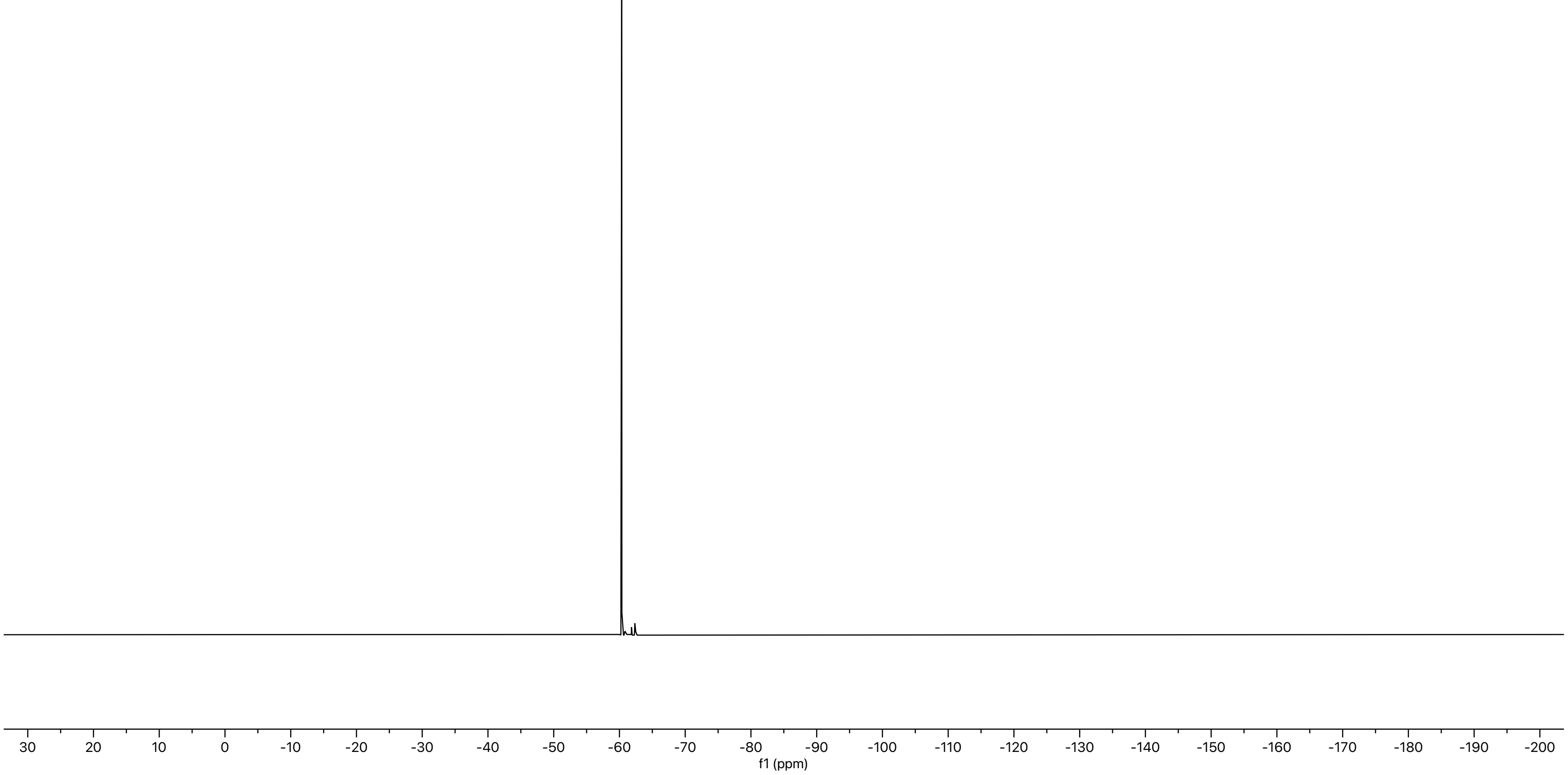

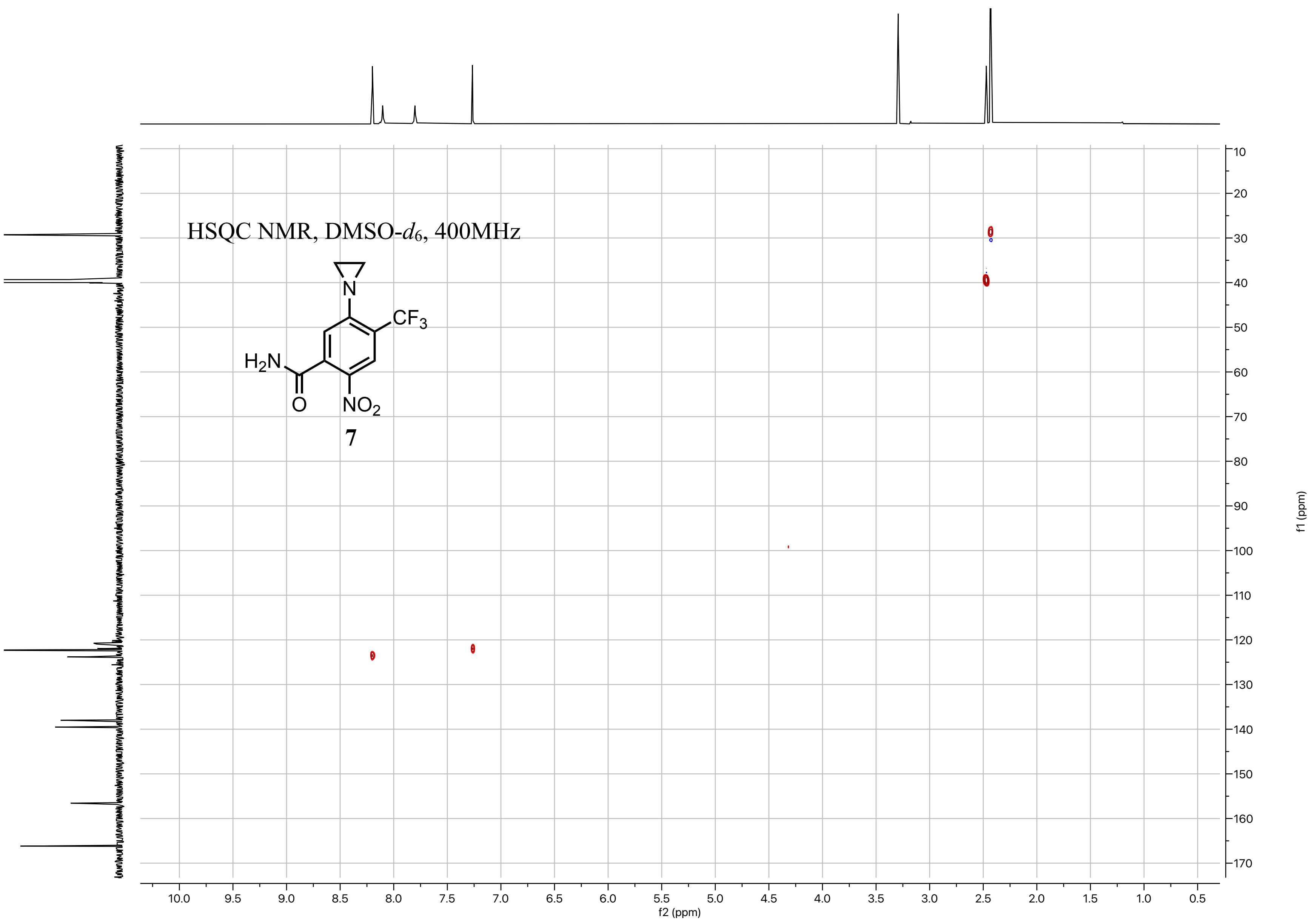
HSQC NMR, DMSO-$d_6$, 400MHz
N
$CF_3$
$H_2N$
O
$NO_2$
7
10.0 9.5 9.0 8.5 8.0 7.5 7.0 6.5 6.0 5.5 5.0 4.5 4.0 3.5 3.0 2.5 2.0 1.5 1.0 0.5
f2 (ppm)
10 20 30 40 50 60 70 80 90 100 110 120 130 140 150 160 170
f1 (ppm)

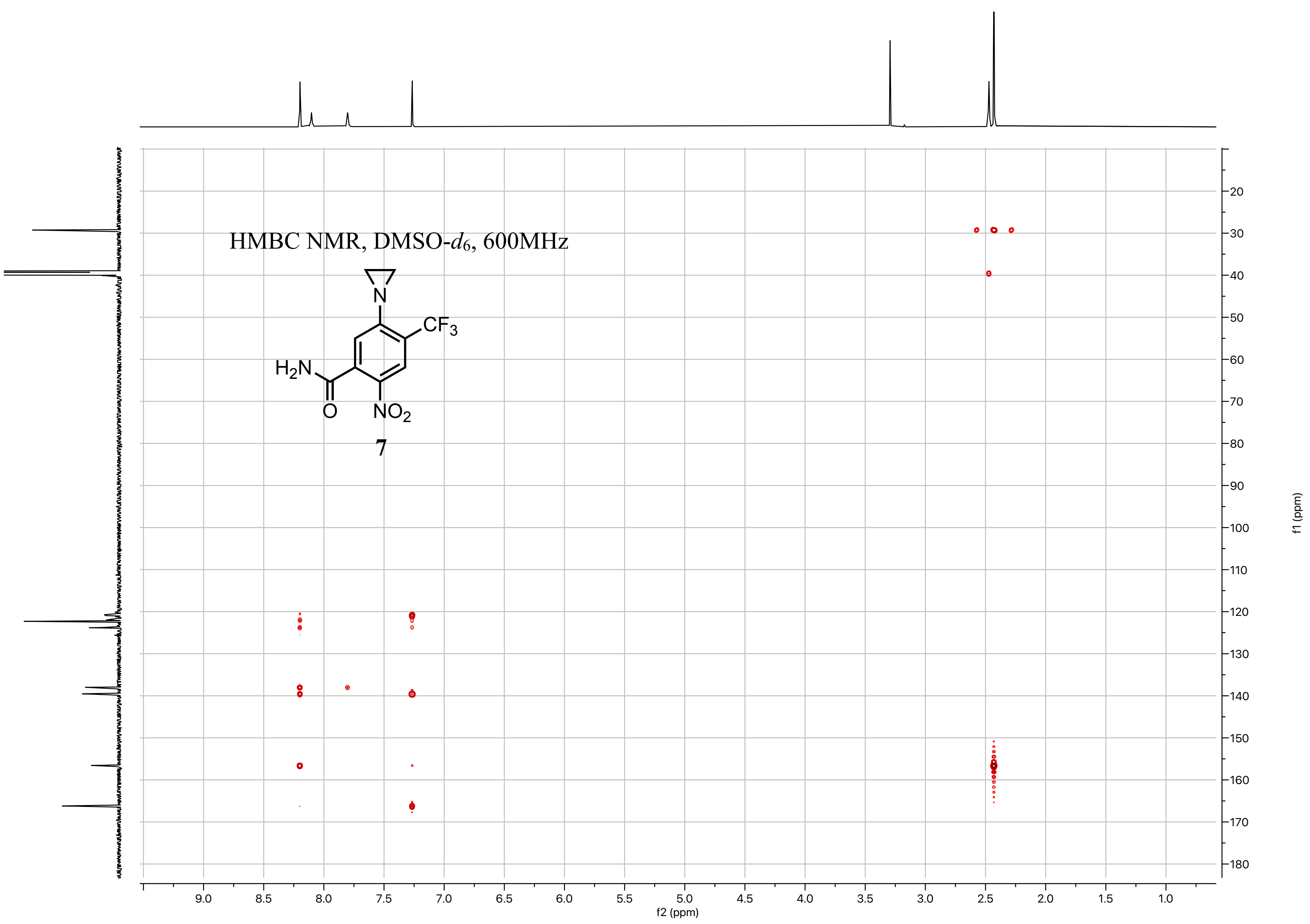
HMBC NMR, DMSO-$d_6$, 600MHz
N
$CF_3$
$H_2N$
O
$NO_2$
7
f2 (ppm)
f1 (ppm)

$^1H$ NMR, DMSO-$d_6$, 400MHz

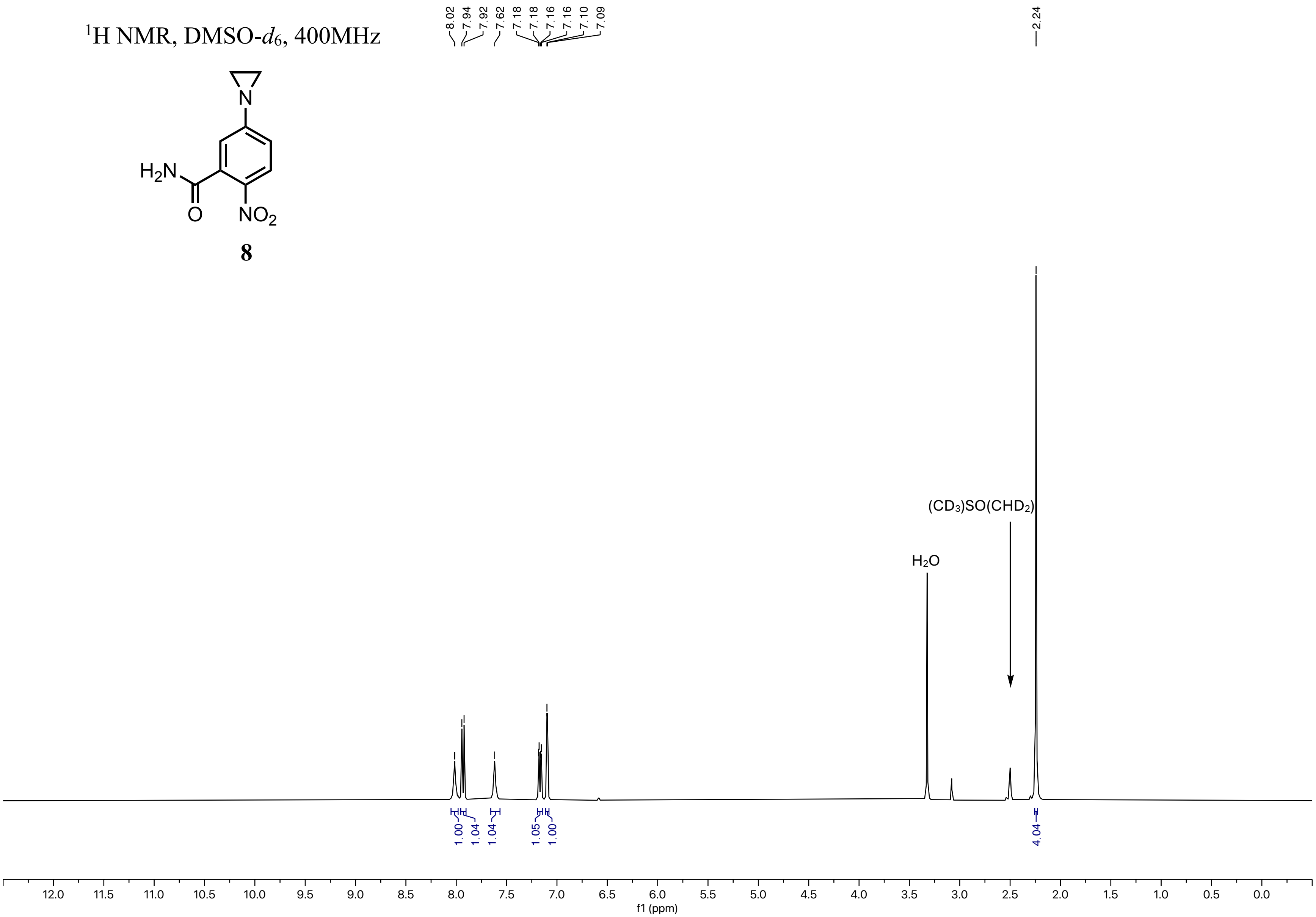

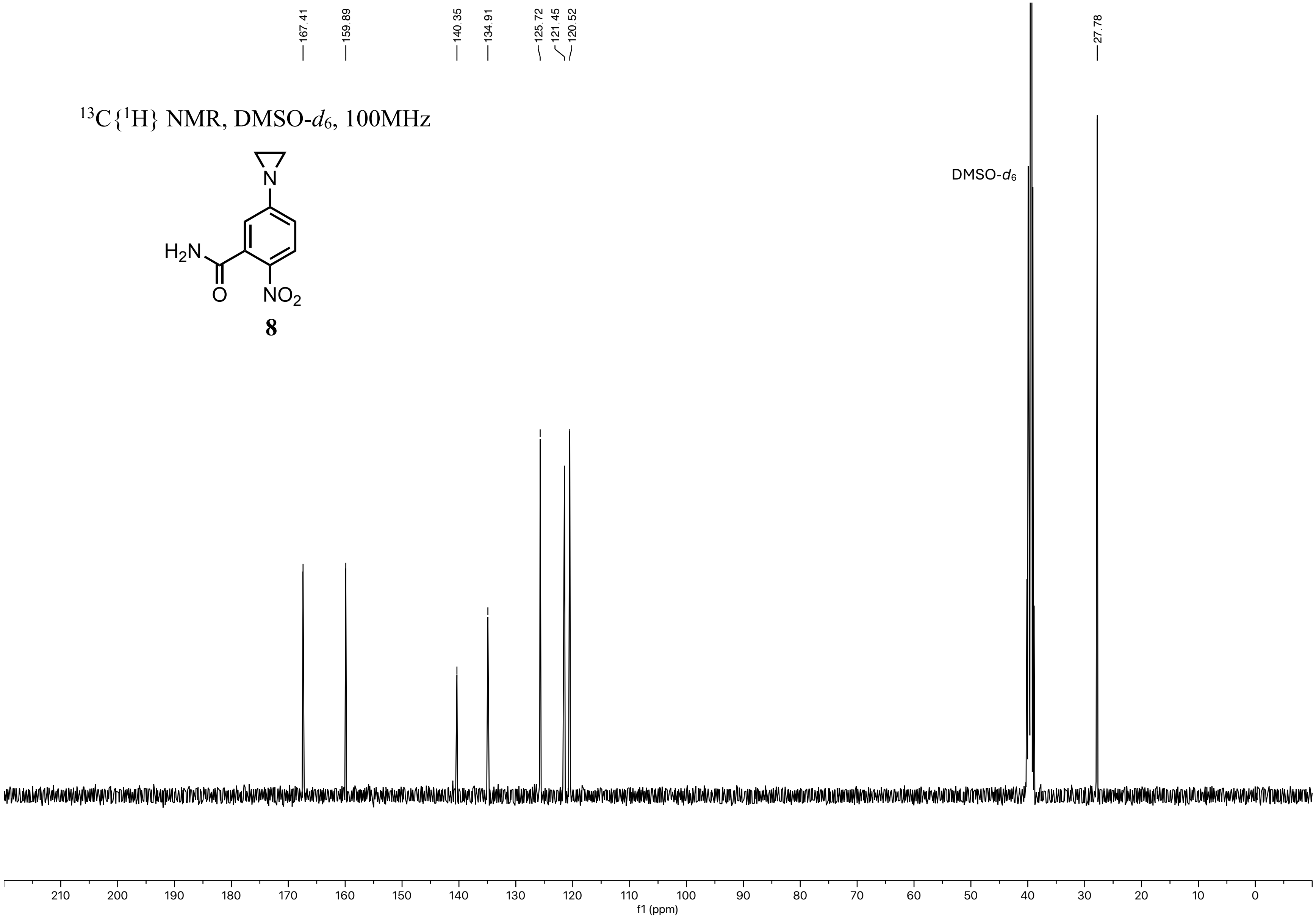
167.41
159.89
140.35
134.91
125.72
121.45
120.52
27.78
$^{13}C\{^{1}H\}$ NMR, DMSO-$d_6$, 100MHz
N
$H_2N$
O
$NO_2$
8
DMSO-$d_6$
210 200 190 180 170 160 150 140 130 120 110 100 90 80 70 60 50 40 30 20 10 0
f1 (ppm)

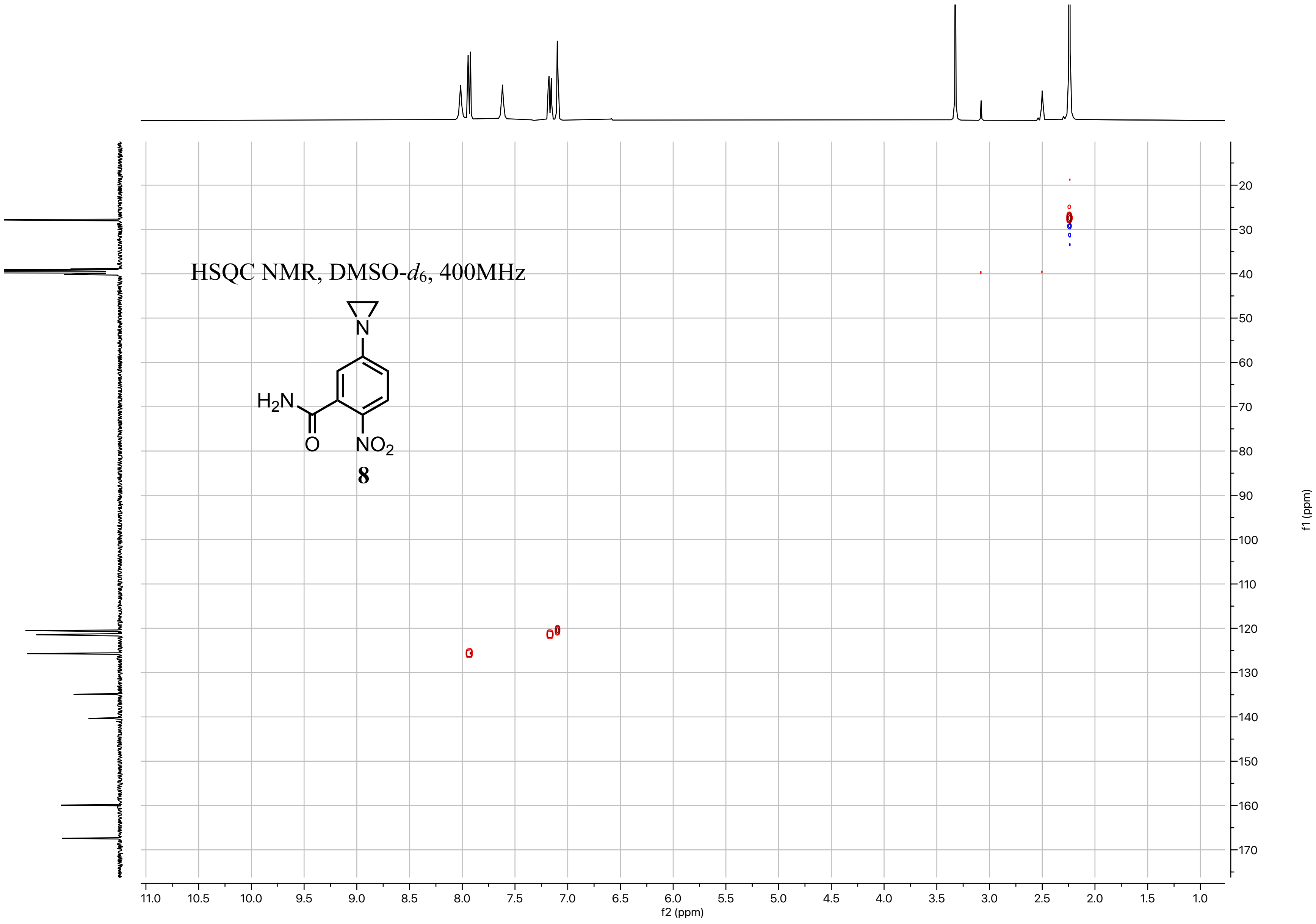
HSQC NMR, DMSO-$d_6$, 400MHz
$H_2N$
O
N
$NO_2$
8
f2 (ppm)
f1 (ppm)
11.0
10.5
10.0
9.5
9.0
8.5
8.0
7.5
7.0
6.5
6.0
5.5
5.0
4.5
4.0
3.5
3.0
2.5
2.0
1.5
1.0
20
30
40
50
60
70
80
90
100
110
120
130
140
150
160
170

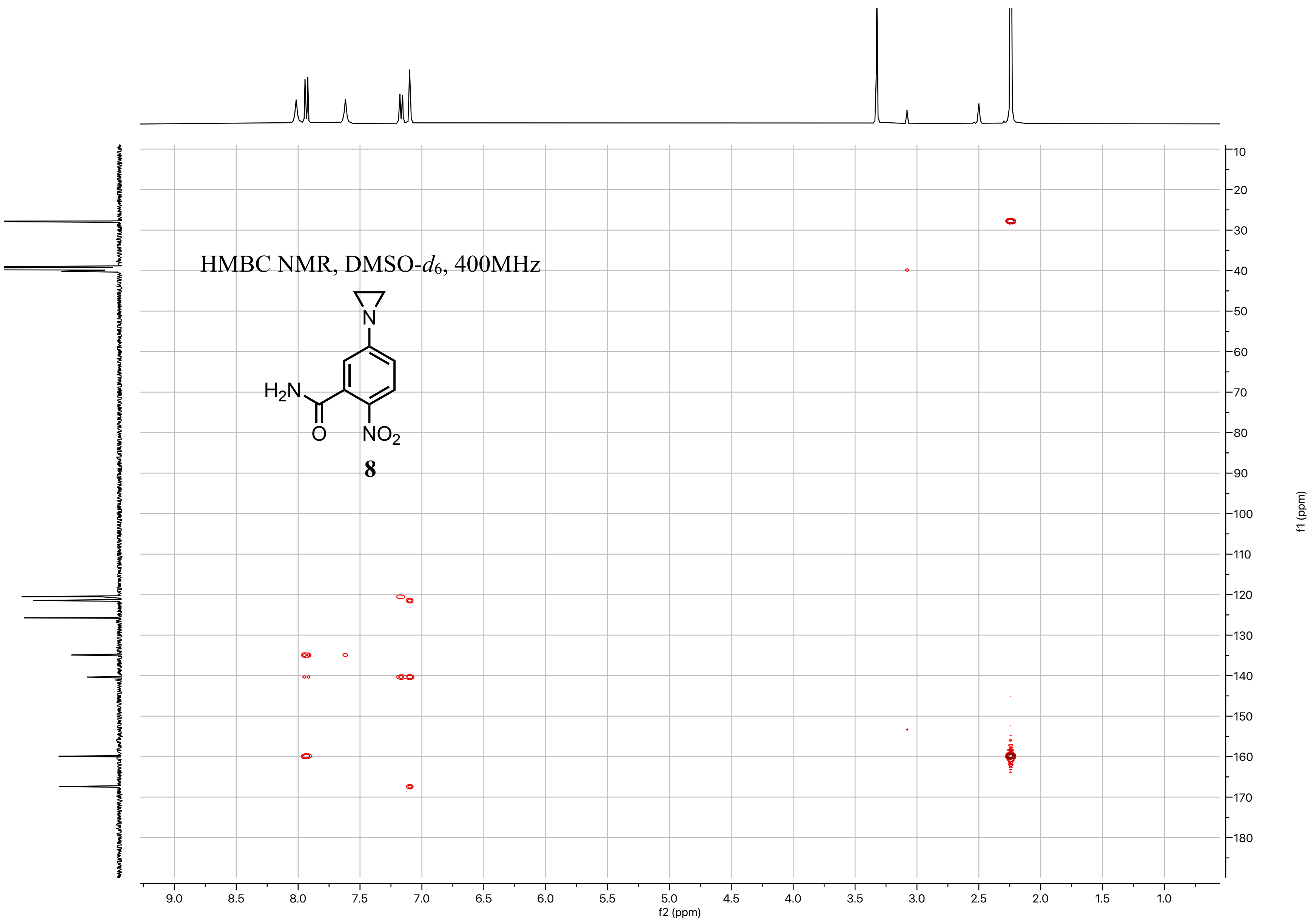
HMBC NMR, DMSO-d6, 400MHz
H2N
N
O
NO2
8
f1 (ppm)
f2 (ppm)

$^{1}H$ NMR, DMSO-$d_6$, 600MHz

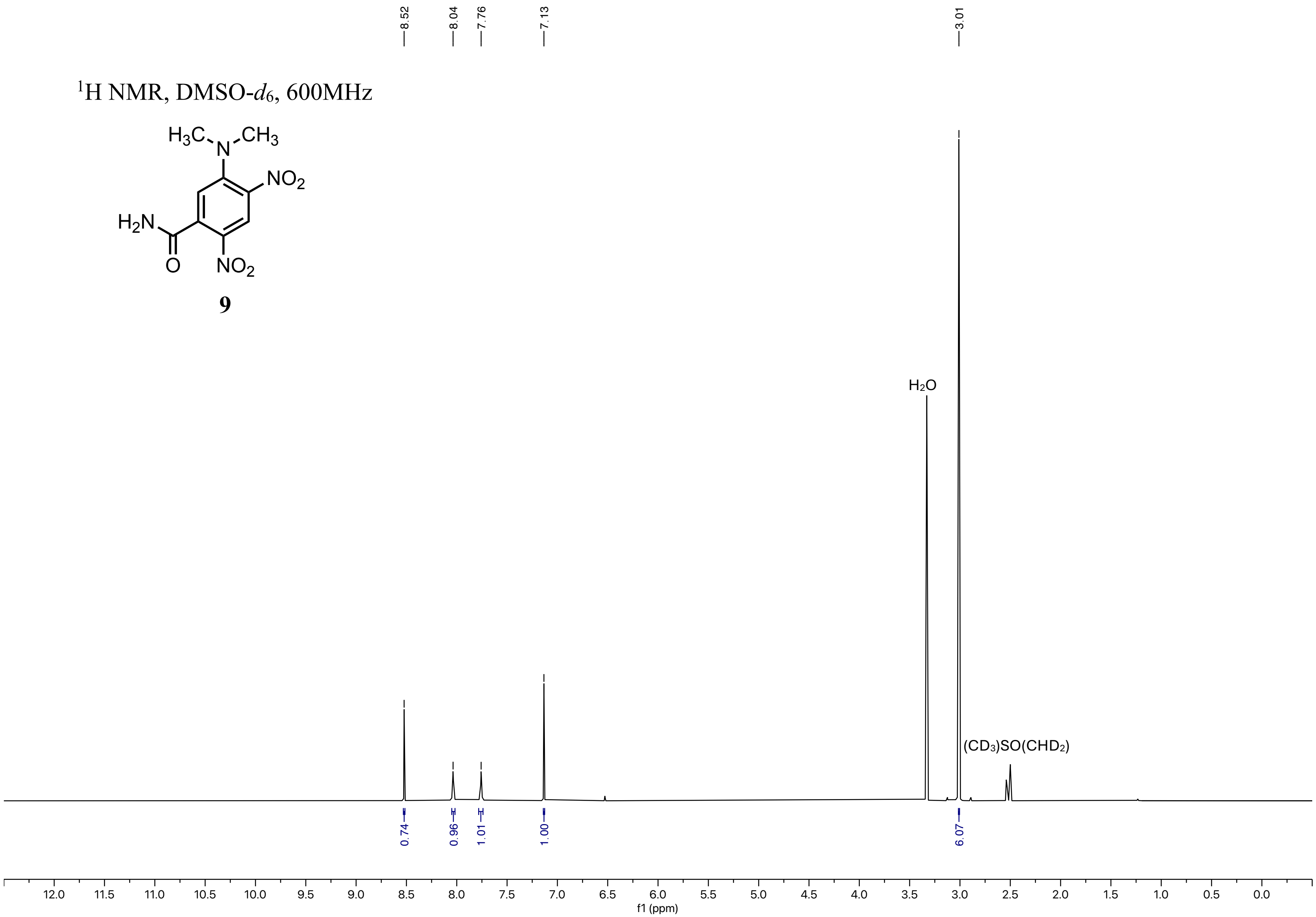


**9**

$^{13}C\{^{1}H\}$ NMR, DMSO-$d_6$, 151MHz

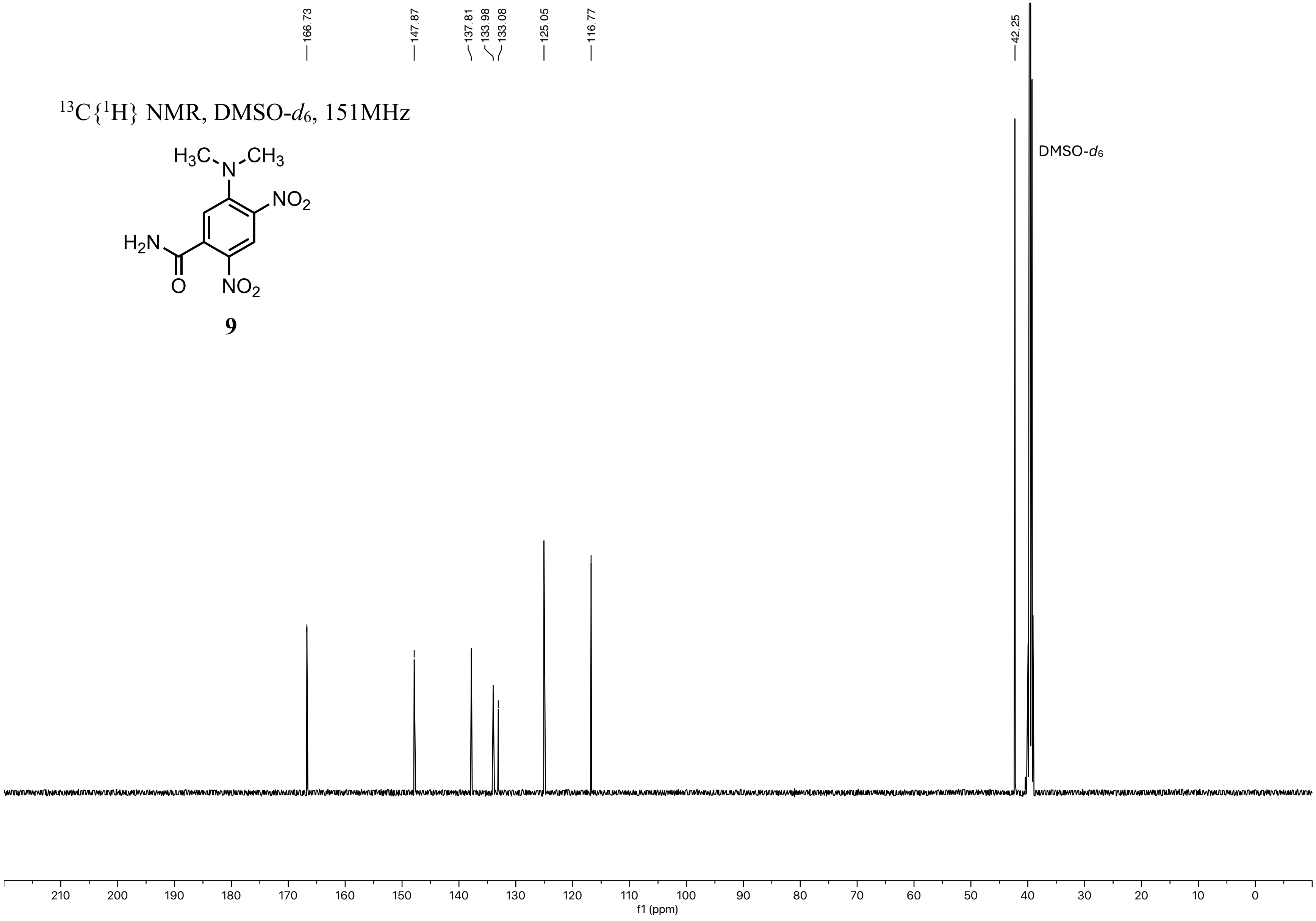

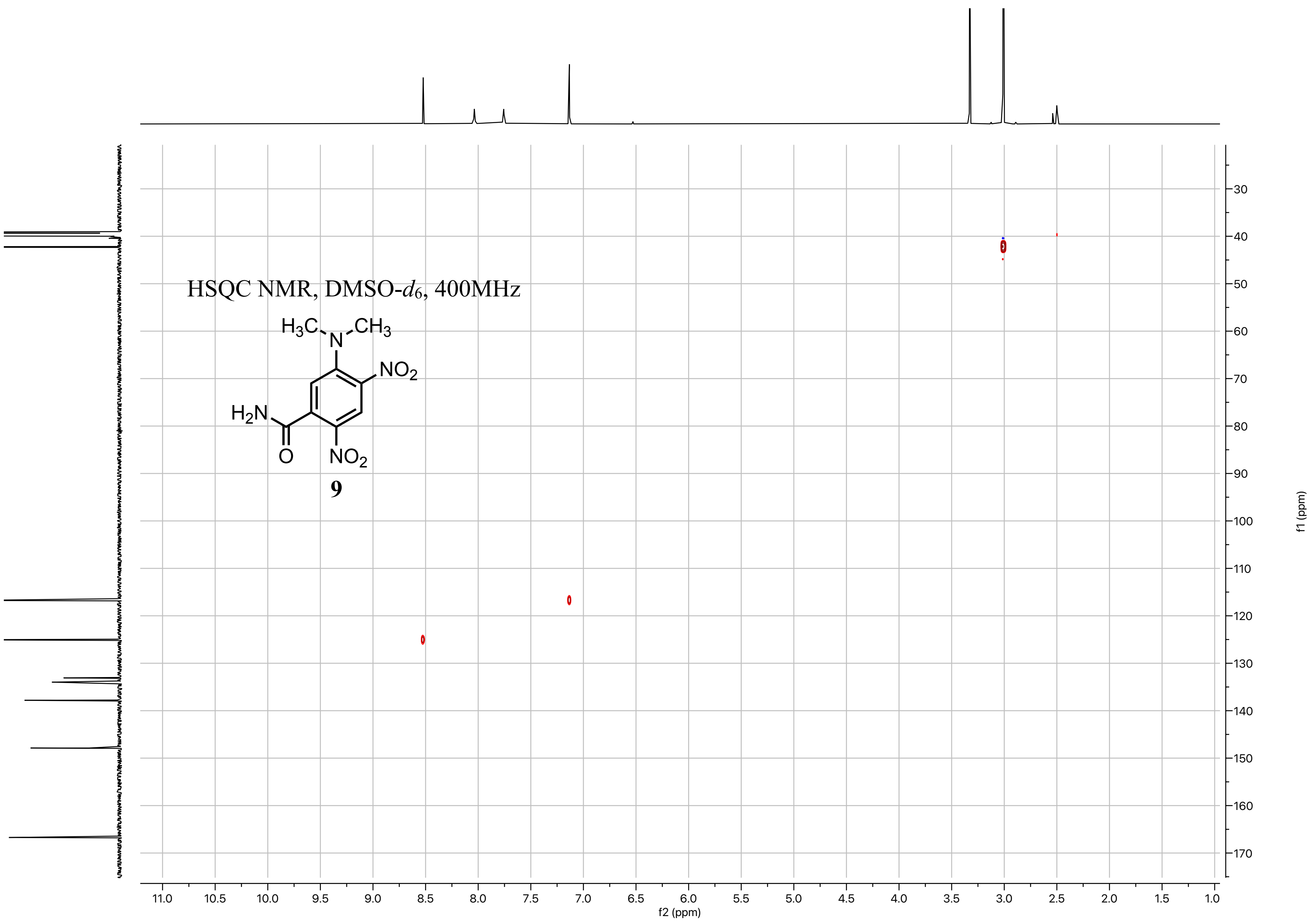
HSQC NMR, DMSO-$d_6$, 400MHz
$H_3C$
N
$CH_3$
$NO_2$
$H_2N$
O
$NO_2$
9
f2 (ppm)
f1 (ppm)

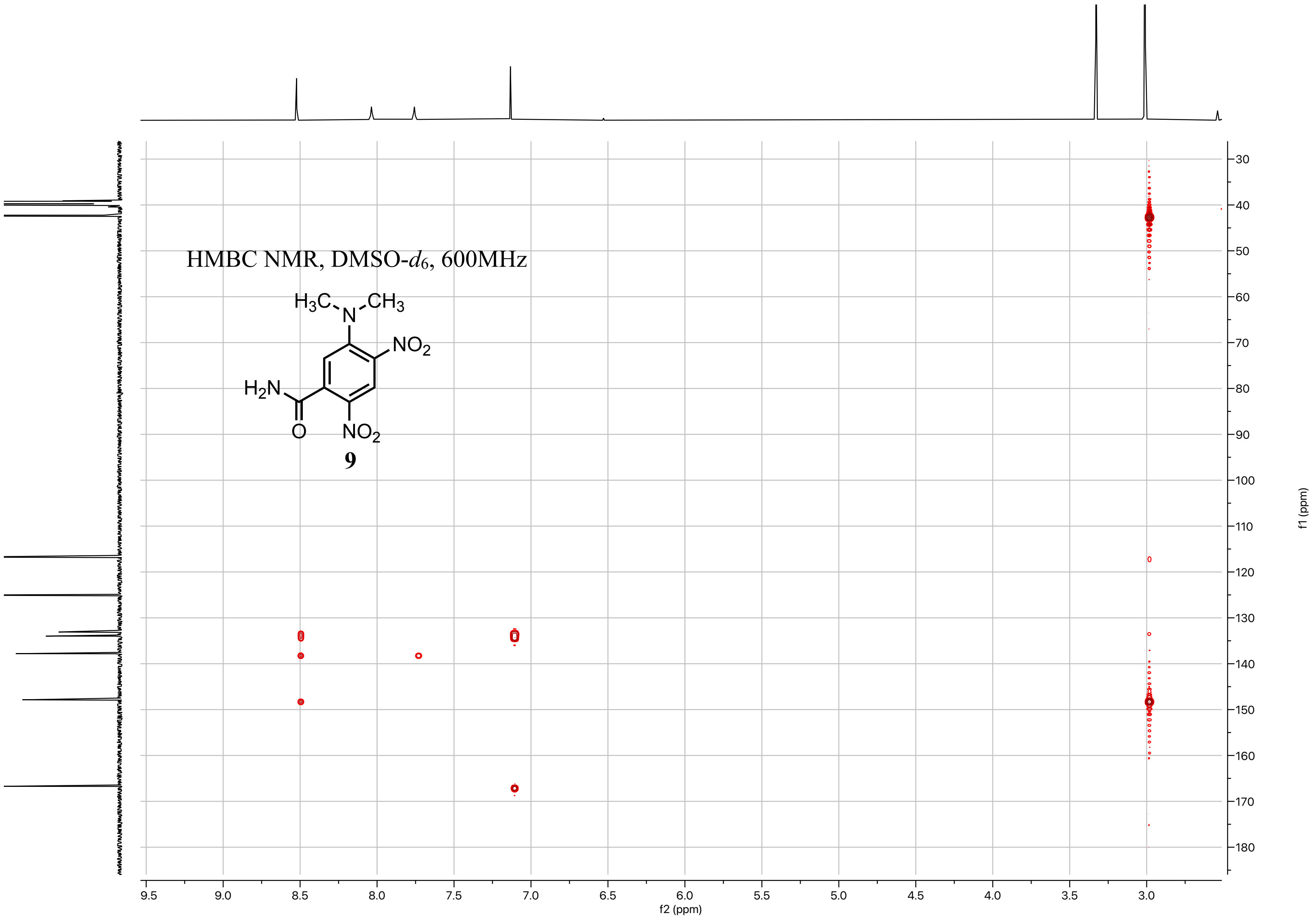
HMBC NMR, DMSO-$d_6$, 600MHz
$H_3C$ N $CH_3$
$NO_2$
$H_2N$
O
$NO_2$
9
f2 (ppm)
f1 (ppm)

$^{1}H$ NMR, DMSO-$d_6$, 600MHz

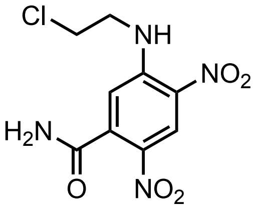

**10**

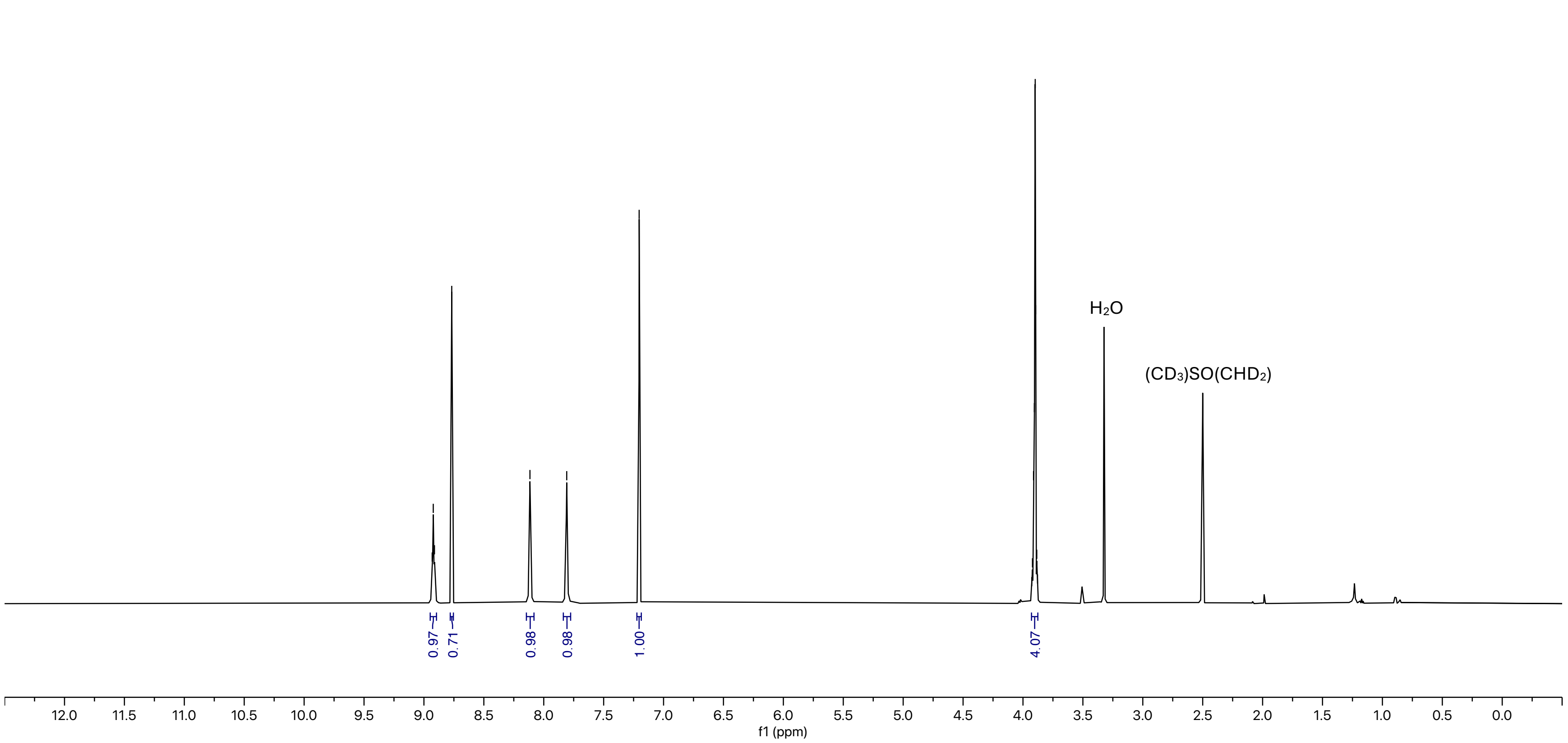

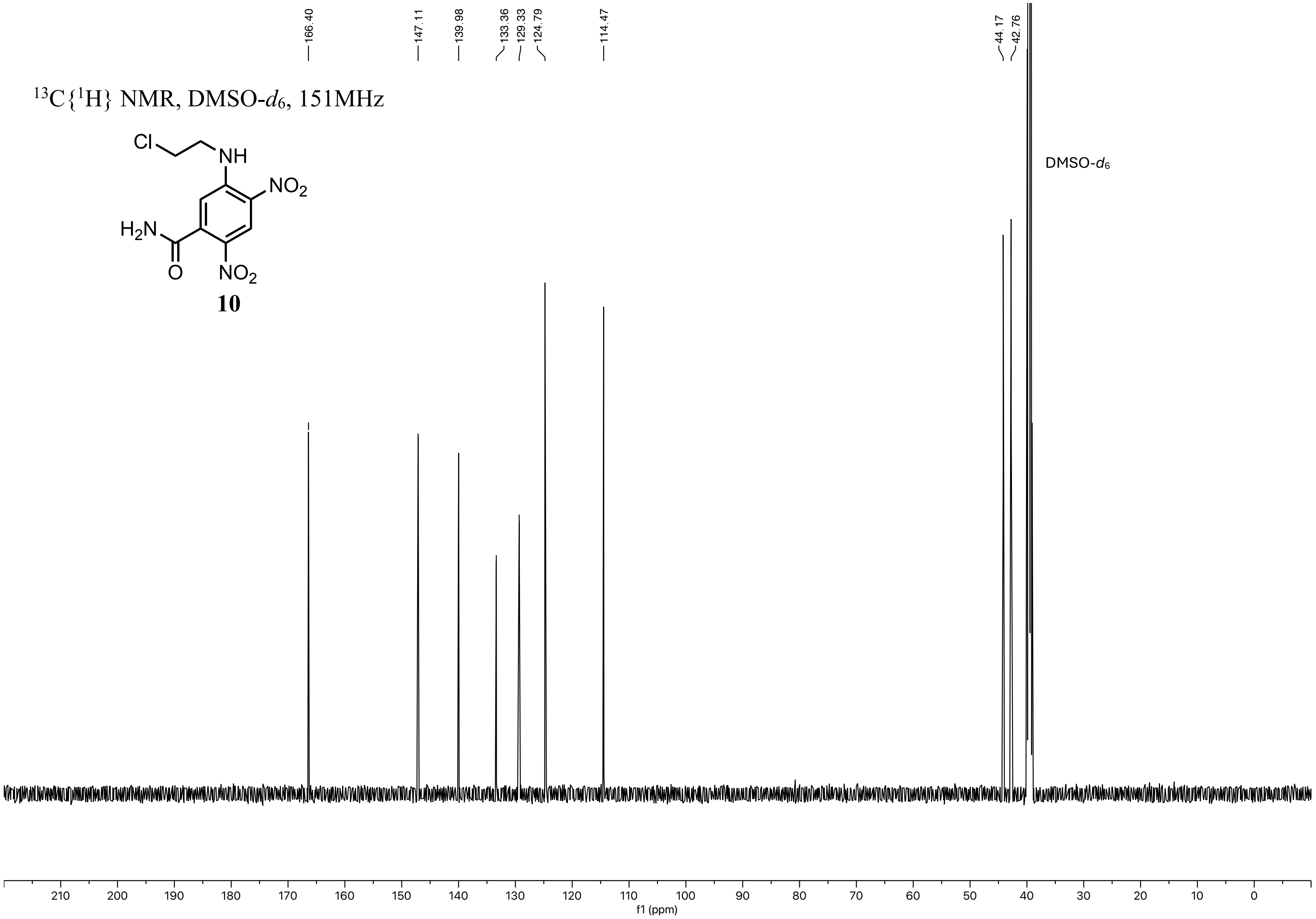

166.40
147.11
139.98
133.36
129.33
124.79
114.47
44.17
42.76
13C{1H} NMR, DMSO-d6, 151MHz
Cl
NH
NO2
H2N
O
NO2
10
DMSO-d6
210 200 190 180 170 160 150 140 130 120 110 100 90 80 70 60 50 40 30 20 10 0
f1 (ppm)

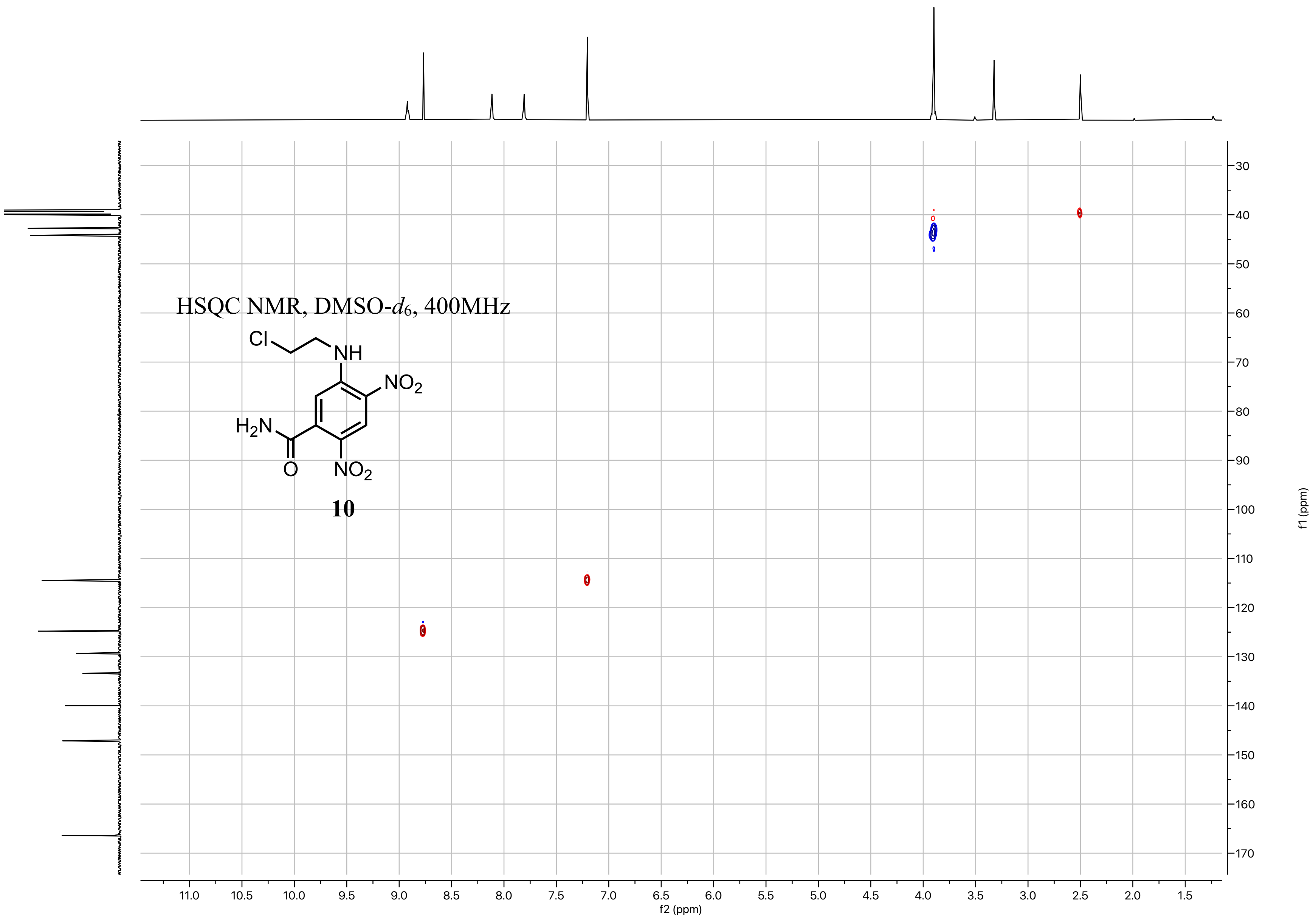
HSQC NMR, DMSO-$d_6$, 400MHz
Cl
NH
$NO_2$
$H_2N$
O
$NO_2$
10
11.0
10.5
10.0
9.5
9.0
8.5
8.0
7.5
7.0
6.5
6.0
5.5
5.0
4.5
4.0
3.5
3.0
2.5
2.0
1.5
f2 (ppm)
30
40
50
60
70
80
90
100
110
120
130
140
150
160
170
f1 (ppm)

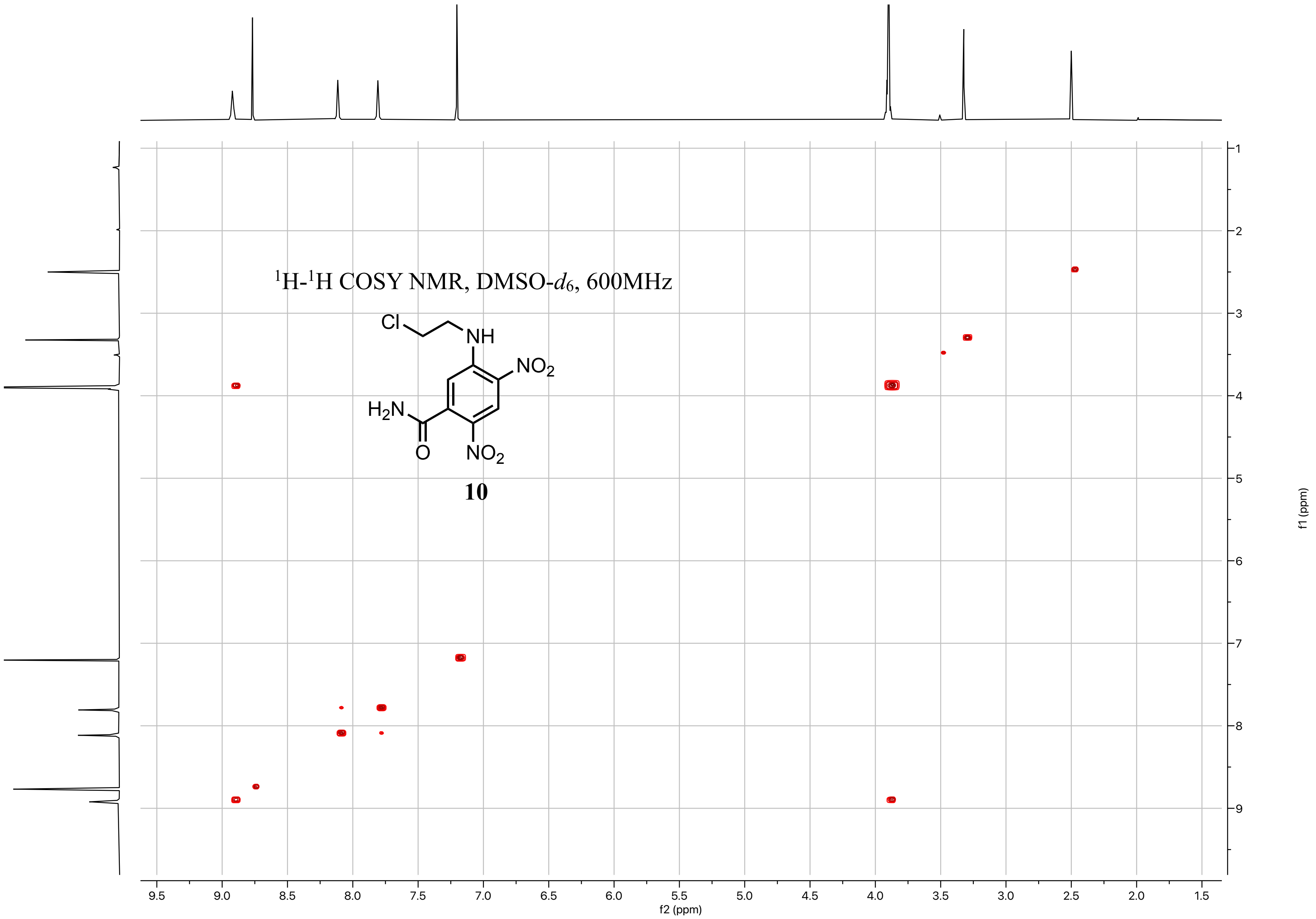

¹H-¹H COSY NMR, DMSO-d6, 600MHz
Cl
NH
NO2
H2N
O
NO2
10
9.5 9.0 8.5 8.0 7.5 7.0 6.5 6.0 5.5 5.0 4.5 4.0 3.5 3.0 2.5 2.0 1.5
f2 (ppm)
1 2 3 4 5 6 7 8 9
f1 (ppm)

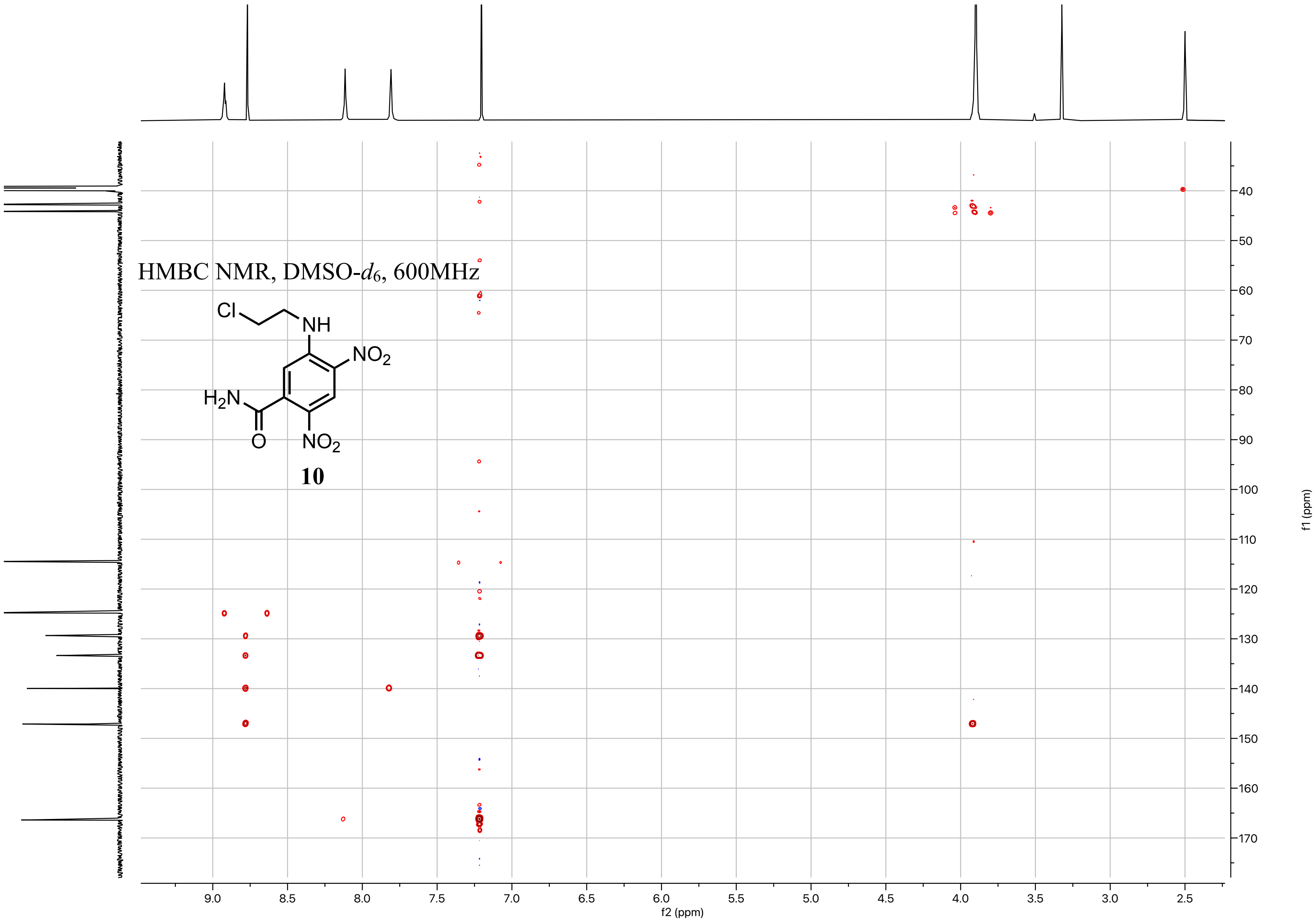
HMBC NMR, DMSO-$d_6$, 600MHz
Cl
NH
$NO_2$
$H_2N$
O
$NO_2$
10
f2 (ppm)
f1 (ppm)

**References.**
